# Learning a shared vocabulary between episodic and semantic memory enhances recall and compositional consolidation

**DOI:** 10.1101/2025.10.03.680209

**Authors:** Albert Albesa-González, Claudia Clopath

## Abstract

Semantic knowledge is thought to emerge through the consolidation of episodic experience, yet the biological mechanisms by which reusable semantic representations are extracted from complex episodes remain unclear. Conversely, semantic representations can themselves be found within the medial temporal lobe, raising the questions of how they arise there and why structured semantic overlap should benefit an episodic memory system. We propose that replay establishes a shared semantic vocabulary between MTL and CTX through two complementary processes. First, episodic replay enables neocortex to extract reusable semantic concepts from overlapping episodes via a form of dictionary learning. Second, spontaneous cortical activity replays these learned concepts back to MTL, allowing semantic representations to become part of the episodic code. We implement this framework in a circuit model using local Hebbian plasticity and homeostatic competition. The model reproduces key experimental observations, including concept-cell-like representations, increased overlap between related memories following consolidation, and synaptic reorganization associated with systems consolidation. Functionally, embedding semantic representations within episodic memory improves recall by providing an error-correcting scaffold and enhancing replay consistency, inducing a generalization bias in the consolidation of compositional representations. This further explains experimental effects hard to reconcile with standard accounts of consolidation, such as the impact of disrupting semantics in episodic recall and semantic deficits following medial temporal lobe lesions. More broadly, our results suggest that consolidation progressively builds a shared vocabulary between episodic and semantic memory, allowing structure extracted from past experience to organize the representation and consolidation of future memories.

## Introduction

The distinction between episodic and semantic memory has been central to cognitive neuroscience since its first formalization (Tulving, 1972). In its original view, episodic memory captures individual experiences, while semantic memory accumulates generalized knowledge that emerges from repeated exposure to similar episodes (Squire, 2004). One influential computational explanation for this division is provided by the Complementary Learning Systems (CLS) framework. This theory posits that a fast-learning system (the hippocampus and the rest of the medial temporal lobe, MTL) specializes in rapidly encoding episodic memories. In contrast, a slow-learning neocortical (CTX) system works in concert with the MTL to consolidate and generalize memories (McClelland et al., 1995; O’Reilly et al., 2014).

One of the central assumptions of consolidation theories is that replay events originating in the hippocampus provide a teaching signal to neocortex. This idea has been remarkably successful in explaining how semantic knowledge can gradually emerge from episodic experience and remains a cornerstone of Complementary Learning Systems theory. However, the precise computational mechanism underlying replay-driven learning remains unclear. On the one hand, several influential models are strongly inspired by the machine learning paradigm (Saxe et al., 2019; Singh et al., 2022; W. Sun et al., 2023; Spens & Burgess, 2024b). As a result, they often rely on a predefined distinction between input, hidden, and output representations as the backbone of cortical learning. How such distinctions could be stored within replay events and translated into appropriate plasticity signals remains unknown. Furthermore, these approaches typically assume that hippocampal replay faithfully reinstates the entirety of episodic neural activity during encoding, an assumption that is difficult to reconcile with evidence that replay is sparse, reconstructive, and only partially reinstates previous activity (Swanson et al., 2020; Stoianov et al., 2022). Finally, learning commonly relies on error-driven or backpropagation-like mechanisms, whose biological implementation remains debated (Lillicrap et al., 2020). On the other hand, more biologically grounded models (Tomé et al., 2022; Chrysanthidis et al., 2022; Guerreiro & Clopath, 2024) often assume a pre-existing shared code between hippocampus and cortex. Consequently, these models primarily address how associations between representations can be transferred from hippocampus to cortex, rather than how semantic representations themselves emerge from episodic experience. In addition, the complexity of the memories they can accommodate is limited, requiring highly decorrelated episodic patterns for stable learning and recall. Together, these considerations suggest the need to reevaluate replay as a pure re-instantiation of cortical targets for error-driven learning and to move towards a biologically plausible account that can still uncover semantics from structured, correlated episodes.

A second unresolved tension between theory and experiments concerns the nature of the episodic code itself. Classical theories of associative memory have long emphasized that storage capacity is maximized when the stored patterns are decorrelated and approximately orthogonal to one another (Hopfield, 1982; Tsodyks & Feigel’man, 1988; Treves & Rolls, 1991). In contrast, a growing body of experimental evidence suggests that representations in the medial temporal lobe, a circuit crucial for episodic memory, contain substantial semantic and relational structure. These include concept cells that respond selectively to abstract concepts across different sensory modalities and contexts (Quiroga et al., 2005; Quiroga, 2012; Courellis et al., 2024), as well as neurons that encode latent variables and higher-order task structure (Aronov et al., 2017; C. Sun et al., 2020). Such representations necessarily introduce structured overlap between memories, apparently violating the pattern separation conditions under which episodic memory is expected to operate most efficiently. Moreover, semantically meaningful material is paradoxically often remembered more accurately than arbitrary information (Chase & Simon, 1973; Murray et al., 1976; Tse et al., 2007; Lin et al., 2021). This raises two fundamental questions: what is the origin of these semantic representations in the episodic memory circuit, and what computational role do they serve?

All in all, while views challenging a strict dichotomy between semantic and episodic memory are almost as old as the semantic-episodic distinction itself (McKoon et al., 1986; Greenberg & Verfaellie, 2010; Gentry & Buckner, 2024), we still lack a mechanistic understanding of how or why these two memory systems interact. Nevertheless, several recent developments point towards a different perspective on episodic-semantic interactions. For example, replay is increasingly viewed as a generative process, rather than providing a verbatim teaching signal for cortical learning (Swanson et al., 2020; Diekmann & Cheng, 2023; Kurth-Nelson et al., 2023). Under this perspective, neocortex may act less as a passive recipient of replayed memories and more as an active interpreter of replayed structure, extracting a set of reusable representations that efficiently explain recurring replay events. Such a process bears a strong conceptual resemblance to dictionary learning frameworks, in which complex activity patterns are decomposed into sparse combinations of reusable latent features (Olshausen & Field, 1996; Bricken et al., 2023). A complementary set of observations concerns the flow of information in the opposite direction. While consolidation theories have traditionally emphasized hippocampus-to-cortex transfer (Moscovitch & Gilboa, 2021), cortex-to-hippocampus teaching has recently been highlighted as an important and largely unexplored aspect of memory consolidation (Liu et al., 2024). Related ideas have begun to emerge in computational accounts that assign an increasingly active role to cortical semantic knowledge during replay (Van de Ven et al., 2020) and episodic encoding (Spens & Burgess, 2024a). Furthermore, experimental work has revealed mechanisms through which cortical activity can shape medial temporal lobe representations, including cortex-initiated hippocampal replay (Rothschild et al., 2017), strengthening of corticofugal projections from cortical assemblies to medial temporal circuits (Lee et al., 2023), and prefrontal signals that control the overlap and linking of episodic memories (de Sousa et al., 2026). Together, these observations suggest a framework in which replay is used not simply to transfer associations from episodic to semantic memory, but to discover reusable semantic structure from episodic experience and subsequently use that structure to shape future episodic representations.

Here, we present a model of episodic-semantic memory interactions that addresses the challenges outlined above. First, we put forward dictionary learning as the primary algorithm of semantic consolidation. Under this view, the role of *episodic replay* is not only to provide targets for cortical learning (*what should one predict in future episodes based on those stored* ), but to generate candidate recurring atomic representations for future episodic encoding (*how should one interpret future episodes that share structure with those stored* ). Second, we propose that these learned cortical representations can be transferred back into the episodic memory system. This allows semantic representations to become part of the episodic code, thus enabling a shared vocabulary in the representations used by the two memory systems. We show that these principles can be instantiated in a circuit model based on Hebbian and homeostatic local plasticity, with neocortex as the primary substrate for semantic memory and the medial temporal lobe as the primary substrate for episodic memory. We further demonstrate that the resulting dynamics reproduce key experimental observations, including concept-cell-like responses, increasing overlap between semantically related memories, and recent findings in synapse-specific changes associated with consolidation. We then investigate the computational role of semantic representations in episodic memory and find that these act as an error-correcting scaffold that benefits recall. Furthermore, semantic representations in the medial temporal circuit enhance neocortex-MTL communication during replay, further supporting semantic consolidation of episodes that are semantically in-distribution. These results provide a computational framework that can explain, unlike classic accounts of systems consolidation, why the episodic memory circuit can benefit from semantic representations. They also make specific predictions that could be tested in a statistical learning paradigm where input is compositional. All in all, we provide a mechanistic account of how semantic knowledge can emerge from complex episodic experience under biological principles, in turn shaping the organization and consolidation of future memories.

## Results

### A model for semantic structure in episodic memory based on dictionary learning

We introduce a model of semantic and episodic memory (Fig. 1a). The model starts from a simple psychological observation: episodic memories often contain both sensory detail and semantic content. For example, consider multiple episodes of someone having their favourite food: *pizza*. These may differ in appearance, smell, taste, and context, but still share semantic structure such as the presence of *pizza, cheese*, and *tomato*. Our model captures this by positing that episodic traces contain two components (Fig. 1a): a sensory component, preserving event-specific perceptual detail, and a semantic component, representing reusable concepts shared across events.

**Figure 1:**
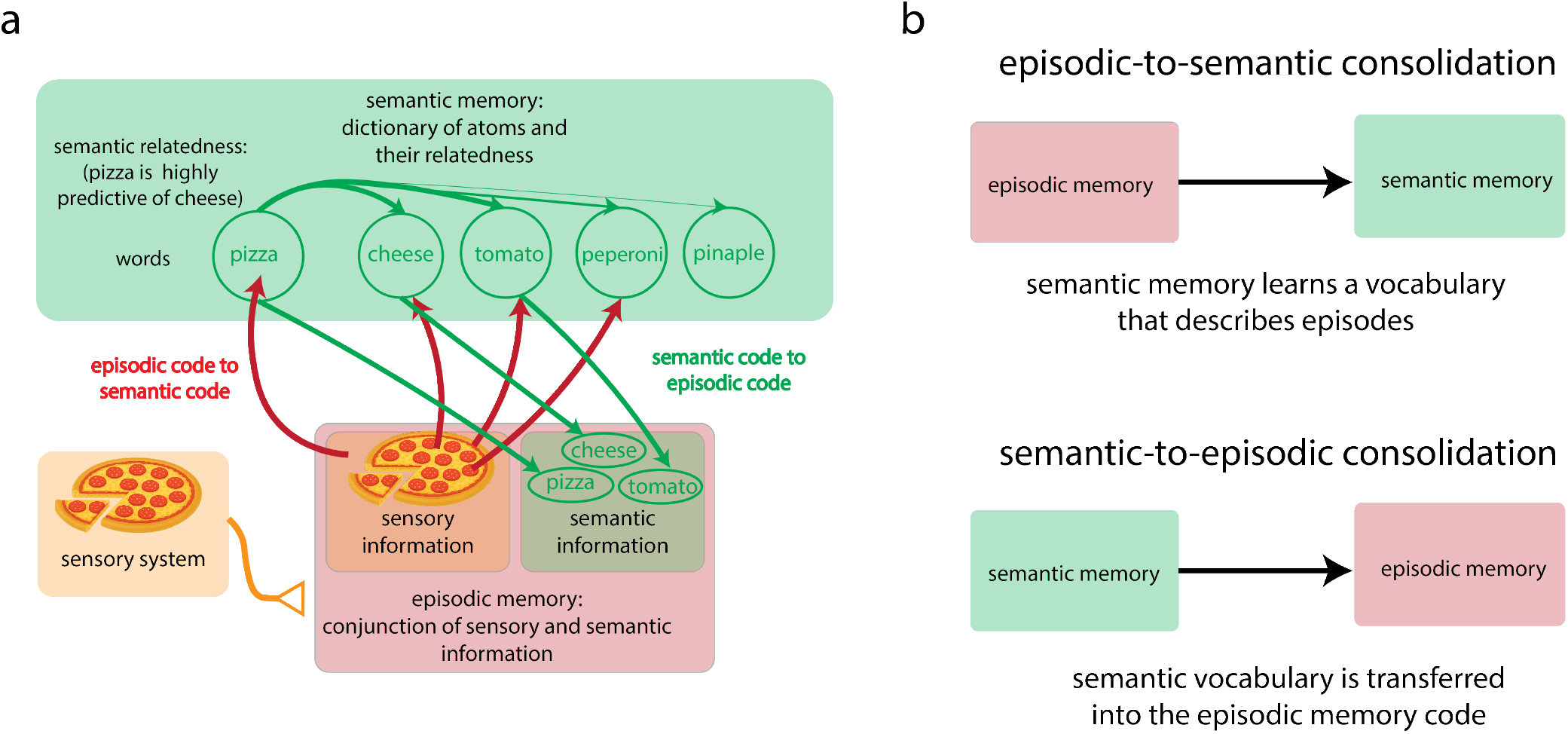
We propose a model in which replay extracts a semantic vocabulary, which is shared between episodic and semantic memory. **a**: Model of episodic (red) and semantic (green) memory. Episodic memories integrate both sensory information (orange) and semantic content (green), while semantic memory extracts regularities across episodes. **b**: Schematic of the proposed consolidation phases: Episodicto-semantic consolidation transfers regularities from episodic memory to semantic memory, while semantic-to-episodic consolidation transfers the formed semantic representations to episodic memory.

At the circuit level, however, the presence of semantic representations in episodic memory raises a computational problem. Although the medial temporal lobe (MTL), which is crucial for episodic memory, has been shown to contain concept-like information, classical attractor models achieve maximal storage capacity when stored patterns are maximally distinct. Semantic representations, by contrast, introduce structured overlap between episodes. At the same time, behavioural evidence points in the opposite direction: semantically comprehensible episodes are better recalled, and lesions in MTL regions such as the hippocampus also impair recall of already consolidated semantic information. How can the episodic memory system not only tolerate this structured overlap but actually benefit from it? To address this question, we develop a complementary circuit implementation of the model to study how semantic information can coexist with episodic storage in MTL. Following standard mappings, semantic memory is assigned to neocortex (CTX) and episodic memory to MTL (Fig. S1). MTL is divided into MTL-sensory, driven by sensory input, and MTL-semantic, driven by semantic cortical input. These populations are recurrently connected, allowing sensory and semantic representations to be bound into individual episodic traces. All regions are modeled as binary networks with homeostasis (see Technical Box 1 and Methods) because of their analytical tractability and direct connection to semantic learning (Albesa-González & Clopath, 2025).

A related question that can be asked with the circuit model is how the cortical semantic representations are formed in the first place. A common framing is that replayed cortical reactivations are used as a teaching signal, which can then be backpropagated. However, this often leaves implicit how the reactivated episodic pattern interacts with cortical activity. Which cortical neurons provide the input? Which ones define the target? Which ones remain plastic? Even biologically motivated alternatives to backpropagation, such as Contrastive Hebbian Learning or Predictive Coding, still require a structured distinction between input, hidden, and output components during learning.

Here, we avoid this problem by proposing replay as a mechanism for dictionary learning (Technical Box 2). Under this view, the role of *episodic replay* is to provide episodic patterns that CTX must interpret, which is achieved via Hebbian plasticity and competitive learning. This avoids backpropagation-like credit assignment and the need to predefine cortical targets. Furthermore, the replayed pattern is not a verbatim copy of awake experience, but a generalized episodic pattern. When projected to CTX, this pattern either refines existing representations or recruits new neurons if no previous representation is sufficiently close.

The same mechanism can also explain how CTX transfers the learned representations to MTL. If recurrent MTL dynamics expose episodic structure to CTX, recurrent CTX dynamics can expose semantic structure to MTL. We call this process *semantic replay* because, like episodic replay, it is internally generated, begins from random activity, and recovers attractor states through recurrent dynamics. However, because these attractors are shaped by CTX recurrent connections, they reflect semantic rather than episodic structure. By projecting them to MTL-semantic, concept-like representations can enter the episodic circuit.

Together, episodic and semantic replay implement in the circuit the learning dynamics proposed by the model (Figs. 1b and S1). Episodic replay supports *episodic-to-semantic consolidation*, allowing CTX to extract a cortical dictionary from episodic structure. Semantic replay supports *semantic-to-episodic consolidation*, allowing MTL-semantic to incorporate the learned cortical vocabulary. We next validate this learning algorithm in simulations of the circuit model. We then use the model to ask how semantic overlap can interact beneficially with episodic attractor dynamics, supporting memory and further consolidation, and connect our results with previous experimental findings.

#### Technical Box 1: Homeostatic Binary Networks

All regions in the circuit model are implemented as Homeostatic Binary Networks, following Albesa-González and Clopath (2025). These networks are binary, with sparsity in each region being controlled by a *K-winners-take-all* mechanism, implemented with a *top*-*K* activation function. For further details and definitions of variables, see Methods.

**Neural dynamics**. Each region is described by a binary activity vector

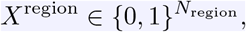

with exactly *K*^region^ active neurons at each timestep. Synaptic input is first integrated into a pre-activation variable

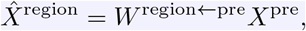

which is then mapped into binary activity by competition:

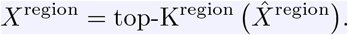

This mechanism enforces competition across neurons to represent the current synaptic input.

**Local learning**. Synapses are updated by a Hebbian rule,

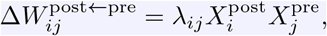

followed by divisive normalization (homeostasis), which prevents runaway growth and keeps the total synaptic strength bounded.

**Homeostatic regimes**. We use two forms of synaptic homeostasis. With *incoming homeostasis*, weights are normalized so that the total input onto each postsynaptic neuron is bounded. With *outgoing homeostasis*, weights are normalized so that the total output from each presynaptic neuron is bounded. Under these two regimes, synaptic weights converge to scaled conditional probabilities:

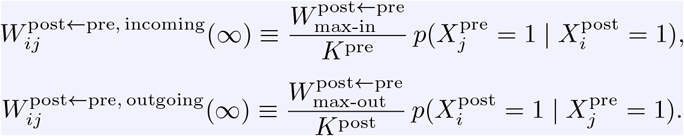

Thus, incoming homeostasis makes synapses approximate the average presynaptic pattern associated with a postsynaptic assembly, whereas outgoing homeostasis makes synapses reflect how strongly presynaptic neurons predict postsynaptic recruitment.

**Functional role**. In recurrent connectivity, these dynamics implement pattern completion. In feedforward connectivity, they implement competitive dictionary learning, causing neurons to become selective to stable components of their input.

#### Technical Box 2: Replay for Dictionary Learning

Instead of viewing replay as perfect reinstatement of stored episodes, we propose it can be seen as a mechanism for dictionary learning, where a *post* region learns relevant *atoms* describing a *pre* region based on presynaptic recurrent connections. The mechanism is as follows:

**Generate a presynaptic pattern** from noise via attractor dynamics:

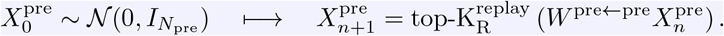

**Project to postsynaptic layer**

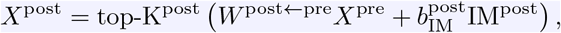

**Update dictionary** For postsynaptic active neuron 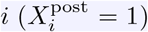.

- If neuron *i* is immature 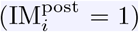: learn new atom

**–** Create a one-shot receptive field 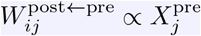

→ neuron *i* becomes maximally selective to *X*^pre^

**–** Make neuron *i* mature 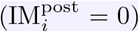

- If neuron *i* is mature 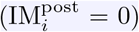: update existing atom

**–** The receptive field is pushed in the direction of *X*^pre^

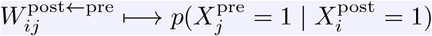

### A neocortex-medial temporal lobe model can form semantic memories via dictionary learning

#### Neocortex forms a dictionary of reusable components via episodic replay

We first consider episodic-to-semantic consolidation (Fig. 2a) as a form of dictionary learning carried out by neocortex. In dictionary learning, complex inputs are represented by combining a small number of reusable elements (typically called *atoms*, Technical Box 2). Here, we refer to these learned elements as *cortical words*: reusable cortical representations that capture recurring parts of episodic experience. In this view, episodic replay in MTL does not simply reinstate full episodes. Rather, sparse replay reveals recurring subpatterns embedded across many episodes. CTX then either matches each replayed candidate to a previously learned cortical word, refining the corresponding receptive field, or treats it as sufficiently novel to recruit immature cortical neurons and establish a new receptive field. In this way, repeated episodic replay gradually builds a cortical dictionary of reusable semantic components (words).

**Figure 2:**
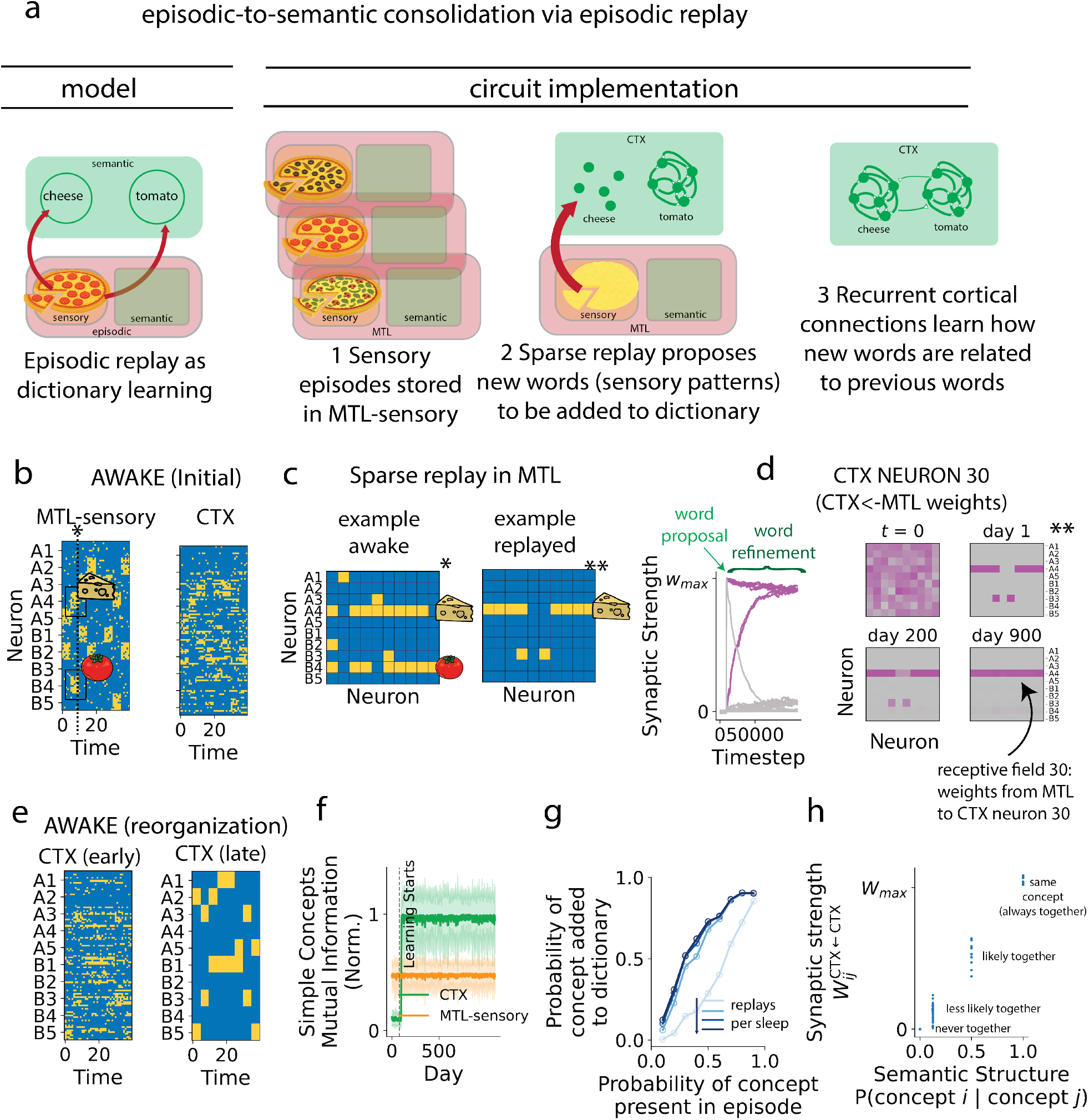
Sparse episodic replay creates compositional representations in semantic memory. **a**: Episodic-to-semantic consolidation via episodic replay. Model (left) and circuit implementation (middle/right). Sensory episodes (no semantics extracted yet) are stored in MTL-sensory, replayed sparsely during sleep to uncover overlapping sub-patterns, and then projected to CTX, where compositional semantic structure is extracted in CTX ←MTL (episodic replay). Predictive relations are slowly embedded within CTX ←CTX synapses. **b**: Example neuronal activity during wake. Input episodes are sampled following a protocol as described in Albesa-González and Clopath (2025). To determine MTL-sensory activity, two different concepts (one *A*_*i*_ ∈ *A* and one *B*_*j*_ ∈ *B*) are simultaneously sampled. 50% of episodes are of the form (*A*_*i*_, *B*_*j*=*i*_), and 50% (*A*_*i*_, *B*_*j*=*i*_). This makes *A*_*i*_ more semantically related to *B*_*i*_ than the other *B*_*j*_ values. 10 neurons in MTL-sensory encode each concept, and each episode lasts 5 timesteps. At each timestep, *N*_swaps_ = 4 random neurons that code for the present concepts are turned off, and *N*_swaps_ = 4 random neurons that code for non-present concepts are turned on (this introduces noise with a consistent signal-to-noise ratio). CTX activity is driven by random initial CTX ←MTL-sensory connections. **c**: Example neuronal activity (one timestep) during wake (left) and episodic replay (right). Sparse replay in MTL-sensory reveals concept-specific sub-patterns (for example, here *A*_4_). Figure 2: *(continued)* **d**: Evolution over time of the receptive field (CTX ←MTL synapses) of an example neuron. Right: weight dynamics for all incoming synapses from MTL-sensory onto the same neuron. Left: Heatmaps show weights associated with each presynaptic MTL-sensory neuron, rearranged in a rectangular pattern according to the concept they code for (each row represents weights from neurons coding for a particular concept in MTL-sensory). Initially random connectivity (*t* = 0) is driven in one shot to represent a replayed presynaptic pattern (day 1), and neuron 30 goes from immature to mature. Then, slow Hebbian and homeostatic plasticity drive statistical learning, uncovering receptive fields that represent semantic components (days 1 to 900). **e**: Comparison of early (before learning) and late (after learning) cortical activity, given the same sensory input. In the late phase, CTX neurons now respond selectively to semantic components *A*_*i*_ and *B*_*j*_. **f** : Evolution over time of the normalized mutual information of cortical and MTL-sensory activity with simple concepts (e.g. *A*_1_), averaged across concepts. **g**: For different marginal probabilities of a concept being present in an episode (horizontal axis), probability of a receptive field selective to that concept being added in CTX during one session of sleep (vertical axis). Darker colors indicate increased number of replay events during sleep. **h**: For CTX ←CTX connections, relationship between synaptic strength and semantic structure (right). Recurrent connectivity captures conditional firing probabilities across concepts.

We first test whether CTX can implement such an algorithm in the model. To do so, we generate episodes by compositionally combining latent factors *A* and *B*, (*A*_*i*_, *B*_*j*_), with *A*_*i*_ ∈ {*A*_1_, …, *A*_5_} and *B*_*j*_ ∈ {*B*_1_, …, *B*_5_} (Figs. 2b and S2, see Methods). Each latent factor can take different values (e.g. *A*_4_, or *B*_4_), each of which corresponds to a concept –a semantically coherent set of sensory features– such as *cheese* or *tomato* (Fig. 2b). The distribution over pairings is structured: matched pairs, such as (*A*_3_, *B*_3_), account for 50% of episodes, whereas all mismatched pairings jointly account for the remaining 50%. Thus, although every *A*_*i*_ can in principle appear with every *B*_*j*_, factor values with matching indices co-occur more often. For example, *A*_4_ is more predictive of *B*_4_ than of *B*_3_.

During wake, episodic inputs are stored in MTL through Hebbian plasticity. During sleep, sparse pattern completion (Technical Box 2) in MTL recovers sub-patterns shared across episodes (e.g. most active neurons correspond to *B*_2_), rather than reinstating complete episodic states (Fig. 2c). These replayed subpatterns are projected to CTX through initially random CTX ←MTL connections (Fig. 2d, *t* = 0). If the replayed pattern is not sufficiently close to existing cortical receptive fields (i.e. no CTX neuron is strongly tuned to the current MTL pattern), cortical activity is biased toward immature neurons (Methods). Because these neurons are more excitable, they are preferentially recruited, forming a one-shot cortical representation of the replayed MTL pattern (Fig. 2d, *word proposal*, day 1). However, when the replayed pattern is sufficiently close to an existing cortical receptive field, the corresponding cortical word is reactivated and refined (Fig. 2d, *word refinement*, from day 1 on). This process slowly tunes cortical neurons to stable, de-noised components of episodic experience (Fig. 2d (day 900) and Fig. 2e). It should be noted that, throughout the study, the only population with a predefined (partial) tuning is MTL-sensory: other labels (e.g. *A*_3_ in CTX) come from examining the selectivity of neurons after learning, and not from a pre-specified encoding (see *Neuronal Ordering* in Methods).

We further confirm the representational change in CTX by measuring the mutual information between cortical activity and the underlying *simple concepts, A*_*i*_ and *B*_*j*_. Cortical activity rapidly becomes maximally informative about latent factor values, reaching a normalized mutual information of approximately 1 (Fig. 2f). *Complex concepts* of the form *A*_*i*_*B*_*j*_ are considered in later sections. We also find that the probability of learning a cortical word increases monotonically with the probability that its corresponding simple concept appears in an episode (Fig. 2g). Thus, this mechanism can be interpreted as a form of prioritizing replay for generalization: frequently recurring compositional elements are more likely to be consolidated into cortical words (semantic representations).

#### Cortical recurrent connectivity represents semantics as predictability across cortical words

In our model (Fig. 1a), we proposed that semantic memory contains information about the relatedness between concepts, which is defined based on their directional predictiveness (e.g. pizza is highly predictive of cheese). In the circuit implementation, when multiple cortical neurons become tuned to the same replayed pattern, they also become strongly recurrently connected, forming a cortical assembly for that word (Methods). This corresponds to the standard replay-mediated formation of recurrent cortical assemblies, in which reactivated neurons are bound into a stable attractor-like representation. However, during wake, slow plasticity further updates recurrent connections across the broader cortical population. Under Hebbian plasticity with outgoing homeostasis, these recurrent weights converge to the conditional firing structure between neurons (Albesa-González & Clopath, 2025). Because cortical neurons are tuned to individual concepts, recurrent connectivity comes to reflect how predictive concepts are of one another (Figs. 2h and S3b). Thus, while feedforward CTX ←MTL weights learn the dictionary entries—what each cortical word encodes— recurrent CTX ←CTX connections learn the relations between entries—how each word is situated within the broader semantic structure. Importantly, this allows storing asymmetric semantic relationships (for example, pizza is much more predictive of cheese than *vice versa*), as we will see in the following sections.

Together, these results suggest that semantic memory can emerge from episodic replay through cortical dictionary learning. Replay extracts candidate words from overlapping episodes, CTX stabilizes them as reusable semantic units, and recurrent cortical connectivity organizes them according to their co-occurrence statistics. In the pizza example, cortical words encode ingredients such as cheese or tomato, whereas corticocortical connections encode the predictive relationships between ingredients.

### Semantic replay can transfer the learned vocabulary into episodic memory

We next ask whether spontaneous cortical activity can transfer the learned cortical dictionary back into episodic memory (semantic-to-episodic consolidation; Fig. 3a, left). We refer to this mechanism as *semantic replay* because it parallels episodic replay: it is driven by spontaneous activity, with activity patterns recovered through attractor dynamics over initial random states. However, here the replayed content comes from recurrent connections representing *semantic* information, as opposed to episodic. In the pizza example, once ingredients such as tomato and cheese have been consolidated as cortical words, semantic replay can imprint these words onto MTL-semantic. Thus, before semantic-to-episodic consolidation, an episode containing tomato sauce may be represented only through sensory features. After consolidation, MTL-semantic additionally contains neurons that explicitly signal the presence of tomato, analogous to MTL concept cells. At the circuit level, this corresponds to MTL-semantic neurons developing receptive fields that are selective for cortical assemblies representing individual concepts (Fig. 3a, right).

**Figure 3:**
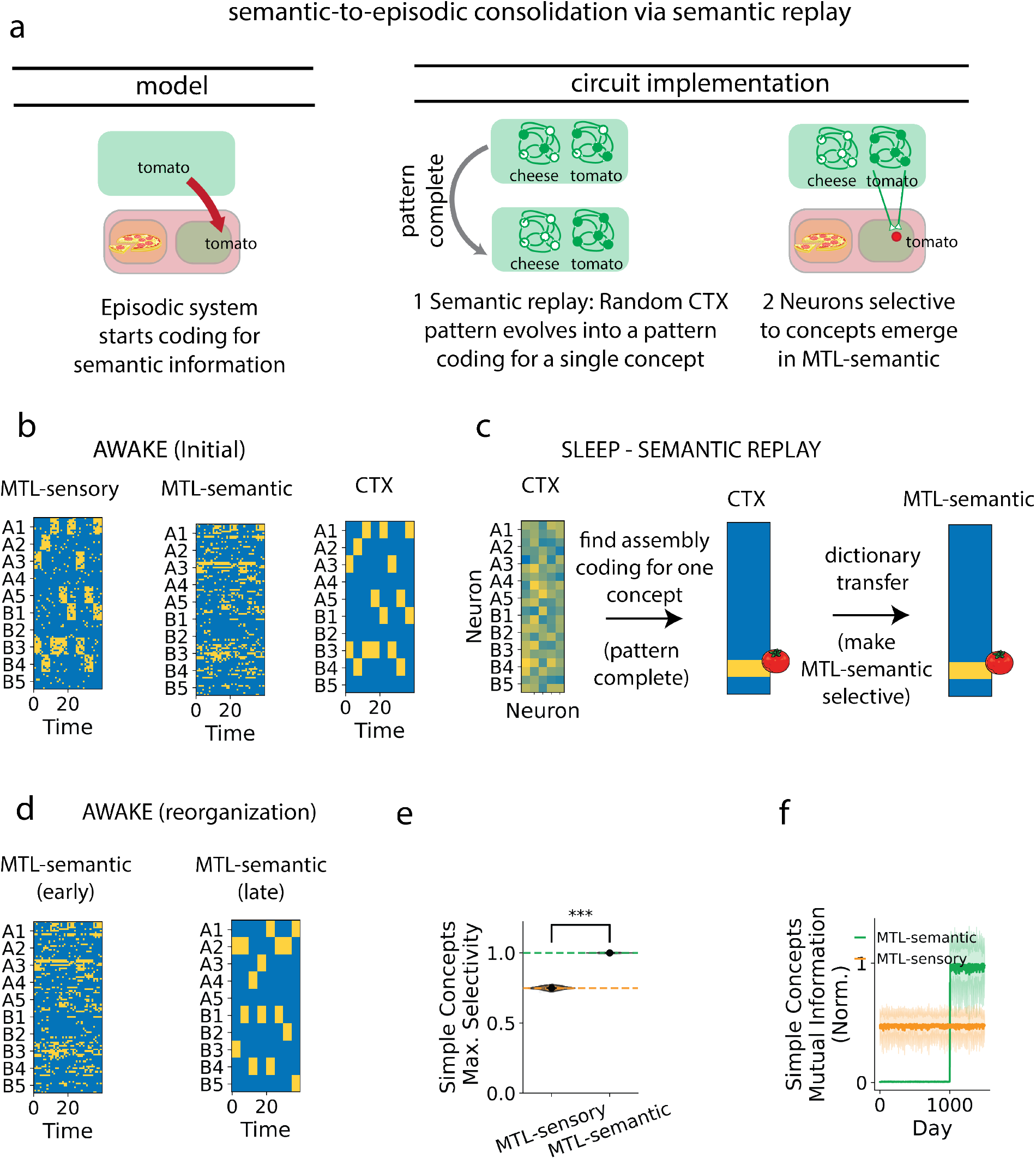
Semantic replay forms abstract representations in episodic memory. **a**: Semanticto-episodic consolidation via semantic replay. Model (left) and circuit implementation (right). During semantic replay, a random CTX pattern converges to a concept-specific attractor, which is then projected to MTL-semantic, creating semantic-specific receptive fields in MTL-semantic and enabling episodic traces to incorporate semantic information. **b**: Example neuronal activity before semantic replay starts. CTX has organized its activity around semantic components, but MTL-semantic is still not tuned to semantic representations. **c**: Example neuronal activity during semantic replay. A random initial CTX state evolves into a complete pattern coding for a single concept, which drives activity in MTL-semantic. A combination of one-shot and slow learning again causes neurons that are initially slightly more sensitive to cortical patterns to become selective to cortical semantic representations. **d**: MTL-semantic reorganization with semantic replay. **e**: Selectivity of neurons in MTL-sensory and MTL-semantic populations (*** *p <* 0.001). **f** : Evolution of mutual information between different MTL-subregions and simple concepts (averaged across concepts).

To support this process, we now equip the simulated model with semantic replay (Algorithm 8). After the first 1000 days of training, the network learns semantic representations in CTX, but MTL-semantic activity depends on random initial connectivity (Fig. 3b). After that, during sleep, semantic replay initializes CTX from random activity and allows recurrent cortical dynamics to recover learned semantic attractors. As in the previous section, sparsity is increased to isolate patterns corresponding to individual concepts (Fig. 3c). These patterns are then projected to MTL-semantic through initially random connections. As a result, receptive fields in MTL-semantic come to represent individual concepts identified by CTX (Fig. 3d). Consequently, while MTL-sensory exhibits a mixed selectivity to different concepts, MTL-semantic neurons show an absolute selectivity to these concepts (Fig. 3d, e), with mutual information becoming maximal rapidly after semantic replay is initiated (Fig. 3f). This reflects not only that the cortical semantic dictionary is maximally informative about episodic latent factors, but that the cortical code is effectively transferred into MTL-semantic.

Once MTL-semantic has acquired concept-selective receptive fields, replay can engage both MTL-sensory and MTL-semantic populations through recurrent MTL connectivity (Fig. S5). As a result, subsequent replay events now contain both sensory and semantic information, inherited from CTX. While semantic representations are identical in content in CTX and MTL, they serve different purposes. In CTX, recurrent connectivity between semantic assemblies stores long-term predictiveness across concepts. However, in MTL-semantic, recurrent connectivity stores the semantic associations of the episodes stored during the day. Extending replay in MTL to joint sensory-semantic episodes expands the previous CTX ←MTL-sensory connections to form cortical receptive fields that jointly respond to MTL-sensory and MTL-semantic patterns (Fig. S3d).

Overall, these results show that semantic replay implements semantic-to-episodic consolidation in the circuit model, as proposed in the model. This provides a mechanistic account of how learned semantic representations can be transferred from neocortex to MTL, giving rise to concept-like cells (Quiroga et al., 2005; Quiroga, 2012), and providing CTX and MTL with a shared vocabulary of semantic representations.

### Variable sparsity during replay controls the complexity of the vocabulary learned

So far, episodic replay has been constrained to high sparsity, a regime that decorrelates episodic subpatterns and promotes the extraction of *simple concepts* (e.g. tomato or cheese). However, if replay is allowed to recruit a larger number of active neurons (decreasing sparsity), the network can in principle recover patterns corresponding to higher-order combinations of concepts. For example, replay may recover conjunctions of concepts such as *crust, cheese*, and *tomato* by allowing a less sparse pattern to emerge (Fig. 4a).

**Figure 4:**
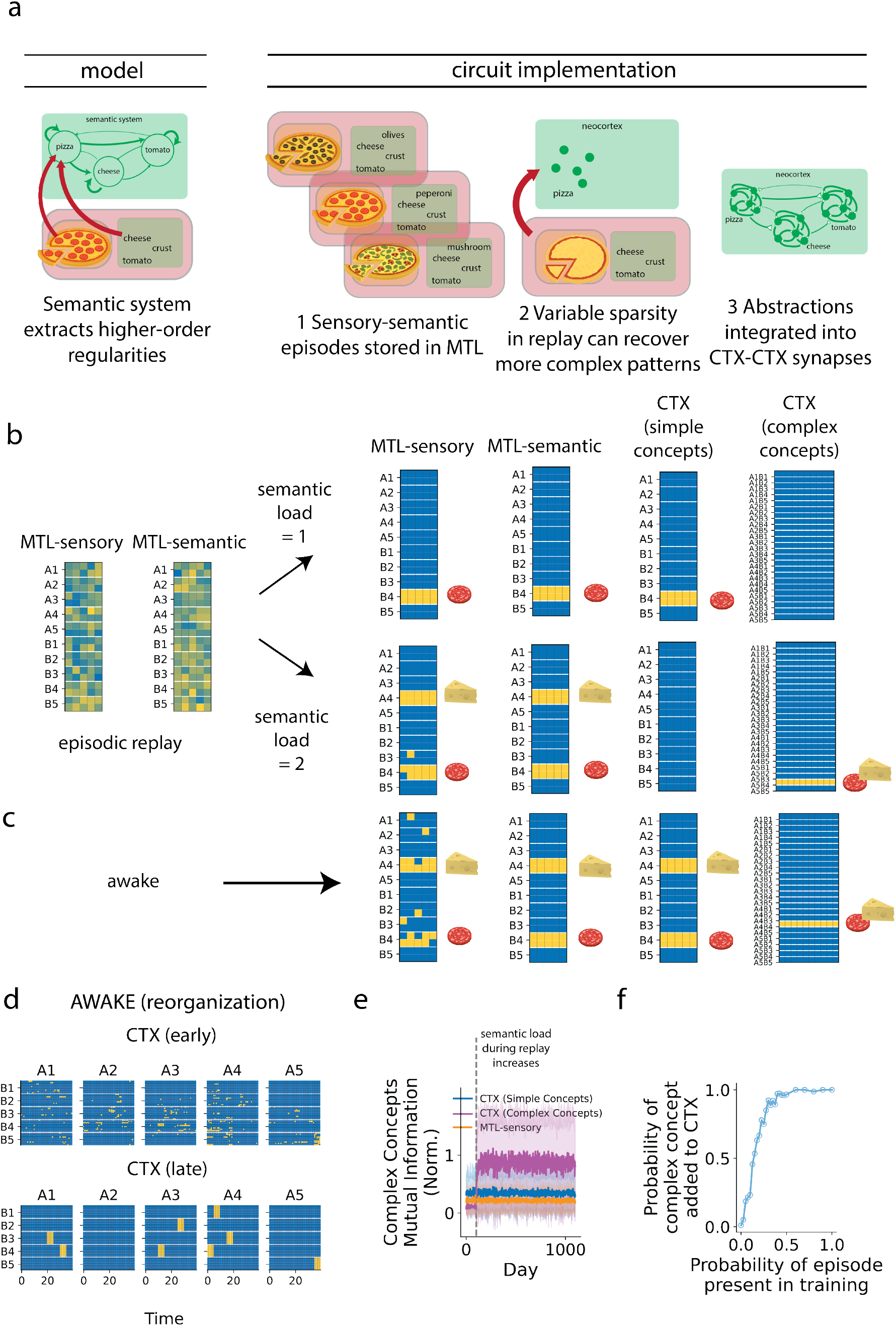
Learning complex concepts based on simple concepts with variable levels of sparsity. **a**: Model (left) and circuit implementation (right) of higher-order concept learning. Multiple sensory–semantic episodes are stored in MTL. Variable sparsity during replay (semantic load) allows the recovery of higher-order associations, which are then consolidated into CTX ←MTL and CTX ←CTX connections. **b**: Example activity during replay with semantic load 1 (top) or 2 (bottom). When one concept is replayed, CTX develops selective coding for single concepts. When two concepts are replayed simultaneously, CTX develops receptive fields for their combinations, supporting higher-order abstractions. **c**: Example activity during wake after training. CTX contains neurons selective to individual concepts *A*_*i*_ and *B*_*j*_, as well as to combinations (*A*_*i*_, *B*_*j*_). **d**: Learning-related reorganization of CTX activity (complexconcept-selective neurons). Conjunctive representations duplicate representations of *B*_*j*_ depending on the *A*_*i*_ context (and can also be interpreted in reverse: they encode *A*_*i*_ in a *B*_*j*_-specific context). **e**: Evolution of normalized mutual information with complex (e.g. *A*_1_*B*_3_) concepts across different regions, averaged across complex concepts. **f** : Same as Fig. 2g, applied to complex concepts.

To test this idea, we relax the previous assumption of the model that replay is necessarily sparse. Instead, we define the *semantic load* of a replay event as the ratio between the number of active neurons in a replay event and the minimum number permitted in that region, with previous sections corresponding to a semantic load of 1. Given that a semantic load of 1 tunes cortical neurons to simple concepts such as *A*_1_, we call this minimum number 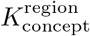 (see Methods). This allows re-interpreting the number of active neuronsin a region 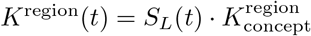 as the co-activation of *S*_*L*_(*t*) neuronal assemblies, each representing a well-defined concept. Thus, the intuition of the semantic load is to control how many concepts are simultaneously expressed in a replayed pattern. In our model, replay events have a semantic load of either 1 or 2 (Fig. 4b). Consequently, MTL attractor dynamics recover a pattern associated with either a single concept (e.g. *A*_1_) or a conjunction of two concepts (such as *A*_1_*B*_2_), depending on how many neurons are allowed to remain active. The resulting MTL activity is then projected to CTX, where a separate cortical subregion learns these higher-order associations.

Under this regime, replay with a semantic load of 2 induces a competition between neurons coding for specific combinations (*A*_*i*_, *B*_*j*_), that is, complete episodic conjunctions rather than isolated concepts. Over time, CTX develops selective receptive fields for these combinations (Figs. 4b–d, *complex concepts*, and Fig. S3d). As a result, cortical activity during wake contains neurons tuned to each *A*_*i*_, to each *B*_*j*_, and to specific conjunctions (*A*_*i*_, *B*_*j*_) (Fig. 4c). In this way, cortical dictionary learning extends from simple cortical words (*atoms* represent simple concepts) to higher-order compositional words (*atoms* represent complex concepts).

When examining CTX neurons selective to complex concepts, we find that learning reorganizes activity into a conjunctive code, at each timestep representing an abstracted individual episode *A*_*i*_*B*_*j*_ (Fig. 4d). Consistently, the mutual information between these neurons and conjunctions of simple concepts, of the form *A*_*i*_*B*_*j*_, increases sharply once variable sparsity is introduced (Fig. 4e). By contrast, the normalized mutual information between CTX simple-concept neurons or MTL-sensory activity and these complex latent values remains notably below maximal (Fig. 4e). Similarly to the case of simple concept vocabulary learning, the formation of new receptive fields is favoured by a higher probability of the corresponding episode being sampled (Fig. 4f).

This transformation is also reflected in corticocortical connectivity. As higher-order cortical words form, CTX recurrent synapses encode the relational structure between simple concepts and their conjunctions (Fig. S3e). It should be noted that the asymmetry of this structure is inherited by the connectivity matrix: a conjunction such as *A*_1_*B*_1_ strongly predicts its constituent *A*_1_, whereas *A*_1_ only weakly predicts the specific conjunction *A*_1_*B*_1_ (Fig. S3e). Consequently, recurrent connectivity can encode directional predictive relationships, overcoming a limitation of many Hebbian models that are restricted to symmetric relationships such as correlation.

Together, these results show that replay sparsity controls the level of abstraction learned by the cortical dictionary. Highly sparse replay favours isolated concepts, whereas less sparse replay allows CTX to consolidate higher-order cortical words corresponding to concept conjunctions. In the pizza example, replay may first extract individual ingredients such as *cheese, tomato*, or *crust*, and later consolidate pizza as their conjunction, capturing increasingly episodic-like semantic structure.

### The model reproduces key neural and synaptic experimental signatures

We next examine whether the circuit emerging from the proposed learning algorithm reproduces key experimental observations. To this end, we compare the representations, connectivity, and learning dynamics of the model against a range of experimental findings focusing on the neocortical-hippocampal dialogue.

#### Formation of concept cells in MTL

We first test whether semantic structure in the model reshapes medial temporal coding in a manner consistent with the defining properties of concept cells, namely concept-level representations that generalize across contexts (Fig. 5a). A central commitment of the model is that semantic abstraction is represented within the episodic circuit itself, rather than emerging only at a downstream cortical readout. This predicts that MTL populations should support concept-level generalization across contexts. Consistent with this, concept information is more readily decoded across held-out contexts from MTL-semantic than from MTL-sensory (Fig. 5b1). At the single-neuron level, MTL-semantic units also show stronger selectivity to simple concepts (Fig. 5b2) and carry more information about their preferred concept (Fig. 5b3).

**Figure 5:**
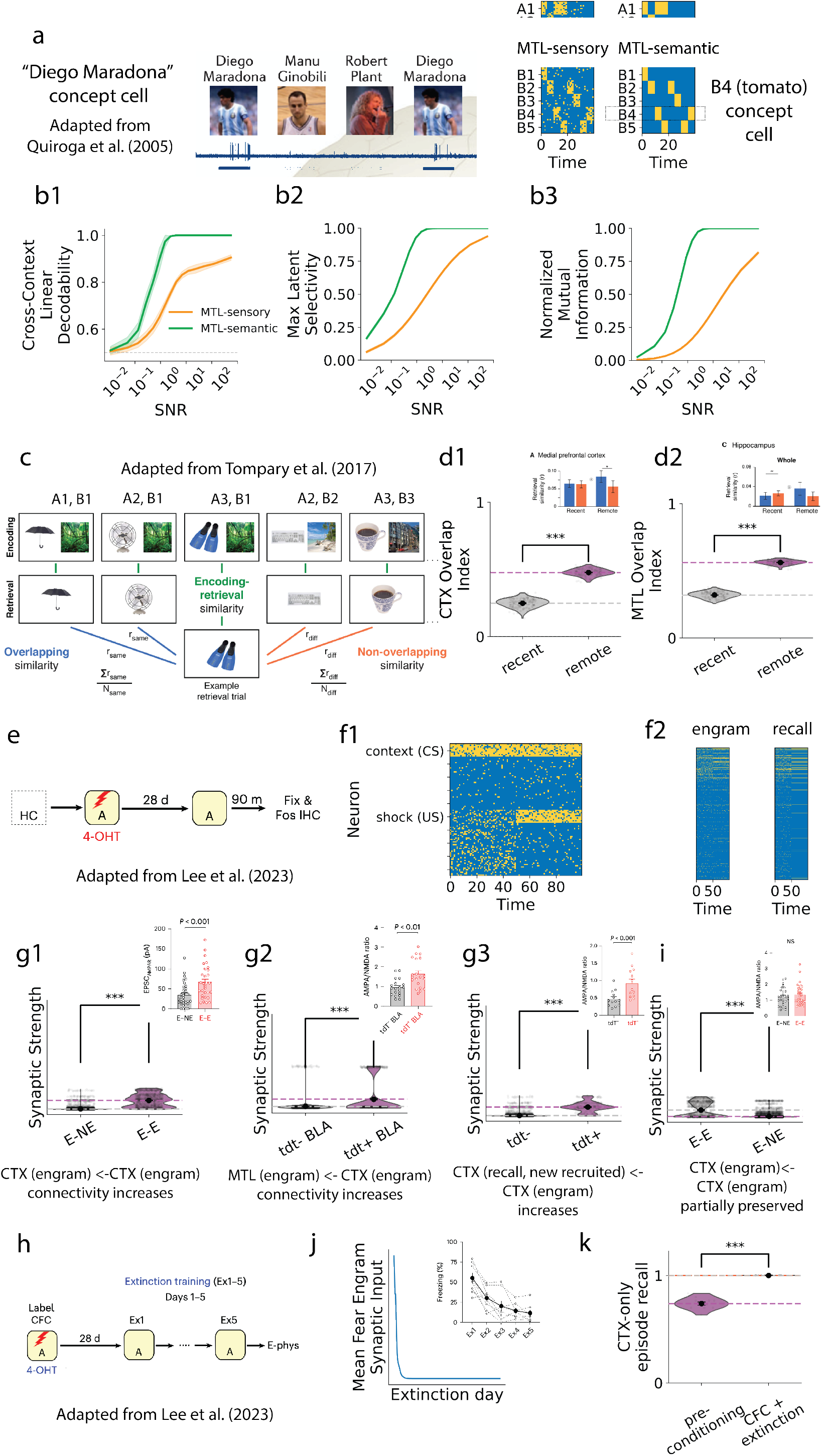
The model reproduces key neural and synaptic signatures associated with semantic memory consolidation and concept representations. **a**: Left: Example of a human medial temporal lobe concept cell selective for the concept “Diego Maradona”, responding across different visual instances while remaining largely inactive to other familiar individuals. Adapted from Quiroga et al. (2005). Right: Extrapolation to our paradigm, where *B*_4_-selective cells emerge in MTL-semantic. **b1**: Cross-context linear decoding of concepts from MTL activity. Concept identity can be decoded across held-out sensory contexts more accurately from MTL-semantic than from MTL-sensory representations, indicating stronger context-invariant coding. **b2**: Maximum selectivity to simple concepts of individual neurons (selectivity to their preferred concept). MTL-semantic neurons exhibit higher selectivity for the different values of the latent factors than MTL-sensory neurons do. **b3**: Normalized mutual information between neuronal activity and simple concept indicators (binary variables indicating the presence/absence of each simple concept). MTL-semantic neurons carry substantially more information about their preferred concept, consistent with concept-like representations. **c**: Experimental paradigm adapted from Tompary and Davachi (2017). Multiple object memories share overlapping scene associations, allowing the assessment of consolidation-induced changes in representational similarity between related memories. **d1**: Representational overlap in neocortex (CTX) before and after consolidation. Memories sharing latent structure become more similar after consolidation, reproducing the increased overlap observed experimentally for related memories. **d2**: Representational overlap in the medial temporal lobe (MTL) before and after consolidation. Consolidation similarly increases overlap between memories sharing common latent factors within episodic circuitry. **e**: Contextual fear-conditioning paradigm adapted from Lee et al. (2023). A conditioned context (CS) is paired with a fear-related unconditioned stimulus (US), enabling analysis of consolidation-induced circuit reorganization. **f1**: Example activity used in the fear-conditioning simulation. Context representations are paired with an interoceptive latent factor (that can take values such as *fear* ) during conditioning. **f2**: Cortical population activity during encoding (engram) and remote recall. Consolidation reorganizes cortical recruitment, producing only partial overlap between neurons active during encoding and those active during subsequent recall. **g1**: Recurrent cortical connectivity after consolidation. Synapses between cortical engram neurons (E–E) become stronger than connections involving non-engram neurons, reproducing experimentally observed engram-specific strengthening. **g2**: Corticofugal connectivity from cortical engram neurons to MTL engram neurons. Consolidation strengthens projections from cortical assemblies recruited during learning onto episodic-memory populations. **g3**: Connectivity from recall-recruited cortical neurons onto cortical engram neurons. Newly recruited recall neurons exhibit stronger connections to the original cortical engram than non-recall neurons. **h**: Extinction-learning paradigm adapted from Lee et al. (2023). Fear conditioning is followed by repeated extinction sessions in which the conditioned context is presented without reinforcement. **i**: Recurrent cortical connectivity after extinction. Original experiment shows a non-significant difference across E-E and E-NE synapses. While we also see a decrease in connectivity (Fig. S6a), both synaptic populations remain different in the model. **j**: Extinction suppresses expression of the fear representation across extinction sessions. **k**: Reacquisition after extinction. Previously extinguished memories are relearned faster than naive memories, demonstrating savings arising from the persistence of the underlying cortical (semantic) circuit.

#### Consolidation increases representational overlap among memories with shared concepts, both in CTX and MTL

We next ask whether consolidation reshapes representational geometry in a manner consistent with the model’s overlap-based replay mechanism. Because replay is structured by shared representations across episodes, the model predicts that memories sharing latent structure should become more similar over consolidation, both in hippocampus and in cortex. A related effect was reported by Tompary and Davachi (2017) in an object-scene paradigm, in which multiple object memories shared a smaller number of scene associations. The authors measured representational similarity during object-cued recall before and after sleep for objects associated with overlapping or non-overlapping scenes. Importantly, this task maps naturally onto our episode generation protocol, with *A*_*i*_ corresponding to objects and *B*_*j*_ to scenes (Fig. 5c). Consistent with the experimental findings, consolidation increases the overlap index in cortex, such that objects sharing a scene become represented more similarly after sleep than before sleep (Fig. 5d1). Importantly, a corresponding increase is also observed in the model’s MTL (Fig. 5d2).

#### Engram-specific potentiation of CTX recurrent synapses, and cortex to MTL synapses

Finally, we ask whether the particular neural and synaptic changes generated by the model could be related to experimentally described circuit motifs. Recent work (Lee et al., 2023) characterized synaptic and representational changes during consolidation within a contextual fear-conditioning task, offering a particularly useful benchmark for our model because it further constrains the underlying circuit mechanisms. These experiments focused on interactions between the medial prefrontal cortex (mPFC), which we map onto the neocortex (CTX) in the model, and basolateral amygdala (BLA), which we map onto the medial temporal lobe (MTL) in the model. To capture the experimental paradigm within our framework, we interpret the latent pair (*A*_*i*_, *B*_*j*_) in a contextual fear-conditioning framework: different *A*_*i*_ represent sensory descriptions of context, with *A*_1_ serving as the conditioned stimulus (CS; *A*_1_), whereas *B*_*j*_ represent internal valence states, including fear as the unconditioned stimulus (US; *B*_1_) (Fig. 5f1). Conditioning is then modeled by presenting the CS together with random activity within neurons coding for *B*, followed within the same day by its pairing with the fear state (consistent presentation of *A*_1_*B*_1_).

Lee et al. (2023) examined how remote contextual fear memories are represented by distinguishing neurons active during conditioning from those active during later recall. In their terminology, *engram neurons* are neurons recruited during the original fear-conditioning episode, whereas *recall neurons* are neurons recruited during remote memory retrieval. These populations only partially overlap: some recall neurons are reactivated engram neurons, whereas others are newly recruited at recall. The model reproduces this reorganization of cortical recruitment following consolidation (Fig. 5f2).

When examining the connectivity motifs, the model develops stronger recurrent synapses among cortical engram neurons than between engram and non-engram neurons, reproducing the selective strengthening of engram-defined cortical connectivity after consolidation (Fig. 5g1). The model likewise reproduces stronger projections from cortical engram neurons to medial temporal engram neurons than to non-engram medial temporal neurons (Fig. 5g2), as well as stronger inputs from recall-recruited cortical neurons onto cortical engram cells than from non-recall neurons (Fig. 5g3). In the model, the strengthening of CTX →MTL projections arises from semantic replay, which repeatedly reactivates cortical semantic assemblies and transfers their structure to MTL. Thus, the experimentally observed increase from the newly formed cortical engram to BLA is consistent with the prediction that semantic knowledge is progressively incorporated into episodic representations through cortex-to-MTL teaching.

Lee and colleagues next asked about the impact of extinction on the synaptic engram (Fig. 5h). In particular, they compared engram-to-engram and engram-to-non-engram cortical synapses after fear conditioning followed by extinction, and found that the strong synaptic contrast observed after consolidation was no longer clearly maintained (Fig. 5i, original experiment). However, while the model reproduces behavioural extinction (Fig. 5j), and consistently experiences a decrease in the E-E connectivity (Fig. S6a), the cortical representations formed during consolidation are not fully erased (Fig. 5i). As a result, extinction suppresses recall expression without fully erasing the underlying representational scaffold, providing an explanation for the enhanced re-learning (savings) observed when conditioning is reintroduced after an initial conditioning-extinction sequence (Fig. 5k). It should also be noted that the difference with experimentally reported results can be explained when the number of synapses is reduced to match original measurements (Fig. S6b).

#### Link between specific model mechanisms and reproduced results

Together, these results link the core mechanisms of the model to experimentally observed neural and synaptic signatures. The cortical dictionary learning is consistent with consolidation-related increases in overlap of conceptually related memories (Tompary & Davachi, 2017) and the reorganization of cortical recruitment observed between encoding and remote recall (Lee et al., 2023). Similarly, while standard consolidation theories predict catastrophic forgetting following extinction, our model explains savings in re-acquisition as an update of the semantic structure across previously existing cortical entries (embedded in recurrent cortical connections). In turn, the transfer of semantic representations into the episodic circuit is supported by the formation of concept cells (Quiroga et al., 2005; Quiroga, 2012) and the fact that consolidation promotes overlap across conceptually similar memories not only in cortex, but also in hippocampus (Tompary & Davachi, 2017). Finally, the strengthening of corticofugal (MTL ← CTX) connections from cortical recurrent assemblies recruited during consolidation (Lee et al., 2023) is consistent with the idea that recurrent-driven spontaneous cortical activity can shape representations in downstream subcortical areas (*semantic replay*).

### Normative benefits of semantic representations in episodic memory

The emergence of semantic representations in the model’s medial temporal lobe allows us to ask what normative role semantic structure plays in episodic memory. This question revisits a longstanding tension: optimal attractor dynamics favor orthogonal representations, yet episodic representations in the medial temporal lobe are richly structured and semantically organized. Furthermore, rather than being detrimental, semantic information can facilitate recall and is usually more easily remembered than sensory detail. Our model provides a framework to examine why such structure might be beneficial, and what computational trade-offs it introduces for recall, replay, and consolidation.

To isolate the contribution of semantic representations, we compare three network modes (Fig. 6a). In the *semantics present* condition (Fig. 6a, green), the model operates normally and MTL-semantic contains concept-level representations learned from CTX. In the *semantics random* condition (Fig. 6a, gray), MTLsemantic remains active but its activity is replaced by random patterns at each timestep, controlling for the presence of additional neural activity without semantic content. Finally, in the *semantics absent* condition (Fig. 6a, orange), MTL-semantic is excluded from MTL attractor dynamics altogether, corresponding to a purely sensory episodic system. Together, these conditions can isolate the effects of semantic information from the effects of increasing the size of the episodic memory network.

**Figure 6:**
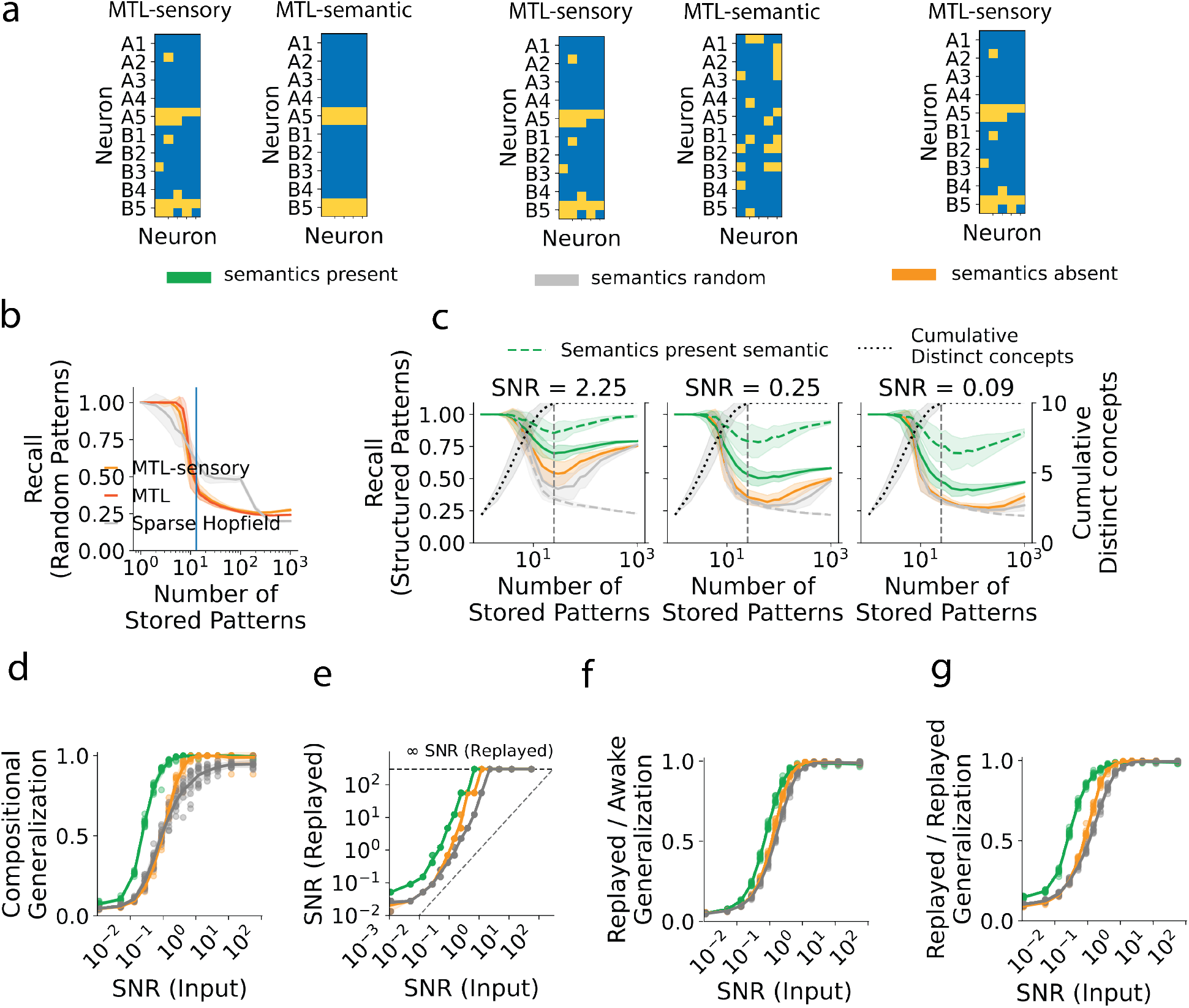
Semantic representations in episodic memory improve recall capacity and promote generalization during replay and consolidation. **a**: Three network conditions used throughout the analysis. In the *semantics present* condition, MTL-semantic contains concept representations learned from CTX. In the *semantics random* condition, MTL-semantic activity is replaced by random patterns with matched sparsity. In the *semantics absent* condition, MTL-semantic is removed from episodic dynamics, yielding a purely sensory episodic memory system. **b**: Episodic recall performance as a function of memory load for random patterns, evaluated for a standard Sparse Hopfield Network, full MTL (200 neurons), and MTL-sensory (100 neurons). **c**: Episodic recall performance with structured patterns generated from the standard structured distribution, as a function of memory load, under different levels of sensory noise (SNR = 2.25, 0.25, and 0.09). Dashed lines indicate recall from MTL-semantic representations, solid lines indicate recall from MTL-sensory representations, and the black curve indicates the cumulative number of distinct concepts encountered. Semantic representations increase effective memory capacity and improve recall of semantically structured episodes, particularly under noisy conditions. **d**: Compositional generalization of cortical representations that are conjunctions of previously learned concepts, as a function of sensory input signal-to-noise ratio (SNR). **e**: Effective signal-to-noise ratio of replayed MTL-sensory patterns as a function of input SNR. **f** : Replayed/Awake generalization. Accuracy of receptive fields derived from replay events in the classification of complex concepts across awake patterns. **g**: Replayed/Replayed generalization. Accuracy of receptive fields derived from replay events in the classification of other replayed patterns.

#### Semantic representations in episodic memory increase memory capacity when episodes are semantically in-distribution

We often recall events that we can semantically comprehend much better than those we cannot (Chase & Simon, 1973; Murray et al., 1976; Lin et al., 2021). This typically leads to an effective amnesia for what we cannot semantically process (as introduced in Tse et al. (2007)). To understand what mechanism might lead to such an effect, we begin by establishing a baseline in which no semantic structure is present. Specifically, we compare recall of purely random sparse patterns in our medial temporal lobe network against a classical sparse Hopfield network with matched size and sparsity (Fig. 6b). Under these conditions, our network shows a minor advantage at low memory loads, followed by a memory cliff as load increases. Thus, the model exhibits the standard overload regime expected for associative storage of random, unstructured patterns.

We then examine recall as the number of stored patterns increases when episodes are sampled from our standard structured distribution (Fig. 6c), across different levels of input Signal-to-Noise Ratio (SNR). One striking difference from unstructured pattern storage, common across all networks, is the presence of a U-shaped curve with the number of stored patterns. Performance initially decreases as more episodes are stored, but later recovers as repeated exposure to the same underlying structure allows recurrent weights to average out sensory noise.

Notwithstanding that, this recovery is weaker when semantic representations are absent or randomly assigned (compare recall for high number of stored patterns across networks, Fig. 6c). Without semantic representations, the dip is more pronounced, whereas networks with aligned semantic representations retain higher recall. The advantage of semantic representations (measured as the difference across networks) also increases with sensory noise (Fig. 6c). This suggests MTL-semantic provides an additional, structured population that compresses the episode into concept-level variables. This increases the number of neurons available to represent the episodic semantic content, providing an error-correcting scaffold for MTL-sensory activity. As a result, semantic representations can improve recall of the sensory component.

Finally, within the *semantics present* network, recall from MTL-semantic is substantially higher than recall from MTL-sensory (Fig. 6c, compare solid (MTL-sensory recall) and dashed (MTL-semantic recall) lines). This is expected because MTL-semantic represents the lower-dimensional conceptual structure of the episode using its own dedicated neural population, whereas MTL-sensory must preserve more variable perceptual detail. This provides a circuit-level account of why the semantic content of memories is often more robustly recalled than their sensory details, which are more prone to being distorted or forgotten.

Together, these results suggest that semantic structure optimizes the recall capacity of episodic memory by finding a compressed code that effectively amplifies the signal of the episodic conceptual structure. When stored patterns are random, the model behaves like a standard associative memory and eventually fails through overload. When episodes are semantically in-distribution, however, overlap is no longer purely interfering, and instead reflects reusable structure that MTL-semantic can encode to stabilize recall. Thus, semantic representations do not generically increase memory capacity; rather, they increase effective capacity when new episodes can be interpreted through the learned semantic vocabulary.

#### Semantic representations in episodic memory induce a compositional generalization bias in replayed and consolidated representations

Following recent theoretical studies proposing that replay optimizes generalization in Complementary Learning Systems (GO-CLS, W. Sun et al. (2023)), we hypothesize that semantic representations can play a role in the generalization of representations consolidated via replay. Thus, the benefit of semantic representations in episodic memory might not be limited to episodic recall, but even affect the semantization of future episodes that are compositions of previously learned abstractions.

First, we investigate whether the presence of semantic content during episodic encoding affects the generalization of concepts that are compositions of previously learned concepts (compositional generalization). We compare the test-time accuracy of the three network modes introduced above in classifying *complex concepts* from cortical activity, after increasing the semantic load to 2 and training for 400 additional days. As the signal-to-noise ratio of the sensory input (SNR input) decreases, the gap in compositional generalization between networks with present and absent (or random) semantic representations increases (Fig. 6d). This gap persists at intermediate noise levels, until the accuracies of both converge to chance.

To understand the origin of this cortical generalization gap, we next examine the replayed representations themselves. The first measure we obtain is the signal-to-noise ratio of the replayed pattern (SNR replayed). This is done by inferring which input-generation SNR would produce a pattern like the replayed one, considering only MTL-sensory. Semantic representations consistently increase the SNR associated with the replayed patterns, coinciding with the region in which there is a generalization gap (Figs. 6d and 6e). However, when explicitly testing the generalization of replayed MTL-sensory against awake patterns (Methods), the effect only partially explains the gap (Figs. 6f and S7a). Although the initially formed receptive field is slightly better at identifying future semantically equivalent episodes, this does not fully correlate with the generalization gap during testing. We additionally measure replayed/replayed generalization: how good are replayed patterns, when used as receptive fields, at classifying future replayed patterns? (Methods, Fig. 6g) This measure is more strongly correlated with compositional generalization (Fig. S7b).

These results can be interpreted in light of two slightly different mechanisms potentially driving better generalization after learning: when an episodic pattern is replayed, MTL is projected to CTX, and the activated cortical neurons update their MTL receptive fields in the direction of the replayed medial temporal pattern. Therefore, the formation of cortical representations during replay depends on two factors: (i) the similarity of the replayed MTL-sensory pattern to semantically equivalent future sensory inputs and (ii) whether the *right* cortical neurons are activated, which depends on the generalization across replayed patterns. That the second factor is more important for post-training generalization is consistent with the idea that cortical representations are not determined by any single replay event, but instead emerge gradually through the accumulation of many replay-driven updates under a small learning rate. As a consequence, even if semantic structure also makes individual replayed MTL-sensory patterns less noisy, the dominant factor for cortical generalization in the model is whether the *appropriate* cortical neurons are consistently recruited across replay events.

These results suggest that semantic representations promote cortical generalization not only by denoising replayed medial temporal patterns, but by providing a low-dimensional scaffold that remains tightly coordinated with cortex and helps constrain which cortical neurons are activated during replay. Together, our findings support the view that a shared semantic vocabulary across medial temporal lobe and neocortex can shape episodic replay into a teaching signal that is inherently more suitable for forming abstract, robust, and generalizable cortical representations.

### Semantic representations in the episodic memory circuit explain a range of behavioral phenomena

We finally ask whether the properties of episodic attractor dynamics characterized above can explain a broader range of behavioral phenomena (Fig. 7).

**Figure 7:**
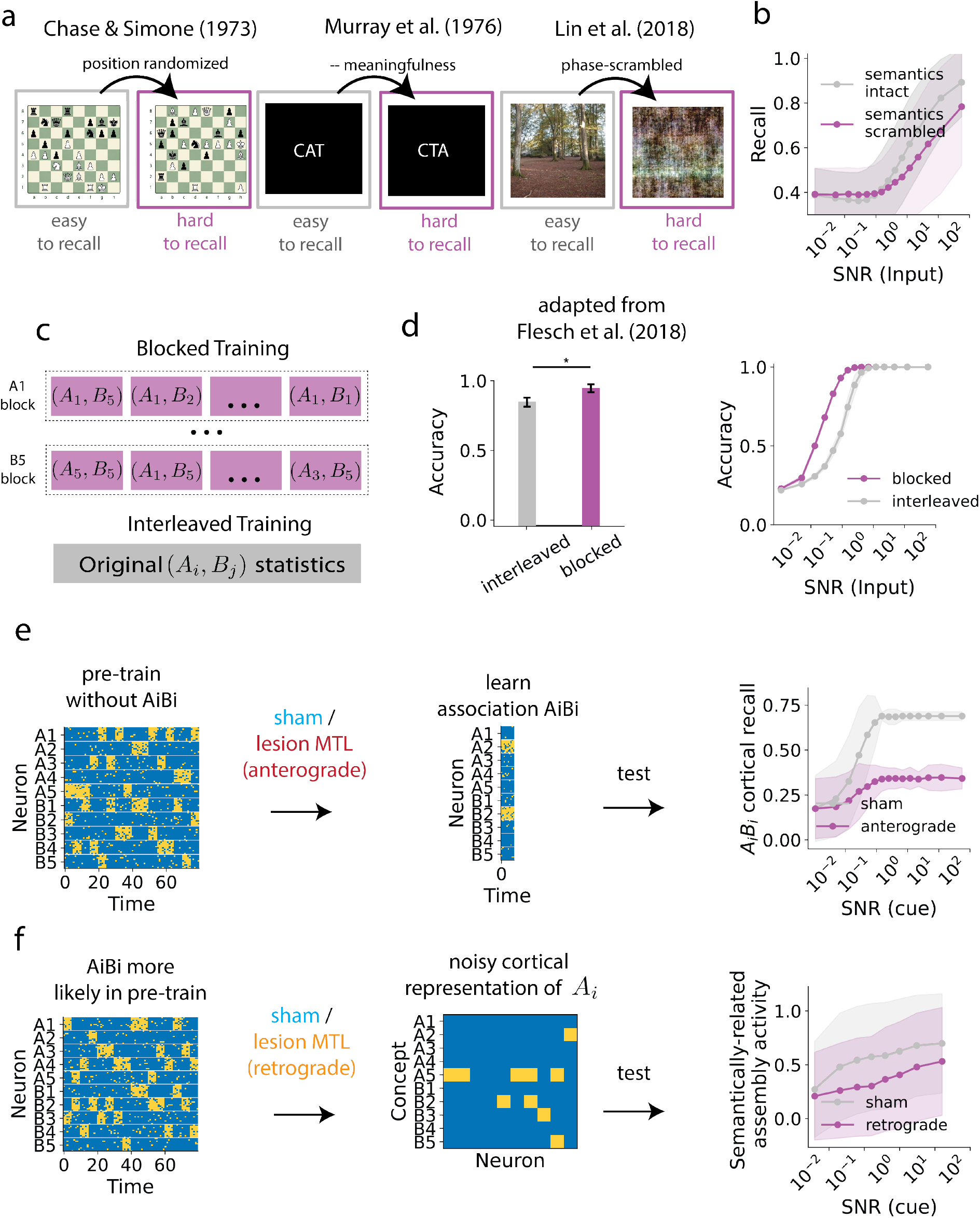
Semantic representations in episodic memory explain a range of behavioral phenomena. **a**: Schematic of previous experiments showing the benefit of recalling semantically recognizable (*intact* ) patterns over randomized (*scrambled* ) ones. **b**: Comparison of recall performance across two network conditions: one that receives input sampled from the same distribution it was trained on (*semantics intact* ) and one that receives input sampled from the same distribution but in which neuronal identity in MTL-sensory has been permuted. **c**: Blocked versus interleaved pre-training. Episodes are presented either in blocks where one concept is persistently present or in an interleaved order while preserving the overall training statistics. **d**: Effects of presentation schedule on behavioural performance in human participants (left) and in the model (right). **e**: Anterograde lesion experiment. After pre-training, a novel association is learned and medial temporal replay mechanisms are disrupted before consolidation. Subsequent cortical recall of the learned association is impaired, reproducing the classical dependence of semantic learning on MTL-mediated consolidation. **f** : Retrograde lesion experiment. Semantic statistical associations are learned during pre-training, then MTL is lesioned. Cortical s_2_e_3_mantic recall is then tested from noisy partial cues.

In the previous section, we show how semantic MTL representations structure episodic memory to benefit recall and consolidation dynamics (Fig. 6). This should be consistent with behavioural studies showing that semantically interpretable episodes are more faithfully recalled, for example when comparing real vs randomized chess positions (Chase & Simon, 1973), actual words with scrambled letters (Murray et al., 1976), or intact images with phase-scrambled ones (Lin et al., 2021) (Fig. 7a). This result lies at the core of the aforementioned tension between theory (patterns should be separated into a content-agnostic code) and experiments (structured episodic overlap is beneficial for recall). To reproduce this effect, we compare the frozen network’s recall of standard patterns (*semantics intact* ) to its recall of fixed permutations of the same patterns (*semantics scrambled* ) (see Methods). The recall performance of patterns that can be semantically interpreted is significantly higher than that of semantically scrambled versions (Fig. 7b), even though the generated episodes are equivalent except for sensory neuronal identity.

Another key difference between the model and previous accounts of semantic consolidation is that the episodic memory system performs an initial generalization step, finding structure across the different stored episodes in the medial temporal lobe. In our model, however, replayed activity is consistently de-noised with respect to sensory input (Fig. 6e). This predicts that enforcing consistent overlapping semantic components within a block benefits the formation of that particular semantic representation, which has previously been shown in humans in the literature on *blocked* versus *interleaved* learning (Flesch et al., 2018). Importantly, this result goes against standard neural network accounts of learning, and a circuit account using only biologically plausible synaptic plasticity has yet to be proposed. We therefore test a similar protocol, training two networks from scratch: one with *blocked training* (all episodes in a day contain a fixed *A*_*i*_ or *B*_*j*_), and one with *interleaved training*, where every episode is equally likely (Fig. 7c, see Methods). Across noise levels, blocked learning shows a significant advantage over interleaved learning (Fig. 7d, right), replicating previous results (Fig. 7d, left). The model suggests that blockwise learning promotes the consolidation of cortical representations for the elements within a block. This mechanism is difficult to explain with models that simply rearrange cortical recurrent connectivity toward replayed targets, a process prone to catastrophic forgetting.

Finally, we use the model to reproduce semantic memory deficits linked to lesioning of the episodic memory system. In particular, we aim to confirm the standard anterograde semantic amnesia (lesioning MTL before consolidation impairs semantic recall) and, more importantly, retrograde semantic amnesia (Duff et al., 2020). Retrograde semantic amnesia is harder to explain with classic models of semantic consolidation, because the role of MTL is only to hash cortical representations.

We first perform a standard anterograde lesion test: a new association is encoded, after which MTL recurrent and replay dynamics are disrupted before consolidation. Under this manipulation, later cortical completion of the associated concepts is markedly reduced relative to sham networks (Fig. 7e). Then, we test whether MTL-semantic also contributes to the retrieval of already consolidated semantic structure. To do so, we allow a new association to be encoded and consolidated, and only then lesion the semantic MTL pathway. We then probe cortical recall from partial cues while progressively increasing cortical noise. Sham networks can still recover the semantically related representation, whereas lesioned networks show a much steeper loss of semantically related cortical activity (Fig. 7f). Thus, while the model matches the classical role of MTL in new learning (Fig. 7e), it also captures an effect that standard one-way consolidation models fail to explain: semantic representations in MTL can continue to support semantic recall even after cortical learning has occurred (Fig. 7f).

Together, these simulations show that the same structured overlap that improves episodic storage also reproduces behavioural effects that are difficult to explain with standard consolidation models. First, the incorporation of semantic representations into episodic memory provides a natural explanation for the well-established advantage of semantically interpretable stimuli during recall, despite the increased overlap such representations introduce between memories (Chase & Simon, 1973; Murray et al., 1976; Lin et al., 2021). Second, the replay-based dictionary learning as a mechanism for consolidation predicts that learning experiences sharing a common factor value should accelerate concept formation, reproducing the behavioural benefits of blocked over interleaved learning (Flesch et al., 2018). Finally, the presence of semantic representations within the episodic memory system explains why medial temporal lobe damage can impair not only the acquisition of new semantic knowledge but also the retrieval of previously consolidated semantic information (Duff et al., 2020).

## Discussion

We have developed a model of episodic-semantic memory interactions based on two complementary principles. First, that the algorithm underlying semantic consolidation can be viewed as a form of dictionary learning over episodic experience. Second, that this learned vocabulary can be transferred back into episodic memory, providing both memory systems with a shared semantic vocabulary. We have proposed a computational circuit of the model, where the interactions of episodic and semantic memory are implemented via interactions of a Medial Temporal Lobe (MTL) and a neocortex (CTX) network. The circuit, based on Hebbian local plasticity, uses episodic replay as the main mechanism to perform dictionary learning over compositional sensory input from MTL to CTX. Similarly, spontaneous cortical activity enables a transfer of CTX representations into MTL, shaping the episodic memory code as proposed by our model. The same circuit model provides mechanistic insights into the role of semantic representations in episodic memory, suggesting that they act as an error-correcting scaffold that improves recall and stabilizes further consolidation dynamics as long as new episodes remain semantically in-distribution.

A first contribution of this work is to recast semantic consolidation as a form of dictionary learning. Classical consolidation theories propose that replay enables the gradual emergence of semantic knowledge from episodic experience (McClelland et al., 1995; O’Reilly et al., 2014), with many computational implementations relying on supervised or error-driven learning mechanisms to achieve this transformation (Saxe et al., 2019; Singh et al., 2022; W. Sun et al., 2023; Spens & Burgess, 2024b). In contrast, our framework suggests that replay may provide candidate structure from which neocortex discovers a set of reusable representations capable of explaining recurring patterns across experiences. This is consistent with growing evidence that replay is selective, reconstructive, and generative rather than a faithful reinstantiation of prior activity (Swanson et al., 2020; Diekmann & Cheng, 2023). More broadly, this perspective provides a unified account of two complementary functions often attributed to systems consolidation: the long-term stabilization of individual episodic memories and the extraction of abstract knowledge across experiences. Existing computational work has largely treated these objectives separately, with high-level models focusing on semantic abstraction and generalization, and biologically grounded circuit models primarily addressing the stabilization of individual memories, often inspired by contextual fear conditioning paradigms. Our results suggest that these could be different outcomes of the same replay-driven learning process, depending on the episodic complexity of the word (or *atom*) formed. This could account for personal semantics (Renoult et al., 2012), when the concept formed corresponds to an intermediate level of complexity (e.g. the concept of someone’s workspace), supporting a continuum of consolidation, from purely episodic to highly abstract.

A second central finding of this work is a mechanism for transferring abstract representations from one sub-network to another. In the model, spontaneous activity in neocortex shapes medial temporal representations, increasing their level of abstraction during episodic encoding. The idea that neocortex can transfer semantic representations to subcortical areas has inspired several computational models in recent years. For example, cortex-initiated generative replay has been proposed to align cortical and hippocampal representations (Van de Ven et al., 2020), offering an alternative explanation for how MTL representations keep track of cortical representational shift (Káli & Dayan, 2004). Similarly, this assumption underlies models in which cortical-extracted regularities can be used for episodic compression (Spens & Burgess, 2024a; Nagy et al., 2025) and by the episodic pattern completion machinery (Whittington et al., 2020; Kang & Toyoizumi, 2023; Spens & Burgess, 2024b; Kang & Toyoizumi, 2024). All of these proposals have in turn been inspired by experimental findings supporting the presence of semantic representations in the episodic memory circuit (Quiroga et al., 2005; Quiroga, 2012; Aronov et al., 2017; C. Sun et al., 2020). While previous work has proposed that semantic representations are first extracted by the hippocampal circuit, and then used by neocortex (Gluck & Myers, 1993), converging lines of evidence suggest neocortex might actively influence hippocampal replay and subsequent consolidation (Remondes & Schuman, 2004; Rothschild et al., 2017; de Sousa et al., 2026). Recent work also points in this direction (Wang et al., 2026), showing that shortly after learning the temporal structure of ripples changes from hippocampus leading (here called episodic replay) to neocortex leading (here called semantic replay), in agreement with our model’s prediction. However, a biologically plausible circuit model of this process has yet to be proposed. While still hypothetical, our results provide a mechanistic account of the formation of semantic representations in episodic memory relying only on local Hebbian plasticity. Furthermore, the model’s mechanism is consistent with experimental findings in which neocortical recurrent engrams serve as a substrate for further synaptic potentiation of downstream corticofugal connections (Lee et al., 2023), in particular from prefrontal cortex (PFC) to basolateral amygdala (BLA), which we reproduce here.

Beyond providing a mechanistic account of how semantic representations emerge and become incorporated into episodic memory, our results suggest a normative role for semantic structure within episodic codes. Classical theories of associative memory have long emphasized the importance of sparse and decorrelated representations for maximizing storage capacity (Hopfield, 1982; Tsodyks & Feigel’man, 1988; Treves & Rolls, 1991). From this perspective, semantic overlap between memories would be expected to introduce interference and impair retrieval. Far from being detrimental, semantic representations can in fact benefit episodic memory (Murray et al., 1976; Tse et al., 2007; Lin et al., 2021). Our results offer one possible explanation for this apparent tension. By providing abstract representations that capture regularities across experiences, semantic concepts act as an error-correcting scaffold during memory retrieval, allowing partially degraded episodic patterns to be reconstructed more accurately. Importantly, this benefit extends beyond recall itself. Because replay relies on the reactivation of stored episodic representations, improved attractor dynamics also lead to more reliable replay events and an increased signal-to-noise ratio during consolidation. In this view, semantic representations do not merely coexist with episodic memories but actively improve their stability, retrieval, and communication with neocortex. More broadly, these findings suggest that semantization may represent a computational trade-off in which a modest reduction in memory specificity is compensated by substantial gains in robustness, generalization, and consolidation efficiency, provided that new experiences remain semantically in-distribution.

Taken together, our results aim to reconcile the generally assumed one-way consolidation of circuit models of systems consolidation (O’Reilly et al., 2014; Kumaran et al., 2016) with a rich body of experiments demonstrating that semantic knowledge shapes episodic memory. These include reconstructive and schema-dependent recall (Bartlett, 1932), false memories for semantically related items in the DRM paradigm (Deese, 1959; Roediger & McDermott, 1995), expertise-dependent memory (Chase & Simon, 1973), and improved recall for semantically congruent episodes (Zöllner et al., 2023). Recent computational models have successfully captured several of these phenomena through semantic compression (Nagy et al., 2020) or semantic completion of partial episodic traces (Fayyaz et al., 2022). In contrast, our work proposes an explicit circuit mechanism based on replay, local plasticity, and bidirectional consolidation, providing a biologically grounded framework in which these semantic effects can emerge. While the specific mechanisms remain to be experimentally validated, we hope this model provides a useful circuit-level scaffold for generating mechanistic hypotheses that bridge psychological phenomena and systems-level learning theories.

Our framework generates several experimentally testable predictions. In an effort to map our episode generation protocol to experiments in rodents, we propose a paradigm for compositional cue (*A*_*i*_ visual, *B*_*j*_ odour)-reward associations (Figure S8). We propose one prediction for each of the three key mechanisms studied here. First, if dictionary learning is the substrate for semantic learning, we anticipate that the number of cells tuned to the different components *A*_*i*_ and *B*_*j*_ will increase monotonically with the marginal probability of that concept being present in any episode (Fig. 2g). A parallel prediction is that holographic manipulation of cortical abstract cells could reflect semantic structure (predictiveness) across concepts, thereby encoding the *world model* in recurrent connections (Fig. 2h). This would support our mechanism for the slow extraction of semantics in CTX. Second, if semantic representations are transferred back into episodic memory through semantic replay, disrupting cortical (with prefrontal cortex being a plausible candidate) activity during consolidation should reduce the abstractness (Bernardi et al., 2020) of medial temporal lobe representations regarding the different factor values (concepts) presented to the animal. Finally, the model predicts that memories composed of previously learned concepts should consolidate more rapidly than entirely novel experiences. Behaviourally, this would imply that learning the reward association to a novel visual-odour (*A*_*i*_, *B*_*j*_) cue would be faster when each individual cue has previously been learned than when the cues are novel (Fig. 6d).

Several limitations of the present work point to promising directions for future research. First, the model intentionally abstracts away much of the anatomical and physiological complexity of hippocampal-neocortical interactions. While this simplification allows us to isolate the main underlying computational principles in a tractable manner, future work could investigate how these mechanisms interact, for example, with the rich circuitry of the medial temporal lobe, including the distinct contributions of entorhinal cortex, hippocampal subregions, or amygdala. Second, our implementation focuses on static representations and does not explicitly address the temporal structure of episodic memories. Extending the framework to sequences and temporally extended experiences is crucial to understand how the abstraction of dynamically changing environments is implemented by the brain. Third, while we study a regime without imposed pattern separation in the hippocampal code (W. Sun et al., 2023; Chandra et al., 2025), it is likely that memories involve both structured overlapping representations and pattern-separated hashing codes (Chettih et al., 2024). The computational mechanisms supporting the flexible use of both during encoding and retrieval have yet to be established. Finally, from a theory of learning perspective, the model could serve as an inspiration for brain-inspired machine learning algorithms, in which semantic and episodic memory conceptualizations are becoming increasingly relevant (Fountas et al., 2025; Dong et al., 2025; Pink et al., 2025).

We hope that the present framework provides a foundation for investigating cortical and subcortical circuits as cooperative components that play a crucial role in the brain’s ability to simultaneously generalize and memorize. In this light, our work suggests that episodic and semantic memory may be best understood as two interacting representational systems that continuously learn models of one another.

## Methods

### Episode Generation Protocol

The Episode Generation Protocol (EGP) is a generative process that samples neuronal activity in the sensory layer. Our proposed EGP follows a similar approach to that described in Albesa-González and Clopath (2025), but is outlined here again for completeness:

### Episodes, latent factors, and concepts

All *episodes* share a structure, such that every episode contains one *episode concept* (value) per episode *latent factor*. This means that, given latent factors *A* and *B*, with each latent factor representing a set of episode concepts *A*_*i*_ ∈ *A, B*_*j*_ ∈ *B*, then the set of all possible episodes is defined as:

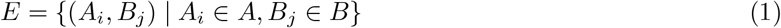

In other words, an episode *e* ∈ *E* is an ordered set of episode concepts *A*_*i*_, *B*_*j*_ such that *A*_*i*_ belongs to latent factor *A* and element *B*_*j*_ belongs to latent factor *B*. In this study we have fixed *A* to have 5 possible values *A*_*i*_ ∈{*A*_1_, *A*_2_, *A*_3_, *A*_4_, *A*_5_ } and likewise for *B*_*j*_ ∈{*B*_1_, *B*_2_, *B*_3_, *B*_4_, *B*_5_}. However, the model is robust against variations in the number of concepts per latent factor (Fig. S9d).

### Episode Probability Distribution

As a generative process, our EGP samples episodes as a preliminary step to sampling sensory input. We do this by fixing a probability distribution over episodes:

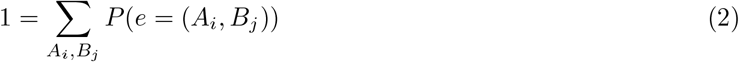

such that *P* (*e* = (*A*_*i*_, *B*_*j*_)) ≡*P* (*i, j*) defines the probability of an episode with factor values (*A*_*i*_, *B*_*j*_) to be sampled when generating episodes.

### Episode-Input Mapping

Ultimately, the EGP samples sensory layer activity. For this reason, one also has to define how a sampled episode is mapped into sensory input *X*^SEN^. Every concept *c* in *A* ∪*B* has a set of associated neurons SEN_*c*_ in *X*^SEN^ such that, given an episode *e*,

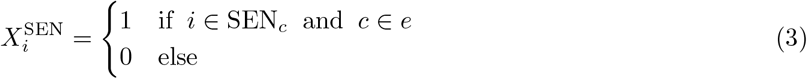

In order to define SEN_*c*_, we specify the number of latent factors *N*_latent-factors_, in this case 2 (*A* and *B*). Then, we define the number of concepts (possible values) per latent factor (|*A*| and |*B*|, here both 5). Thus, and *N A*_*i∈*[1,5]_ and *B*_*j∈*[1,5]_. Then, we fix the number of neurons per concept 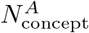, and 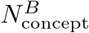 define SEN_*c*_ as:

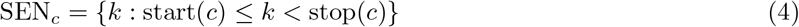

with 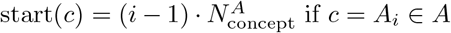 and 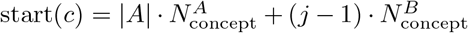 if *c* = *B*_*j*_ ∈ *B*, if and similarly 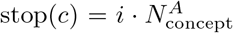 if *c* = *A*_*i*_ ∈ *A* and 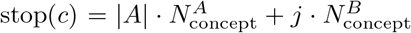. Intuitively, this can be thought of as a compositional extension of the standard orthogonal split of neuronal assemblies, with two neuronal pools of orthogonal representations being concatenated.

To account for variability between different presentations of the same episode (not every presentation of the same episode, even though it contains the same concepts, will be exactly the same), we further add some stochasticity by randomly picking *N*_swap_ inactive neurons and *N*_swap_ active neurons and inverting their activity (a total of 2*N*_swap_ neurons randomly change their activity). This ensures that activity sparsity in the sensory layer is maintained (the number of swaps from 0 to 1 is the same as the number of swaps from 1 to 0). An illustration of the episode-input mapping is included in the supplementary material (S2).

#### Input temporal structure

Generated input *X*_input_ has dimensions (*N*_days_, *T*_day_, *N*_SEN_). Every day, a total of *T*_day_*/T*_episode_ episodes (of *T*_episode_ length) are sampled, and within each timestep an independent noise realization is sampled.

#### Semantic Structure of an EGP

Following Albesa-González and Clopath (2025), we refer to *semantic field theory* (Bussmann et al., 2006) in order to define what is the *semantic structure* of an EGP. According to this theory, the meaning of a word is not isolated but dependent on its relation to the rest of the words. While our task is not one of language, we can use this same paradigm to define the *meaning* of episode concepts. In this sense, the meaning (semantics) of our episode concepts depends on how they are related to the rest (also note this is how semantics are established in a language dictionary, where each concept is defined via its relationship with the rest of the concepts).

Following this proposal, we use the conditional probabilities of being present in an episode between episode concepts as a proxy for the semantic structure of an EGP. In other words, extracting the semantics of an EGP is equivalent to (i) identifying the underlying concepts, and (ii) extracting how likely one concept *c*_*i*_ is to be present in an episode *e* if a concept *c*_*j*_ is also present in *e*. We define the Semantic Structure SS of an EGP with *N* ^concepts^ different concepts *c*_*i*_, as a *N* ^concepts^ × *N* ^concepts^ matrix of the form:

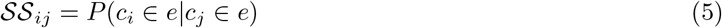

For example, the intuition that pizza has a stronger semantic connection to cheese than cheese does to pizza is reflected in their conditional probabilities (it is very common to have cheese in a pizza, but much less to have pizza in any dish containing cheese).

### Circuit Model

#### Architecture and nomenclature

##### Regions and neural activity

The circuit model contains 3 main regions: sensory cortex (SEN), neocortex (CTX), and medial temporal lobe (MTL). In turn, MTL is split into two sub-regions: MTL-sensory and MTL-semantic. CTX is likewise partitioned into two subregions CTX-simple and CTX-complex, which differ in number of neurons and neuronal excitability, and learn concepts at different levels of abstraction. We use *X* to denote the neural activity, which is written as *X*^region^ (for example *X*^CTX^) when the region is to be made explicit. The same follows for pre-activation (synaptic) input, which is denoted by 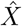 (or 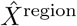). The activity of neuron *i* in a region is 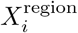. The number of neurons in a region is denoted by *N*_region_ (e.g. *N*_CTX_).

##### Connectivity

Connectivity matrices connecting a region pre to a region post take the form *W* ^post*←*pre^, and a synapse connecting neuron *j* in region pre with neuron *i* in region post is 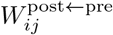. For example, 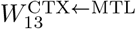 represents the connection from neuron 3 in MTL to neuron 1 in CTX. Connectivity is classified as either feed-forward (connecting two different regions), which includes *W* ^MTL-sensory*←*SEN^, *W* ^CTX*←*MTL^, and *W* ^MTL-semantic*←*CTX^, or recurrent (connecting a region with itself), which includes *W* ^CTX*←*CTX^, and *W* ^MTL*←*MTL^. There is only one set of *W* ^CTX*←*MTL^ connections. When an algorithm specifies *W* ^CTX*←*MTL-sensory^ that means taking only the columns that correspond to MTL-sensory neurons. All connections are plastic except *W* ^MTL-sensory*←*SEN^, which is fixed. Given a connectivity matrix *W* ^post*←*pre^, we call the receptive field of a postsynaptic neuron 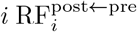 the vector representing the connections from region pre to postsynaptic neuron *i*

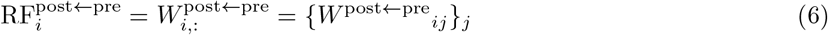

#### Synaptic Initialization

##### Sensory cortex to MTL-sensory

*W* ^MTL-sensory*←*SEN^ is initialized as the identity, mapping to MTL-sensory an exact copy of *X*^SEN^.

##### Feed-Forward connections

The rest of feed-forward connections are sampled from a Gaussian distribution with mean 0 and standard deviation *W* ^post*←*pre^ as a network parameter.

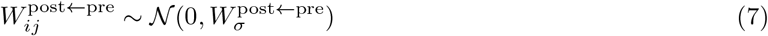

##### Recurrent connections

Recurrent connections are initialized at 0. MTL connections are reset every *day* (see Algorithm 7)

##### Mature and immature neurons

CTX and MTL-semantic neurons (those with synapses that perform statistical learning) in the model can be either *mature* or *immature*. The intuition is: do these neurons contain meaningful representations via input received from another region (mature) or are mostly nonselective and random (immature)? More details on the dynamics and plasticity of mature and immature neurons can be seen below, but to summarize: (i) immature neurons have higher excitability during feedforward input processing (not during pattern completion) and (ii) immature neurons have a higher learning rate during sleep. Consequently, during replay, immature postsynaptic neurons tend to be recruited when the presynaptic pattern is poorly matched by existing receptive fields. By doing this, if a presynaptic pattern is similar to previously formed representations, it slightly changes the already formed representations. If not, a set of immature neurons win the competition, develop a receptive field due to their higher plasticity, and then become mature. This allows one-shot formation of new postsynaptic representations of presynaptic input when an input pattern is too different from those presented to the network before, and then refining these representations with a slower learning rate. CTX (outgoing) recurrent connections of immature neurons also benefit from higher plasticity rates during sleep. Maturity in a region is denoted by IM^region^, where 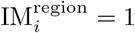 if neuron *i* in the region is immature, and 0 otherwise.

#### Outgoing and incoming homeostasis across types of connections

Below, we formalize two regimes of homeostatic plasticity: outgoing homeostasis and incoming homeostasis. Outgoing homeostasis guarantees that the sum of the outgoing connections from neuron *j* in a connectivity matrix *W* does not exceed a certain 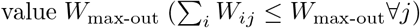. Similarly, incoming homeostasis guarantees a bound in incoming connections 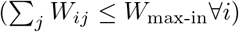. Depending on the type of connectivity we choose:

- outgoing homeostasis for CTX ←CTX: this makes the connectivity matrix reflect semantic structure as defined in (5), as the connection from neuron coding for concept *j* to neuron coding for concept *i* becomes proportional to the conditional probability of neuron *i* firing given neuron *j* fires (AlbesaGonzález & Clopath, 2025)
- incoming homeostasis for feed-forward connectivity: in this homeostatic regime, connection from neuron *j* to neuron *i* becomes proportional to *p*(*j* active|*i* active) (Albesa-González & Clopath, 2025). The *trick* is that if an initial one-shot receptive field makes neuron *i* selective to concept *c*_*i*_, receptive fields approximate the average presynaptic activity pattern (prototype) given the concept *c*_*i*_ is present (neuron *i* is active), thus reinforcing higher selectivity in noisier regimes.
- Homeostatic reset in MTL ← MTL connections. In order to account for symmetric plasticity kernels in hippocampus (Mishra et al., 2016; Li et al., 2023), we use standard Hebbian learning in MTL ←MTL. Homeostasis is obtained by resetting MTL recurrent connections to 0 at the start of *wake* (Algorithm 7).

### Network Operations

#### Neuronal activation

Synaptic input is mapped to neural activity via a *K-winners-take-all* mechanism, in which the *K* neurons with the highest pre-activation input 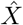 set their activity to 1, and the rest to 0. This can be written as

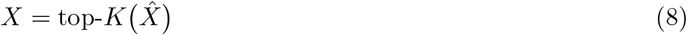

In order to determine *K* in a particular region (or subregion), first we define

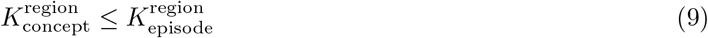

such that each concept is represented by 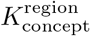 neurons, and a typical episode during wakefulness contains 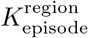 active neurons. In this context, given *K* active neurons in a region, we define the semantic load *S*_*L*_ to be the number of concepts represented by *K* neurons, such that

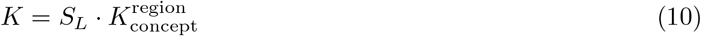

In other words, by fixing the semantic load, one fixes how many neurons in total are active, making *K* higher if more concepts are being represented. The maximum number of concepts that can be encoded in an episode is thus:

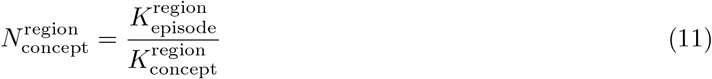

Therefore, we can define Activate (Algorithm 1) as an algorithm that maps presynaptic input 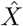, given a semantic load *S*_*L*_, into neural activation *X* in the following way:

##### Algorithm 1 Neuronal Activation

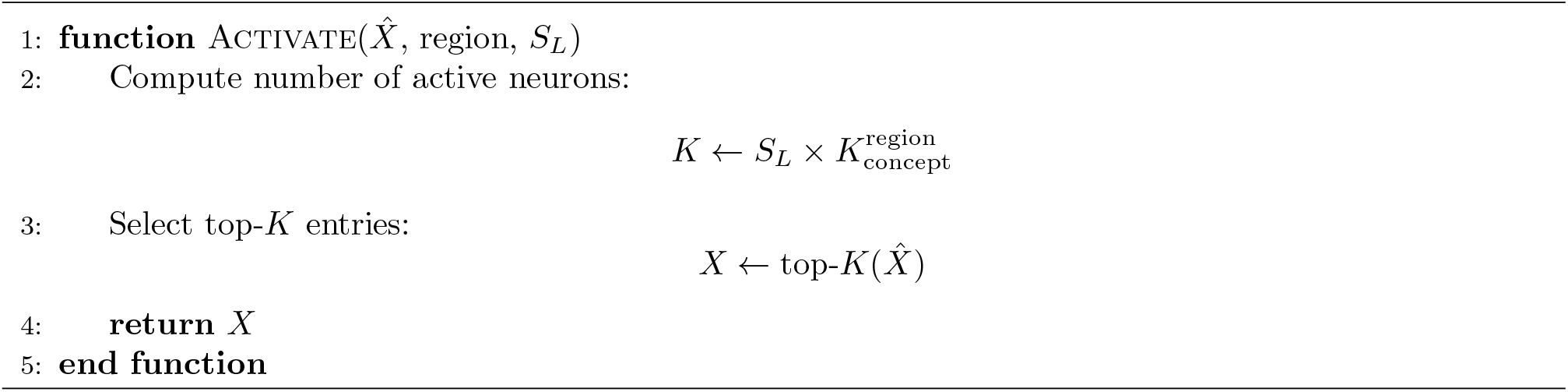

#### Feed-forward input processing

Input processing is computed for a single pre and post region at a time, with pre-activation input 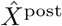 obtained via

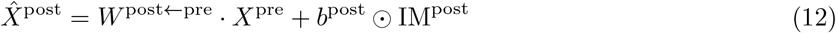

*b*^post^ is the extra excitability that immature neurons have in region post.

#### Pattern completion (recurrent input processing)

Given a recurrent network, pattern completion is typically defined as an iterative update on neural activity *X*^region^ that is initiated in an initial state 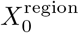:

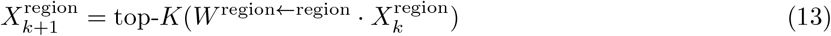

with *W* ^region*←*region^ the region recurrent connectivity matrix, *K* dependent on the specified semantic load *S*_*L*_, and *k* a timestep operating on a smaller timescale, than the overall timescale *t*. This means that within a timestep *t*, there are pattern complete iterations iterations on temporal index *k*. This is used to define PatternComplete as Algorithm 2:

##### Algorithm 2 Pattern Completion

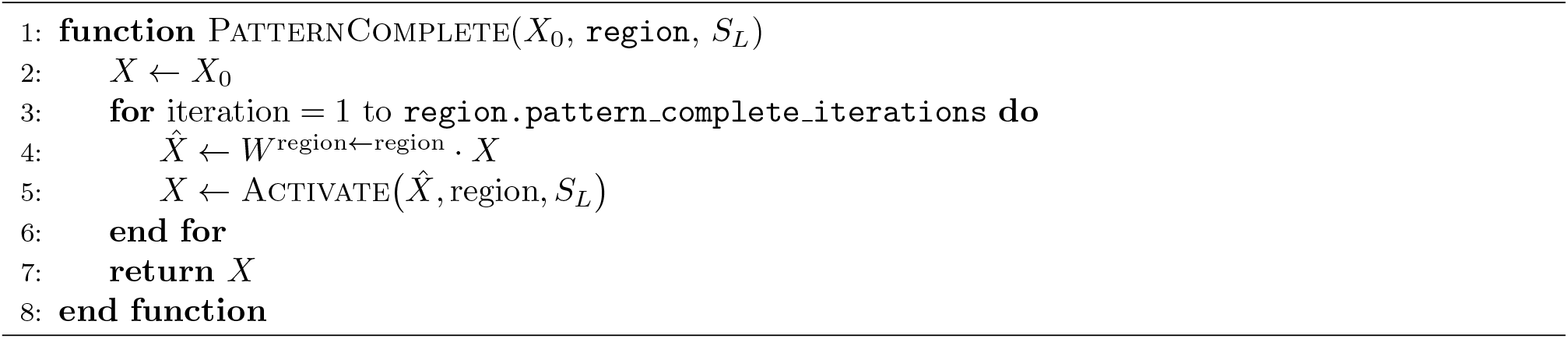

#### Hebbian Learning

The connection between a presynaptic neuron *j* and a postsynaptic neuron *i* subject to Hebbian learning is updated as:

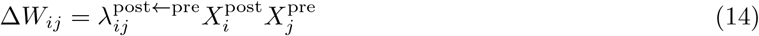

for a presynaptic region pre and a postsynaptic region post, where 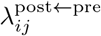 is the learning rate from neuron *j* in region pre to neuron *i* in region post.

##### Maturity-dependent learning rate

In order to combine one-shot and statistical learning (see AlbesaGonzález and Clopath (2025)), the learning rate of connections depends on the *maturity* of neurons, which is given by IM. Neurons start in an *immature* state IM_*i*_ = 1, allowing them to form initial *connectivity estimates*. Once this initial connectivity is formed, the learning rate is reduced to capture the statistical regularities of the neural activity patterns (the neuron becomes *mature*, IM_*i*_ = 0). To model this, we define a quick learning rate

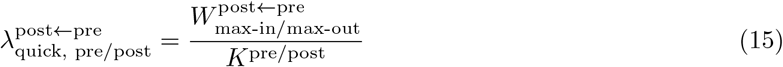

where again 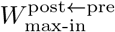 (or 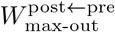) is a network parameter determining the maximum sum of incoming (or outgoing) connections the matrix *W* ^post*←*pre^ has for every postsynaptic (or presynaptic) neuron. Then, at each Hebbian update, we have for incoming-bounded connections:

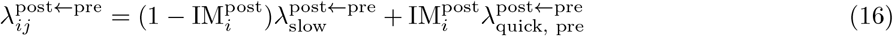

and for outgoing-bounded connections:

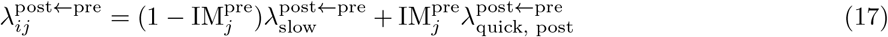

where 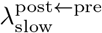 is a network parameter indicating the slow learning rate (used throughout all simulations except for the first plasticity event in which the neuron transitions from immature to mature).

Note how the quick learning rate (Eq. (15)) is defined such that when the neuron transitions from immature to mature, with the *first shot of learning*, the condition 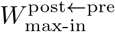, for incoming normalized weights, is met:

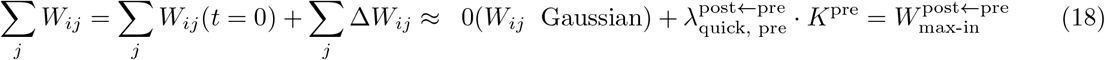

(and similarly for outgoing normalized weights). Also note how for incoming-bounded connections, as are feed-forward synapses, this makes the formed receptive field proportional to *X*^pre^ (making it maximally selective to it). For outgoing-bounded connections, here recurrent cortical synapses, this is equivalent to making *X*^post^ the initial guess in meaning of neuron *j*.

##### Hebbian learning algorithm

Altogether, this defines Hebbian (Algorithm 3) as the following update in *W* ^post*←*pre^:

#### Homeostatic Plasticity

We use multiplicative normalization that guarantees, for every weight matrix *W* ^post*←*pre^, the total sum of incoming or outgoing connections is capped to a certain value. Following Albesa-González and Clopath (2025), we use *incoming homeostasis* as:

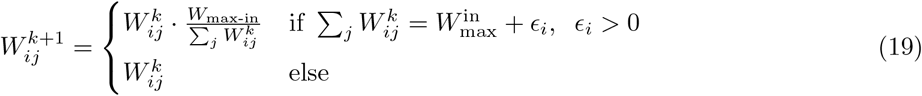

where 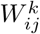 denotes the value of synapse *ij* before applying homeostasis and 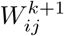 the value after applying homeostasis (for readability, we simply write *W* ≡ *W* ^post*←*pre^). Similarly, we have *outgoing homeostasis* to be defined by:

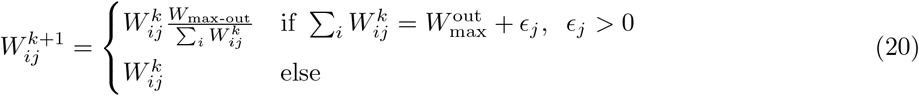

##### Algorithm 3 Hebbian Learning

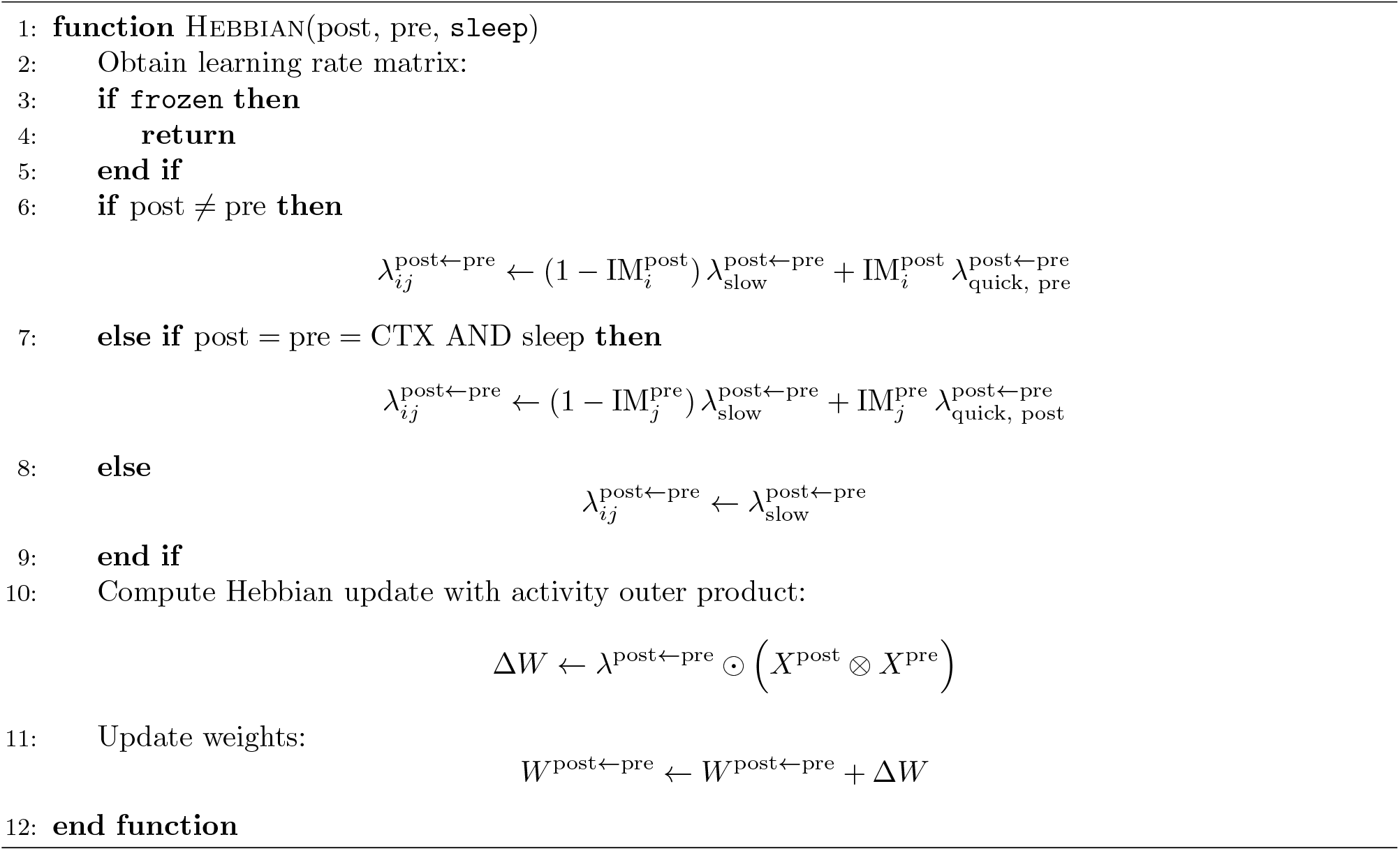

This allows, for a presynaptic region pre and a postsynaptic region post, defining Homeostasis as Algorithm 4.

#### Fixed points of synaptic dynamics

The combination of *K-winners-take-all* (top-*K*) activation function with Hebbian and homeostatic plasticity as defined here has been recently shown (Albesa-González & Clopath, 2025) to make synaptic connections converge to conditional firing probabilities. For outgoing homeostasis, the fixed points are:

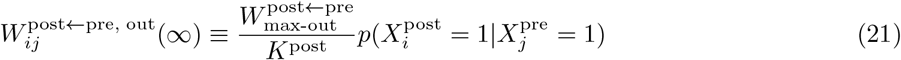

where *K*^post^ corresponds to the number of active neurons during plasticity. Given that the conditional probability can be at most 1, we define in this scenario 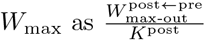 . Similarly, for incoming homeostasis, the fixed points are

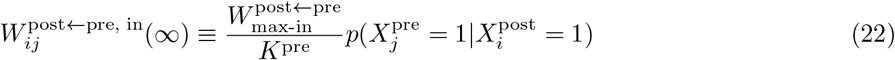

with 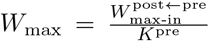 . These results have an important associated intuition, which is that if neurons code for individual concept (have a concept selectivity close to 1), synaptic connections reflect the associated Semantic Structure (Albesa-González & Clopath, 2025) under outgoing homeostasis (as happens in ←CTX). In MTL-semantic CTX and CTX ←MTL, which follow incoming homeostasis, when initial one-shot receptive fields induce a selectivity to a concept close to 1, feed-forward synapses slowly approximate to the associated prototype, where each synapse is becomes proportional to the contribution of each presynaptic neuron to the concept encoded in the postsynaptic neuron.

#### Replay

We define Replay as a learning mechanism involving a pre and a post region, such that *W* ^post*←*pre^ extract statistical structure from *W* ^pre*←*pre^ connections. A replay operation takes as argument a specific semantic load *S*_*L*_, which determines the conceptual complexity of the extracted regularities. To do this, we first pattern complete an initial presynaptic pattern 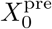, defined as a Gaussian random vector of the same size as the pre region:

##### Algorithm 4 Homeostasis

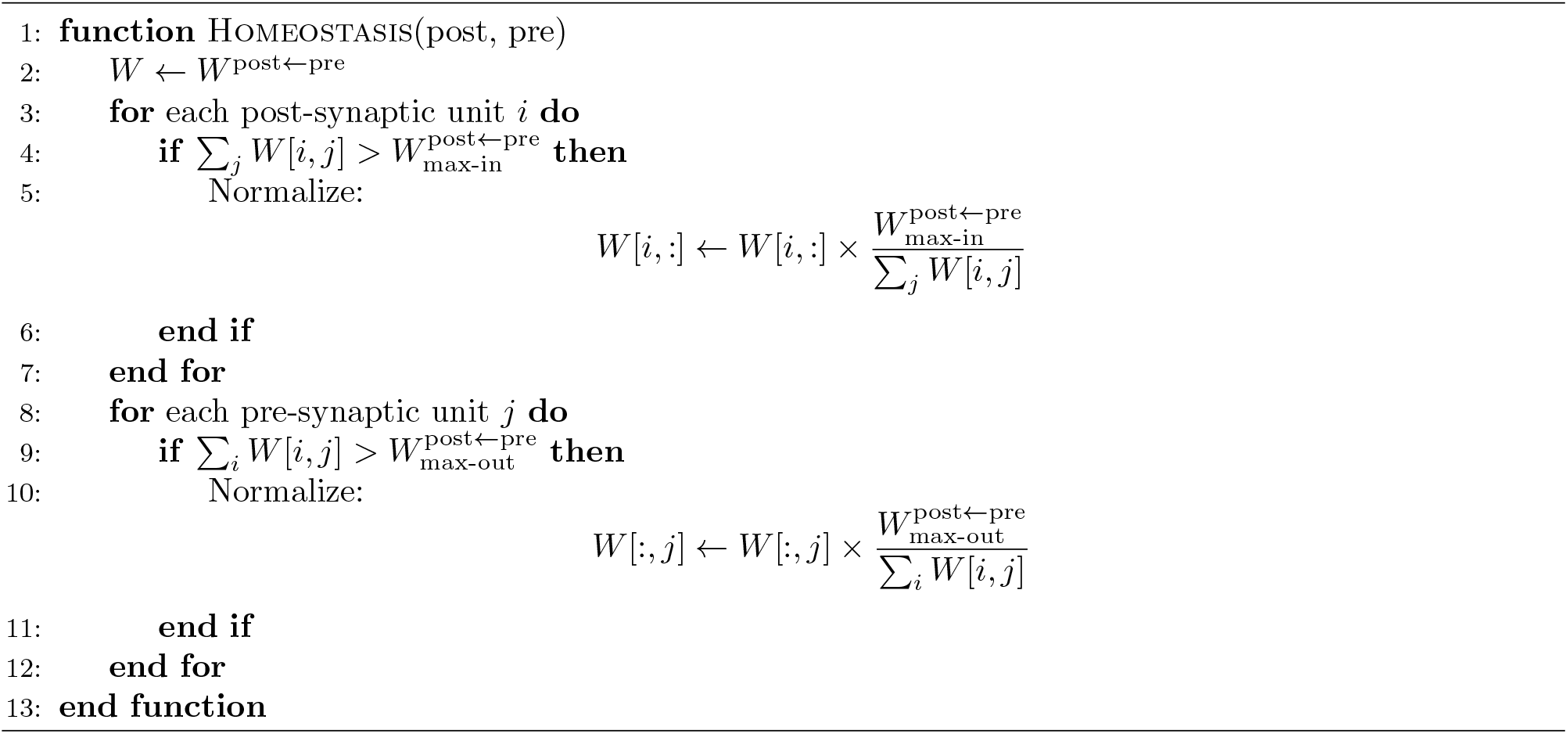

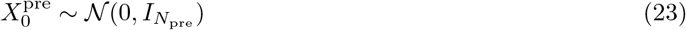

where *N*_pre_ is the size of region pre and (0,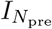) is a multivariate Gaussian distribution with 0 mean and identity covariance. This initial random pattern is pattern completed to obtain *X*^pre^, and then projected to the post region, to give *X*^post^, using in both operations the same fixed semantic load. Then, Hebbian and homeostatic plasticity operations as defined in Algorithms 3 and 4 are used in the (post, pre) region pair. Replay is computed in simulations following Algorithm 5. In the text, we call replay from MTL to CTX *Episodic Replay*, because it finds regularities across recurrent synapses that contain episodic information, and replay from CTX to MTL-semantic *Semantic Replay*, because in this case recurrent synapses contain semantic information.

##### Algorithm 5 Replay

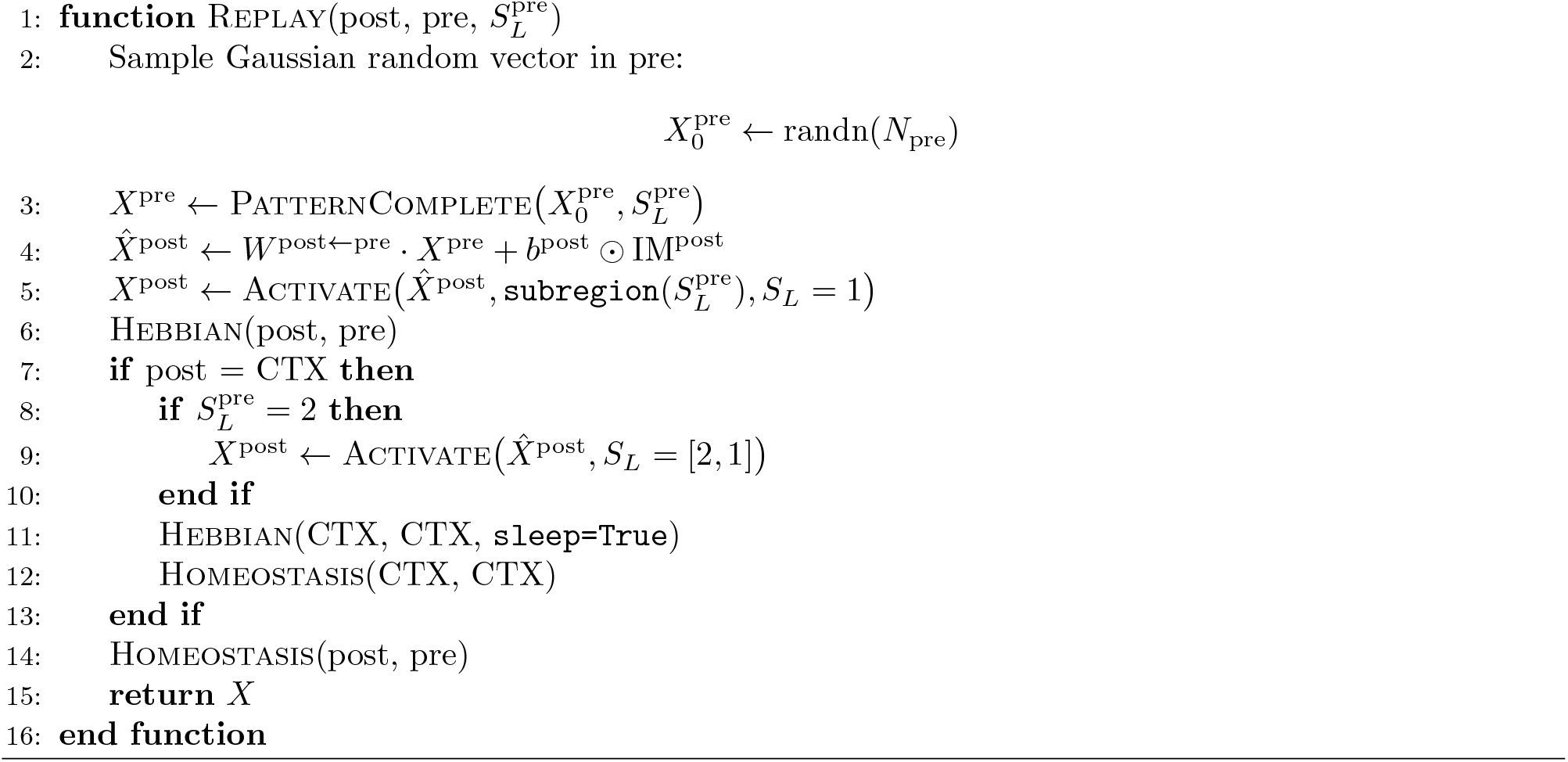

### Learning Algorithm

For each day of experience, assuming an input *X*_input_ of shape (*N*_days_, *T*_day_, *N*_SEN_), we define the following learning algorithm Train (Algorithm 6), where Wake is defined as in Algorithm 7 and the Sleep as in Algorithm 8. When it is mentioned that the network is *tested*, or *frozen*, that in practice means that Train (Algorithm 6) omits Sleep, with episodic memory being actually intact (but no consolidation or long-term learning takes place). All operations used have been previously defined, with these algorithms showing the order in which they are performed. duration phase A represents the number of days in which only episodic replay takes place (Fig. 2), and duration phase B the number of days in which only MTL-sensory synapses are used for episodic replay, but semantic replay starts taking place (Fig. 3). From day duration phase B on, MTL replays both sensory and semantic representations (Starting with Fig. S5). Therefore, during duration phase A days, episodic memories containing only sensory information are stored in MTL during Wake (Algorithm 7). During Sleep (Algorithm 8), cortical abstractions of sensory information are formed. After duration phase A days, cortical abstractions are transferred to MTL-semantic via semantic replay during Sleep. Finally, after duration phase B days, episodic memories contain both sensory and semantic information, and so does episodic replay. This, together with a variable semantic load during replay, facilitates the formation of new semantic representations that are based on previous abstractions.

#### Algorithm 6 Learning Algorithm Across Days

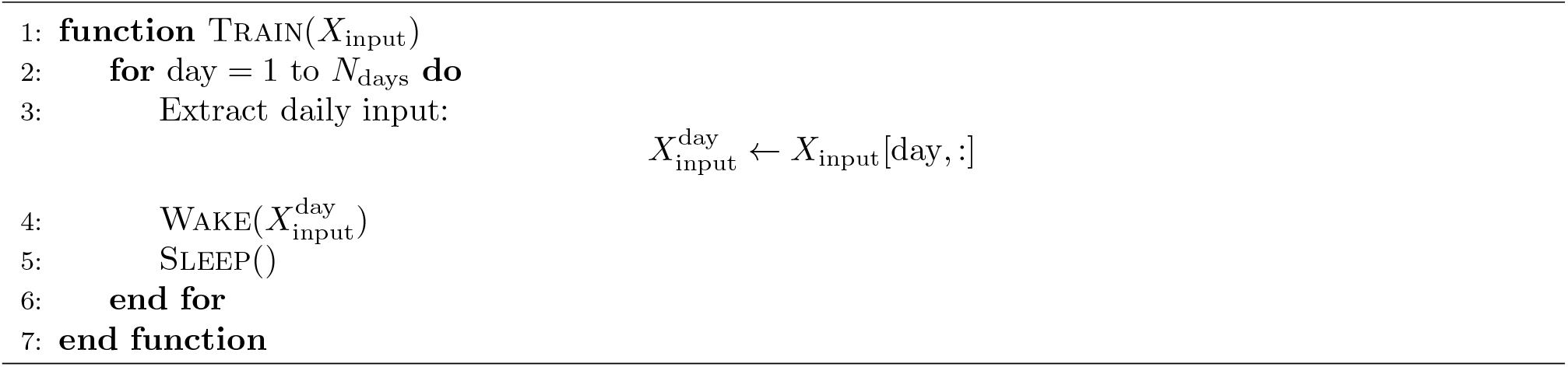

### Default Simulation parameters

Default input, temporal, and model parameters can be found in Tables 1-3. Figure-specific parameters are discussed in the following sections.

**Table 1:** Summary of Episode Generation Protocol parameters.

| Variable | Description | Value |
| --- | --- | --- |
| $N_{\text{latent-factors}}$ | Number of episode latent factors | 2 |
| $\vert A\vert , \vert B\vert $ | Number of factor values (concepts) per latent factor | [5, 5] |
| $N_{\text{concept}}^A, N_{\text{concept}}^B$ | Number of active neurons per concept | [10, 10] |
| $P(A_i, B_j)$ | Episode probability | 0.5/5 if $i = j$ ; 0.5/20 if $i \neq j$ |
| $N_{\text{swaps}}$ | Number of swaps (total noise is $2N_{\text{swaps}}$) | 4 |

**Table 2:**
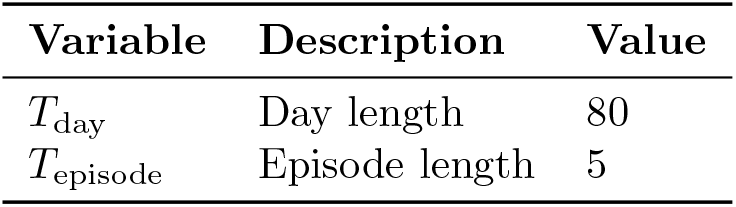
Summary of default input temporal parameters.

**Table 3:**
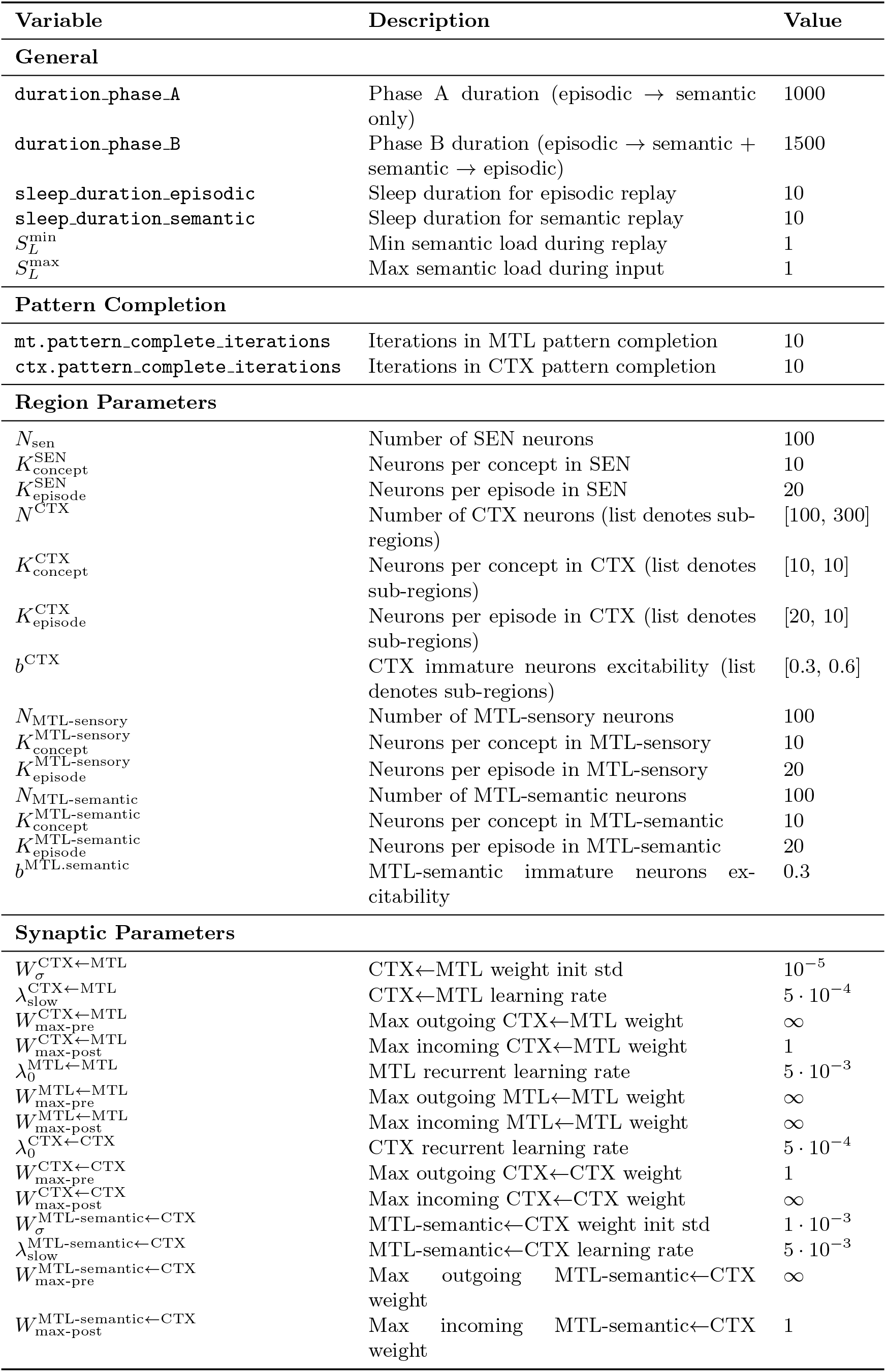
Summary of network parameters used in the simulations (default unless specified otherwise).

**Table 4:** Default data-analysis parameters. Figure-specific tables report deviations from these values.

| Variable | Description | Value |
| --- | --- | --- |
| $K_{\text{assembly}}$ | Neurons assigned to each concept-specific assembly | 10 |
| $T_{\text{NMI}}$ | NMI trajectory window size | 160 timesteps |
| $\Delta T_{\text{NMI}}$ | NMI trajectory stride | 80 timesteps |

**Table 5:** Figure 2 parameters (different than or not specified in default)

| Variable | Description | Value |
| --- | --- | --- |
| Example network used in Figs. 2b–h |  |  |
| $N_{\text{days}}^{\text{warmup}}$ | Wake-only warm-up days | 100 |
| $N_{\text{days}}^{\text{train}}$ | Phase A training days | 1000 |
| Fig. 2b |  |  |
| Displayed timesteps |  | First 40 awake |
| Fig. 2c |  |  |
| Wake snapshot |  | day 0, timestep 9 |
| Replay snapshot |  | day 0, timestep 2 |
| Fig. 2d |  |  |
| Example CTX neuron index |  | 30 |
| Fig. 2e |  |  |
| Displayed awake timesteps |  | last 40 awake timesteps |
| Fig. 2f |  |  |
| Window size | Mutual-information window | 160 timesteps |
| Stride | Mutual-information stride | 80 timesteps |
| Fig. 2g |  |  |
| $N_{\text{days}}$ | Wake training days per simulation | 1 |
| Focal probability grid | Sampled marginal probabilities | $\{0.1, 0.2, \dots, 0.9\}$ |
| Dirichlet concentration | Non-focal marginal sampling | 1.0 |
| Seeds | Independent simulations | 1200 |
| $T_A$ | Sleep A durations compared | 1, 10, 30, 100 |
| $T_B$ | Sleep B duration | 0 |

**Table 6:** Figure 3 parameters (different than or not specified in default)

| Variable | Description | Value |
| --- | --- | --- |
| Example network used in Figs. 3b–f |  |  |
| $N_{\text{days}}^{\text{Phase A}}$ | | 1000 |
| $N_{\text{days}}^{\text{Phase B}}$ | | 1500 |
| Fig. 3b |  |  |
| Awake example |  | day 1000 |
| Displayed timesteps |  | first 40 awake timesteps |
| Fig. 3c |  |  |
| Initial CTX state |  | Gaussian |
| Fig. 3d |  |  |
| Input used after learning |  | same MTL-sensory pattern as Fig. 3b |

**Table 7:** Figure 4 parameters (different than or not specified in default)

| Variable | Description | Value |
| --- | --- | --- |
| Example network used in Figs. 4b–e |  |  |
| Initial network |  | checkpoint from Fig. 3 |
| $N_{\text{days}}^{S_L=1}$ | | 100 |
| $N_{\text{days}}^{S_L=2}$ | | 1000 |
| Fig. 4c |  |  |
| Timestep |  | -40 |
| Fig. 4d |  |  |
| Timesteps |  | [-40:] |
| Fig. 4e |  |  |
| $K_{\text{assembly}}^{\text{CTX-simple}}$ | otherwise not enough to build 25 assemblies | 4 |
| Fig. 4f |  |  |
| $N_{\text{days}}$ | Wake training days per simulation | 1 |
| Seeds |  | 1200 |
| Semantic load during sleep |  | 2 |

**Table 8:**
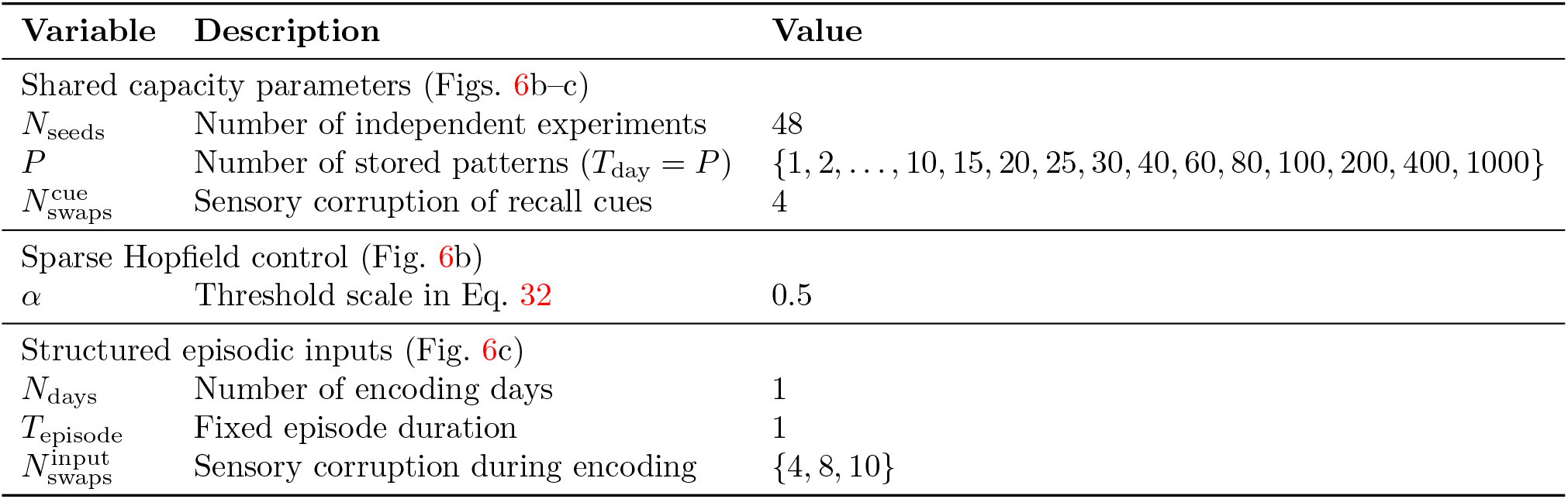
Figure 6 - Recall Benefits parameters (different than or not specified in default)

| Variable | Description | Value |
| --- | --- | --- |
| Shared capacity parameters (Figs. 6b–c) |  |  |
| $N_{\text{seeds}}$ | Number of independent experiments | 48 |
| $P$ | Number of stored patterns ($T_{\text{day}} = P$) | $\{1, 2, \dots, 10, 15, 20, 25, 30, 40, 60, 80, 100, 200, 400, 1000\}$ |
| $N_{\text{swaps}}^{\text{cue}}$ | Sensory corruption of recall cues | 4 |
| Sparse Hopfield control (Fig. 6b) |  |  |
| $\alpha$ | Threshold scale in Eq. 32 | 0.5 |
| Structured episodic inputs (Fig. 6c) |  |  |
| $N_{\text{days}}$ | Number of encoding days | 1 |
| $T_{\text{episode}}$ | Fixed episode duration | 1 |
| $N_{\text{swaps}}^{\text{input}}$ | Sensory corruption during encoding | $\{4, 8, 10\}$ |

**Table 9:** Figure 6 - Consolidation Benefits parameters (different than or not specified in default)

| Variable | Description | Value |
| --- | --- | --- |
| $N_{\text{seeds}}$ | Number of network realizations | 1 |
| $N_{\text{swaps}}^{\text{complex}}$ | Sensory corruption during complex-concept training and evaluation | $\{1, 2, \dots, 15\}$ |
| $T_{\text{day}}$ | Timesteps per wake day | 80 |
| $T_{\text{episode}}$ | Timesteps per episode | 5 |
| $N_{\text{days}}^{\text{Phase A}}$ | Duration of Phase A | 200 days |
| $N_{\text{days}}^{\text{Phase B}}$ | Duration of Phase B | 400 days |
| $N_{\text{days}}^{\text{simple}}$ | Simple-concept training duration | 600 days |
| $N_{\text{swaps}}^{\text{simple}}$ | Sensory corruption during simple-concept training | 4 |
| $N_{\text{days}}^{\text{complex}}$ | Complex-concept training duration | 400 days |
| $N_{\text{days}}^{\text{eval}}$ | Frozen complex-concept evaluation duration | 100 days |
| $N_{\text{RF}}$ | Replayed patterns sampled as receptive fields per complex concept | 10 |

**Table 10:** Figure 5 Concept Cell parameters (different than or not specified in default)

| Variable | Description | Value |
| --- | --- | --- |
| $N_{\text{seeds}}$ | Number of network realizations | 1 |
| $N_{\text{swaps}}$ | Sensory corruption during all concept-cell assays | $\{1, 2, \dots, 15\}$ |
| Cross-context decoding (Fig. 5b1) |  |  |
| $N_{\text{blocks}}^{\text{train}}$ | Number of deterministic training blocks | 2 |
| $T_{\text{episode}}^{\text{train}}$ | Duration of each training block | 1 timestep |
| $T_{\text{day}}^{\text{test}}$ | Duration of the excluded-context test day | 200 timesteps |
| $T_{\text{episode}}^{\text{test}}$ | Duration of each test episode | 5 timesteps |
| $N_{\text{repeats}}$ | Random context-contrast pairs per target concept | 10 |
| Selectivity and mutual information (Figs. 5b2-b3) |  |  |
| $T_{\text{day}}$ | Duration of the frozen evaluation day | 10,000 timesteps |
| $T_{\text{episode}}$ | Duration of each evaluation episode | 5 timesteps |
| $p(A_i, B_j)$ | Probability of each latent pair during evaluation | 1/25 |

#### Algorithm 7 Wake

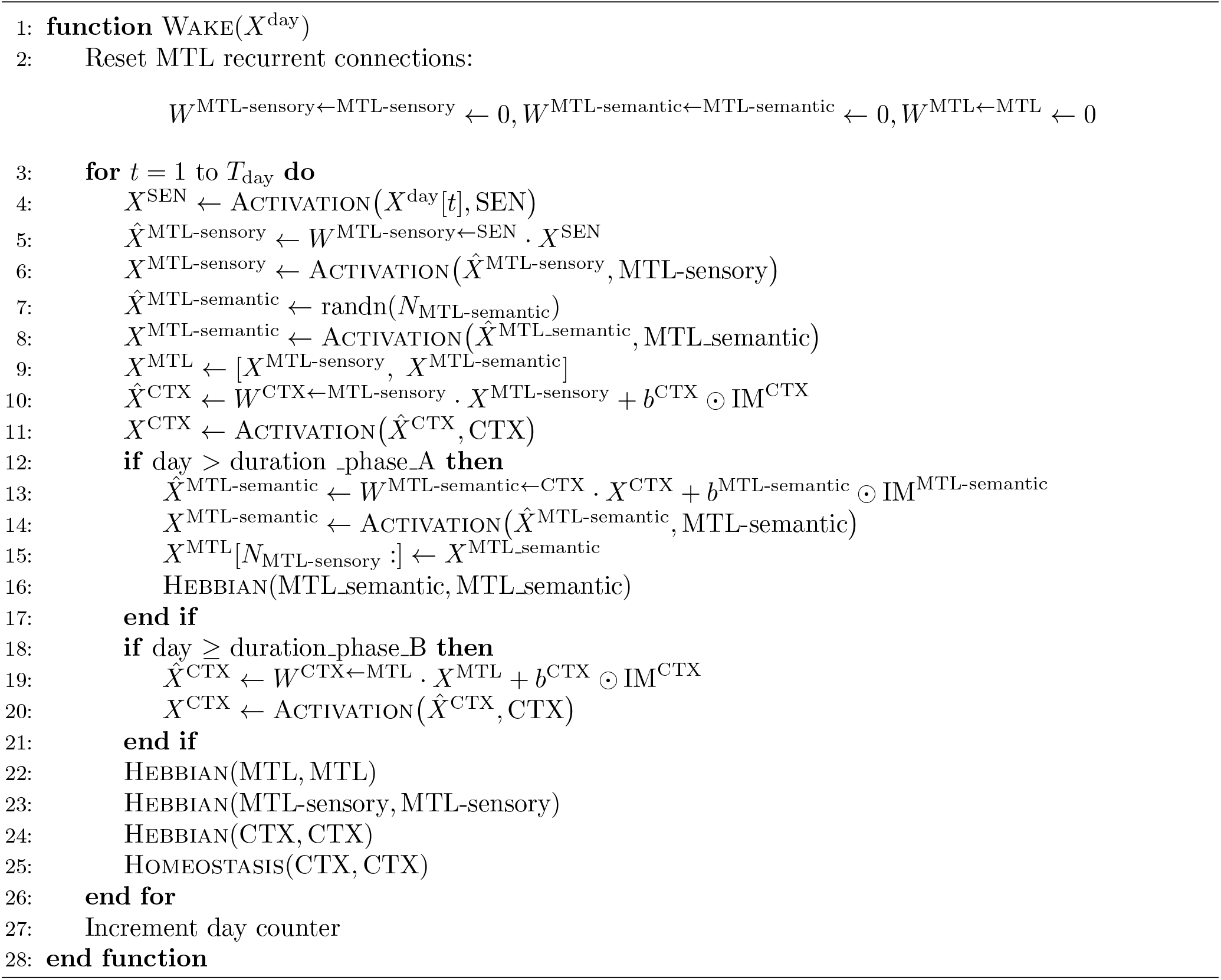

#### Algorithm 8 Sleep

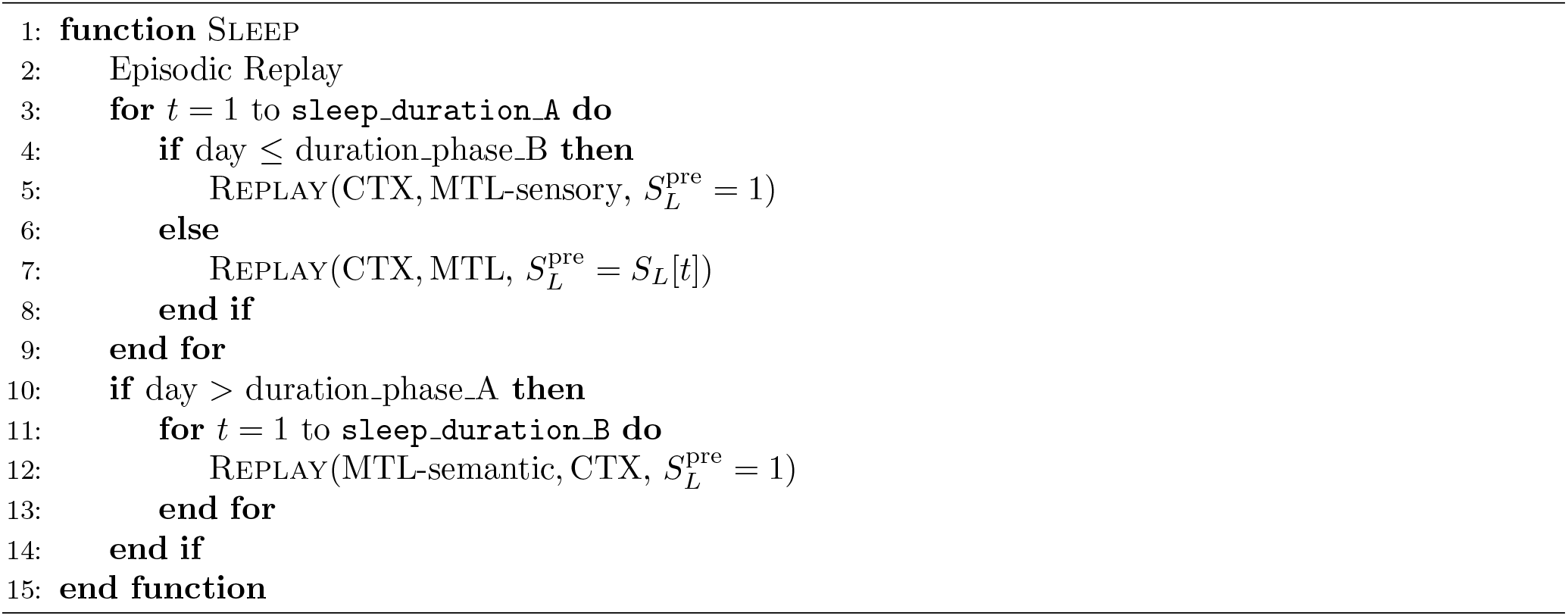

### Considerations regarding modeling choices

The present work has required making several modeling choices, ranging from architectural decisions to model and input generation parameters. In the main results, we have already explored variations of certain critical input parameters, such as noise levels (variations across signal-to-noise ratio) and statistics (variations across episode generation probability distributions). Another relevant parameter studied was the amount of sleep (equivalent to the number of replayed episodes per sleep cycle). However, here we discuss other parameters and modeling choices that have been held at their default values in the main simulations For each, we provide a motivation and/or do additional simulations showing the model performance across parameter sweeps.

#### Input

- **Day length:** Here, all *days* consist of 16 episodes of 5 timesteps each (80 timesteps in total). Robustness to the number of days was studied in simulations (Fig. S9a). As expected, given that consolidation depends on uncovering structure across overlapping patterns, when the total number of presented episodes is very small the selectivity of the formed receptive fields decreases. Behavior becomes relatively consistent for *N*_episodes_ = *T*_day_*/T*_episode_ ≥8, where the chances of episodes sharing conceptual structure increases.
- **Episode duration:** The model remains stable across all tested episode duration values (Fig. S9b).

#### Network model

- **Number of pattern-complete iterations:** The model shows a slight benefit to increased pattern complete iterations when its value is ≤ 3, then stabilizes (Fig. S9c).
- **Sparsity (and number of concepts per latent factor):** Sparsity levels are imposed in the circuit model via the choice of the different *K* values across regions and sleep phases. At the same time, it is also directly related to the number of distinct concepts that can be orthogonally encoded in a group of neurons. We therefore co-vary the network sparsity with the number of distinct concepts within latent factors *A* and *B*, and the network size. For example, a sparsity of 0.1 is associated with the representation of 1+1 out of 10+10 distinct concepts. This in turn means that more cortical neurons are needed to represent a total of 20 simple concepts, and 100 complex concepts. No qualitative change in network selectivity after learning for different sparsity levels was observed (Fig. S9d).
- ***K-winners-take-all* activation function:** Some of the most typical activation functions in neural networks include sigmoidal, step (Heaviside), or rectified linear unit (ReLU) functions. However, these tend to suffer from representational collapse when attractor dynamics deal with highly overlapping patterns such as the ones used here. A top-K activation function, which we use here, can be seen as a generalization of a step function in which the threshold is dynamically adjusted such that the number of active neurons remains stable (or meta-controlled, as in this same study we allow different sparsity levels across learning). In turn, a dynamic threshold can be interpreted as adjustable inhibition with a biological inspiration in adjustable inhibition, which has been observed experimentally (Wehr & Zador, 2003; Vogels et al., 2011). While the particular implementation of the dynamics that maintain sparsity levels stable is abstracted, we speculate one of the functional roles of adjustable inhibition is likely to be the facilitation of learning in the presence of highly correlated overlapping patterns.
- **Neurogenesis, excitability and plasticity rates:** Our dictionary learning mechanism depends on immature neurons being more excitable and plastic than mature neurons. Evidence for this has been found experimentally (Aimone et al., 2011), and has been used in computational models (Gozel & Gerstner, 2021).
- **Homeostatic regimes across connectivity:** In additional simulations (Fig. S9e) we find that, in our model, incoming homeostasis (the sum of incoming synapses is capped) is necessary for selective receptive field formation in feed-forward synapses (as opposed to outgoing homeostasis). Incoming homeostasis is a crucial ingredient for our proposed mechanisms for replay-based dictionary learning, with receptive fields converging to *what is the average pattern that causes active neurons to fire?*. Outgoing homeostasis reverses the map from conditional firing probabilities to connectivity, breaking the learning algorithm. Incoming normalization (akin to L1-norm regularization) is one of the most common choices in synaptic homeostasis, with recent work exploring learning in outgoing homeostatic regimes(Fung & Fukai, 2023), and can be derived from spike-time-dependent synaptic plasticity (Gütig et al., 2003).

### Data Analysis

#### Neuronal Selectivity

Given a matrix *X* that contains neuronal activity across T timesteps -with dimensions (*T, N*_neurons_)-, we define a *concept indicator matrix Z*, a binary matrix that encodes the presence (or absence) of concepts across timesteps. In practice, *Z*(*t*) takes the form

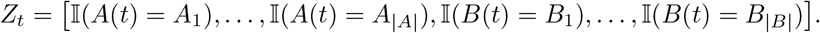

with dimensions (*T*, |*A*| + |*B*|), when selectivity to simple concepts is examined, and

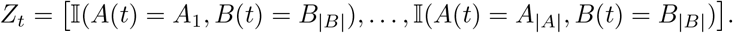

with dimensions (*T*, |*A* || *B*|), when selectivity to complex concepts is examined. To measure selectivity, we obtain the normalized values:

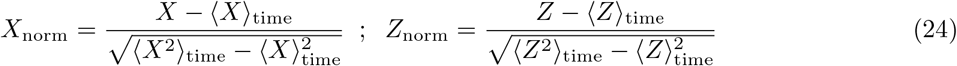

And then obtain the matrix

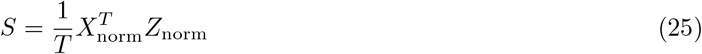

which, for neuron *i* and concept *j* contains their correlation as:

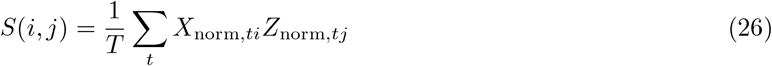

The Maximum Selectivity of neuron *i* is the maximum selectivity value of neuron *i* across all factor-values (concepts) *j*:

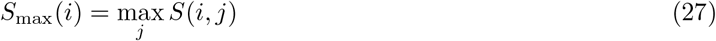

#### Neuron Ordering

Given a selectivity matrix 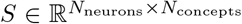, we construct an ordering of neurons into concept-specific assemblies of fixed size *K*_assembly_. The procedure is iterative and greedy. At each round, the concepts encoded across the second dimension are visited in a random order. For each concept in that order, we assign the still-unassigned neuron with the highest selectivity for that concept, and remove that neuron from the global pool of available neurons. This process is repeated until each concept has been assigned exactly *K*_assembly_ neurons. The final ordered index vector is then obtained by concatenating the neurons belonging to each concept-specific assembly. This effectively returns a permutation of a subset of the given neurons, organizing them in blocks of *K*_assembly_ units that code (as much as possible) for the same concept. Note how a natural choice is 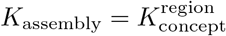.

#### Signal-to-Noise Ratio

In order to obtain the Signal-to-Noise Ratio (SNR) of a generated pattern, we first define the signal *S* as the expected overlap (number of common neurons) of the corrupted pattern with the original, and subtract the expected overlap between two randomly generated patterns of *K*^region^ active neurons across *N* ^region^ neurons:

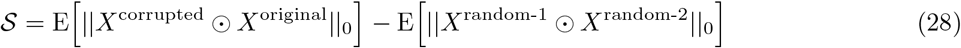

Because we fix that *N*_swaps_ active neurons (in the de-noised pattern) flip from 1 to 0, the first term is always *K*^region^ −*N*_swaps_, while the second corresponds to the expected value of a hypergeometric distribution, given by (*K*^region^)^2^*/N* ^region^. Then, we define the noise to be the number of non-common neurons between the corrupted and the original pattern, which is 2*N*_swaps_, giving:

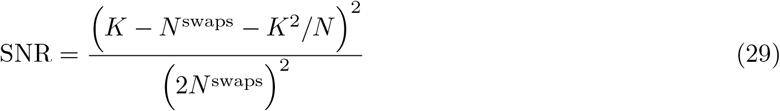

#### Mutual Information

##### Discrete Mutual Information

To quantify how informative neural activity is about latent variables, we use discrete mutual information and entropy estimated from the empirical distribution of awake samples. For two discrete random variables *X* and *Z*, with empirical joint distribution *p*(*x, z*) and marginals *p*(*x*) and *p*(*z*), the mutual information is

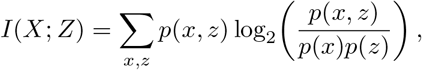

and the entropy of *Z* is

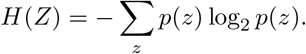

##### Normalized Mutual Information (NMI)

When reporting normalized mutual information (NMI), we divide by the entropy of the latent variable,

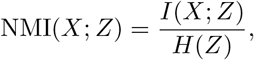

with the convention that NMI = 0 if *H*(*Z*) = 0. This normalization yields values between 0 and 1, where 0 indicates that neural activity carries no information about the latent variable and 1 indicates that it fully determines it.

##### Simple Concepts NMI and Complex Concepts NMI

The latent variable *Z* is always binary and indicates whether a particular concept is present at a given timestep. For *simple concepts* normalized mutual information, we define one binary variable for each simple concept *A*_*i*_ and *B*_*j*_:

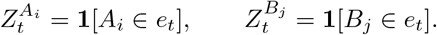

For *complex* normalized mutual information, we instead define one binary variable for each episode-level conjunction *A*_*i*_*B*_*j*_:

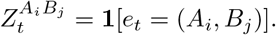

Thus, the distinction between simple concepts and complex concepts NMI lies entirely in which latent variables are used: simple concepts correspond to latent factor values (e.g. *A*_3_), whereas complex concepts correspond to their conjunction (e.g. *A*_3_*B*_2_).

##### NMI Learning Trajectories

To construct the mutual-information learning trajectories (Figs. 2 4), we compute NMI in sliding windows over time. For each latent variable *Z*^*ℓ*^, each neuron activity trace *X*_*n*_ is restricted to a time window, and we compute

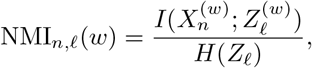

where (*w*) denotes the samples contained in that window, and *H*(*Z*_*ℓ*_) is estimated over the full recording used for that trajectory. Neurons are first grouped into concept-specific assemblies using the selectivity-based ordering described above, and the reported trajectory for latent *ℓ* is obtained by averaging NMI_*n,ℓ*_(*w*) across the neurons assigned to the corresponding assembly. In this way, the trajectories quantify how strongly each learned neural assembly tracks its associated latent variable (concept) as training progresses. Each window *w* contains *T*_NMI_ consecutive timesteps, and successive windows are offset by a stride of Δ*T*_NMI_ timesteps, which is a way of estimating the evolution of mutual information across time, assuming it remains locally stable across small temporal windows.

#### Probability of adding a concept to the cortical vocabulary (forming a receptive field in cortex that is selective to a particular concept)

To quantify how exposure statistics shape cortical dictionary formation (Figs. 2g and 4f), we measure, across many independently simulated networks, whether a targeted simple concept or a targeted abstract episode (complex concept) becomes part of the cortical dictionary after a single session of sleep as a function of its probability of occurrence in the training distribution. In both analyses, each simulation produces a binary formation outcome for focal items, and the plotted quantity is the empirical probability of formation obtained by averaging this binary outcome across all simulations sharing the same focal probability.

For the simple-concept analysis, each simulation defines independent marginal distributions over the two latent factors *A* and *B*. In each dimension separately, one focal concept is chosen uniformly at random, and its marginal probability is sampled from the grid

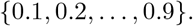

If the focal concept in dimension *A* is 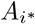, we set

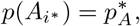

and distribute the remaining mass 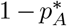 across the other four *A* concepts using a Dirichlet draw. The same construction is performed independently for dimension *B*, yielding

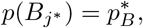

with the remaining mass 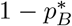 distributed across the other *B* concepts by an independent Dirichlet draw. Episode probabilities are then constructed as a product distribution,

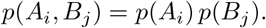

Thus, increasing the focal probability of a concept affects all episodes containing that concept proportionally, while concepts belonging to the same latent factor compete for the remaining marginal mass. By contrast, the opposite latent factor is unaffected except through the multiplicative factor implied by the independent product structure.

For the complex concepts, we evaluate the first day of training with 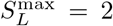 (allowing semantic load 2 during replay). In each simulation, one focal episode (*A*_*i*_, *B*_*j*_) is chosen uniformly from the 25 possible episodes, and its probability of occurrence in training is sampled from a discrete grid

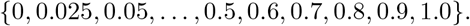

The focal episode is assigned this probability, and the remaining mass is distributed across the other 24 episodes again with Dirichlet sampling.

After training, we evaluate whether the focal concept (simple or complex) has been incorporated into the cortical dictionary. To do so, we extract the receptive fields from CTX, and compare them with de-noised representations of each concept in MTL-sensory. For each concept, we obtain

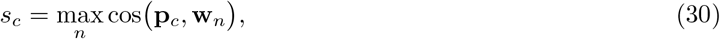

where **p**_*c*_ is the de-noised sensory version of concept *c* and **w**_*n*_ is the receptive field of CTX neuron *n*. The focal concept is considered *added to CTX* when for some receptive field the corresponding de-noised version was the one with highest cosine similarity. Each simulation yields a pair

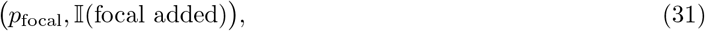

and the plotted curve is obtained by averaging I(focal added) across all simulations with the same focal episode probability.

### Simulations

**Figure 2: Sparse episodic replay creates compositional representations in semantic memory**

To generate Fig. 2, we initialize the network from scratch and train it with the default input and model parameters using *Train* (Algorithm 6). For the example simulation used in Figs. 2b–h, the network first undergoes 100 wake-only warm-up days and is then trained for 1000 days in the Phase A regime, that is, with episodic-to-semantic consolidation only. We first show representative awake activity in MTL-sensory and CTX (Fig. 2b), with CTX neurons ordered using the selectivity-based assembly ordering defined above. We then compare a wake MTL-sensory pattern and a Sleep A replay pattern from the same network (Fig. 2c).

Next, we show the evolution of the feedforward receptive field of one example CTX neuron in *W* ^CTX*←*MTL^ across learning, together with the corresponding synaptic trajectories (Fig. 2d). We then compare cortical responses to the same MTL patterns before and after learning (Fig. 2e): *early* activity is obtained by projecting MTL-sensory patterns through the initial untrained CTX ←MTL weights, whereas *late* activity is taken from the recorded CTX responses of the trained network.

We then quantify the normalized mutual information between neural activity and the simple concepts *A*_*i*_ and *B*_*j*_ in sliding windows over training (Fig. 2f), averaging within selectivity-defined assemblies. We also estimate the probability that a simple concept is incorporated into the cortical dictionary as a function of its marginal probability in the training distribution (Fig. 2g).

Finally, we compare the learned recurrent CTX ←CTX synaptic weights with the conditional-probability structure implied by the episode distribution (Fig. 2h).

**Figure 3: Semantic replay forms abstract representations in episodic memory**

To generate Fig. 3, we initialize the network from scratch and train it with the default input and model parameters using *Train* (Algorithm 6). For the example simulation used in Figs. 3b–f, the network is first trained for 1000 days in the Phase A regime and is then trained for a further 500 days in the Phase B regime, in which semantic replay is enabled. We first show representative awake activity from day 500, before semantic replay begins, in MTL-sensory, MTL-semantic, and CTX (Fig. 3b), with neurons ordered using the selectivity-based assembly ordering defined above. We then initialize CTX from a random state and run cortical pattern completion to show the semantic attractor recovered during semantic replay (Fig. 3c).

Next, we take the same MTL-sensory pattern shown in Fig. 3b and pass it through the fully trained frozen network to visualize the corresponding MTL-semantic response after learning (Fig. 3d). We then compare the maximal selectivity of MTL-sensory and MTL-semantic neurons to the simple concepts *A*_*i*_ and *B*_*j*_ in the trained network (Fig. 3e). Finally, we compute the normalized mutual information between awake MTL activity and the simple factor values (concepts) in sliding windows across the full 1500-day training trajectory, averaging within selectivity-defined assemblies (Fig. 3f).

#### Figure 4: Learning higher-order concepts based on simpler representations is facilitated by the presence of semantic representations during replay

We start from the network trained in the generation of Fig. 3, and continue training under variable replay sparsity using *Train* (Algorithm 6). In the example simulation used for Figs. 4b–e, the network is first trained for 100 additional days with replay semantic load 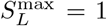, and then for 1000 further days with 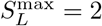 . We first illustrate the replay mechanism by initializing MTL from one random state and comparing the patterns recovered under semantic load 1 and 2 (Fig. 4b), together with the corresponding cortical responses. We show one representative awake snapshot near the end of training (Fig. 4c), highlighting the coexistence of simple-concept and conjunctive cortical representations.

Next, we examine the emergence of complex-concept selectivity in the corresponding cortical subregion by comparing its *early* and *late* responses to the same final awake MTL patterns (Fig. 4d). The early activity is obtained by projecting those MTL patterns through the initial CTX ←MTL weights inherited from Fig. 3, whereas the late activity is taken from the trained network. We then quantify the normalized mutual information between neural activity and the complex latent variables *A*_*i*_*B*_*j*_ in sliding windows across the full higher-order training trajectory (Fig. 4e), comparing MTL-sensory, the simple-concept CTX subregion, and the complex-concept CTX subregion.

Finally, we estimate the probability that a specific episode *A*_*i*_*B*_*j*_ is incorporated into the higher-order cortical dictionary as a function of its marginal probability in the training distribution (Fig. 4f).

**Figure 6A: Semantic representations in episodic memory improve recall capacity**

To generate Figs. 6b–c, we start from the trained network of Fig. S5 and evaluate episodic recall while varying memory load. For the random-pattern baseline (Fig. 6b), we compare three recurrent-memory systems: MTL-sensory alone, the full MTL network, and a sparse Hopfield network as control. For each memory load, we store a set of random fixed-sparsity patterns, probe recall from corrupted cues generated with 4 swaps, and quantify retrieval by the cosine similarity between the recalled and stored patterns, averaging across stored patterns and across 50 seeds.

We then repeat the capacity analysis for structured episodic inputs generated from the standard latent distribution (Fig. 6c). Using the same pretrained network, we compare three modes: *semantics absent, semantics random*, and *semantics present*. For each memory load, we generate one day of experience with one timestep per episode, vary the sensory corruption level across *N*_swaps_ ∈{4, 8, 10 }, run the network on that day, and then probe recall from cues corrupted by 4 swaps. We report recall from MTL-sensory for all three modes and, when semantic representations are available, recall from MTL-semantic as well. To prevent autaptic self-stabilization from inflating high-load recall, recurrent self-connections are set to zero before pattern completion in all capacity analyses. Finally, alongside the recall curves, we plot the cumulative number of distinct concepts encountered as memory load increases.

To provide an associative-memory baseline in Fig. 6b, we implement a sparse Hopfield network that stores *P* binary MTL-sensory patterns 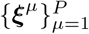 using centred Hebbian weights without self-connections (Eq. 32). During recall, the corrupted cue initializes **h**^(0)^; the network undergoes synchronous threshold dynamics and a final top-*K* projection restores the prescribed activity level, for fair comparison with the output of the MTL network.

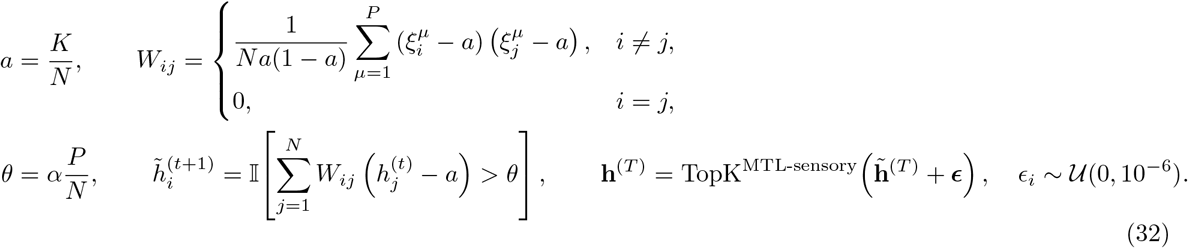

#### Figure 6B: Semantic representations in episodic memory promote compositional generalization during consolidation

In Figs. 6d-g, the same three conditions (*semantics present, semantics absent*, and *semantics random*) are examined in the context of compositional consolidation (complex concepts based on previously learned simple concepts). We pre-train each network with simple concept learning with a relatively low level of noise 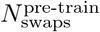 for 600 days, with otherwise default input temporal parameters. Then, we train the network for a further 400 days within complex concept learning, with the maximum replay semantic load set to 2, and with a variable input noise level 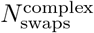.

To quantify generalization of the learned complex concepts representations (*Compositional Generalization* in Fig. 6d), we further test the network over 100 frozen wake days. We use activity in the complex-concept CTX subregion and divide the recorded samples into equal fitting and test partitions. In the fitting partition, we identify ten CTX neurons per complex episode on the basis of selectivity. In the test partition, we average activity within each selected neuronal group and decode episode identity from the most active group (Fig. 6d).

Next, we estimate the SNR of the replayed patterns (Fig. 6e). For samples of replayed MTL-sensory patterns (with semantic load 2), we find its closest noiseless pattern (which de-noised sensory pattern would be more similar to the replayed one?) and infer the effective corruption level of replay by computing the Hamming distance between the evaluated replay pattern and its closest prototype, converting this distance to the equivalent number of input swaps, and then computing the corresponding SNR.

We then estimate the *Replayed/Awake Generalization* of the replayed patterns (Fig. 6f). Intuitively, we want to answer: *how good are the replayed patterns at identifying during wakefulness the concept they are more selective to?* For each network, for each complex concept *A*_*i*_*B*_*j*_, we take 10 random replayed MTL-sensory patterns which, interpreted as receptive fields, are most selective to that concept. This provides a reservoir of receptive fields in which for each complex concept there are 10 examples of receptive fields that could have been proposed by replay. Then, we take the inputs used to test Compositional Generalization and, for every day, we create an episode classifier with one random receptive field from the reservoir for each concept. Then, we use that classifier (class established from the complex concept whose cosine similarity with each input pattern is highest) and obtain an accuracy measure. The average over days (and thus over random classifiers built from replayed MTL-sensory patterns) is the final Replayed / Awake Generalization metric.

Finally, we obtain the *Replayed/Replayed Generalization* (Fig. 6g). This aims at quantifying whether replayed patterns that correspond to the same complex concept consistently recruit the same cortical neurons. We follow a similar process, but instead of measuring how well the receptive fields classify awake input, we measure how well they classify other replayed patterns, where the nature of the episode is again determined by obtaining the cosine similarity with de-noised versions of MTL-sensory patterns that correspond to each episode. Similarly, the accuracy in the classification of the replayed patterns by the random sample of receptive fields establishes the Replayed/Replayed Generalization.

### Reproduction of experimental results

#### Concept cell formation (Quiroga et al., 2005; Quiroga, 2012)

##### Experimental goal

We tested concept-cell formation via different measures aimed at quantifying their main defining properties: a high selectivity to individual concepts that generalizes across contexts and metrics. To do so, we measure (i) cross-context linear decodability, (ii) concept selectivity and (iii) normalized mutual information.

##### Network

We use a pre-trained network (same training regime as Figure 3) and evaluate activity in MTL-sensory and MTL-semantic.

##### Cross-context linear decodability of concept representations

To quantify whether individual concepts are represented in a way that generalizes across episodic contexts, we use a cross-context linear decoding assay. We evaluate activity in MTL-sensory and MTL-semantic under frozen weights across sensory-corruption levels *N*_swaps_ ∈ {1, 2, …, 15}.

For each target concept *c* ∈ *A* ∪ *B*, we construct a binary classification task. If the target is *A*_*i*_ (*c* = *A*_*i*_), we sample one concept *B*_*j*_ to define a *training context* ; if the target is *B*_*j*_, we analogously sample one concept *A*_*i*_. Training data consist of 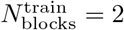 deterministic blocks: one positive episode containing the target concept and one negative episode in which the target concept is replaced by a contrast concept from the same latent factor. Thus, for a target *A*_*i*_ and training context *B*_*j*_, the two training episodes are (*A*_*i*_, *B*_*j*_) and (*A*_*k*_, *B*_*j*_), with *k*≠ *i*. Likewise, for a target *B*_*j*_ and training context *A*_*i*_, the two training episodes are (*A*_*i*_, *B*_*j*_) and (*A*_*i*_, *B*_*ℓ*_), with *ℓ j*. Each block lasts 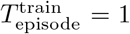 timestep, yielding a minimal balanced decoder-training set by construction.

From this deterministic input sequence, we extract awake activity from MTL-sensory and MTL-semantic, denoted *X*^MTL-sensory^ and *X*^MTL-semantic^. Labels are defined as

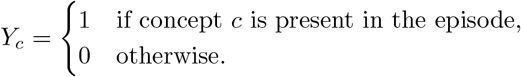

For each region separately, we train a linear logistic classifier to predict *Y*_*c*_ from the corresponding neural activity. In all cases, the same labels are used for both regions, so differences in decoding performance reflect only differences in the underlying neural representations.

To test cross-context generalization, we generate an excluded-context test dataset lasting 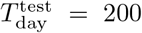 timesteps, with episodes of duration 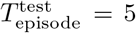 timesteps. If the decoder is trained to recognize *A*_*i*_ in the context of *B*_*j*_, all test episodes containing *B*_*j*_ are excluded. The remaining probability mass is redistributed so that the marginal probability of the target concept remains 0.5, whereas the remaining 0.5 is distributed uniformly across non-target concepts. The same logic is applied symmetrically when decoding a target *B*_*j*_. Thus, the target concept can only be decoded if its representation generalizes beyond the specific opposite-attribute context used for decoder training.

For each target concept, we repeat this procedure for *N*_repeats_ = 10 random choices of training context and contrast concept. Decoder performance is quantified as held-out classification accuracy on the excluded-context test dataset. For each value of *N*_swaps_, the final cross-context linear decodability score for each region is obtained by averaging accuracy across repeats and across all target concepts in *A* ∪*B*. Fig. 5b1 compares the resulting concept-level decoding accuracies for MTL-sensory and MTL-semantic.

For details on *Neuronal Selectivity* and *Mutual Information* measures see above.

#### Consolidation promotes the emergence of representational overlap in the hippocampus and medial prefrontal cortex (Tompary & Davachi, 2017)

##### Experimental goal

We test whether offline consolidation increases the similarity of neural representations for episodes that share a common scene context. As in Tompary and Davachi (2017), overlap is evaluated before and after sleep-induced consolidation.

##### Episode set and latent structure

Each episode contains one object latent factor (*A*) and one scene latent factor (*B*). We use 128 distinct objects and 4 scenes. Objects are assigned evenly to scenes (32 objects per scene), yielding a deterministic training sequence in which each object is presented with its assigned scene.

##### Encoding protocol

Networks are initialized from scratch and trained for one wake day. Each object– scene episode is presented once, with fixed duration (5 timesteps per episode). Trial-level variability is introduced again via *N*_swaps_ bit flipping. This encoding regime produces *T*_episode_ noisy realizations per episode.

##### Pre- and post-consolidation probing

Representational overlap is probed with partial object cues: noisy (*N*_swaps_ = 4) object information is provided, whereas scene input channels are set to zero. For each cue, MTL cue is pattern completed, maintaining the cued neural activity fixed. The recovered MTL state is then propagated to CTX, in turn fed back into MTL-semantic. This is done immediately before sleep (*pre*) and repeated after consolidation (*post* ).

##### Consolidation manipulation

Consolidation consists of one sleep cycle including both replay streams (episodic and semantic replay). Semantic replay is enabled from the beginning of training (i.e., no initial phase with episodic-only replay). Replay durations follow the Figure experiments configuration listed in Table 11.

**Table 11:** Figure 5 Overlap Increase parameters (different than or not specified in default)

| Variable | Description | Value |
| --- | --- | --- |
| $\vert A\vert $ | Number of object concepts | 128 |
| $\vert B\vert $ | Number of scene concepts | 4 |
| $N_{A/B}$ | Objects assigned to each scene | 32 |
| $(N_{\text{SEN}}^A, N_{\text{SEN}}^B)$ | Object and scene sensory-population sizes | (1280, 40) |
| $(K_{\text{SEN}}^A, K_{\text{SEN}}^B)$ | Active sensory units per object and scene concept | (10, 10) |
| $N_{\text{days}}^{\text{encoding}}$ | Number of wake encoding days | 1 |
| $N_{\text{presentations}}^{\text{object}}$ | Presentations of each object during encoding | 1 |
| $T_{\text{episode}}^{\text{encoding}}$ | Duration of each object-scene episode | 5 timesteps |
| $N_{\text{swaps}}^{\text{encoding}}$ | Sensory corruption during encoding | 4 |
| $N_{\text{swaps}}^{\text{cue}}$ | Sensory corruption of partial object cues | 4 |
| $N_{\text{days}}^{\text{phase A}}$ | Duration of the episodic-only replay phase | 0 |
| $N_{\text{sleep}}$ | Number of consolidation sleep cycles | 1 |
| $T_{\text{sleep}}^A$ | Episodic-replay duration per sleep cycle | 50 timesteps |
| $T_{\text{sleep}}^B$ | Semantic-replay duration per sleep cycle | 50 timesteps |

**Table 12:**
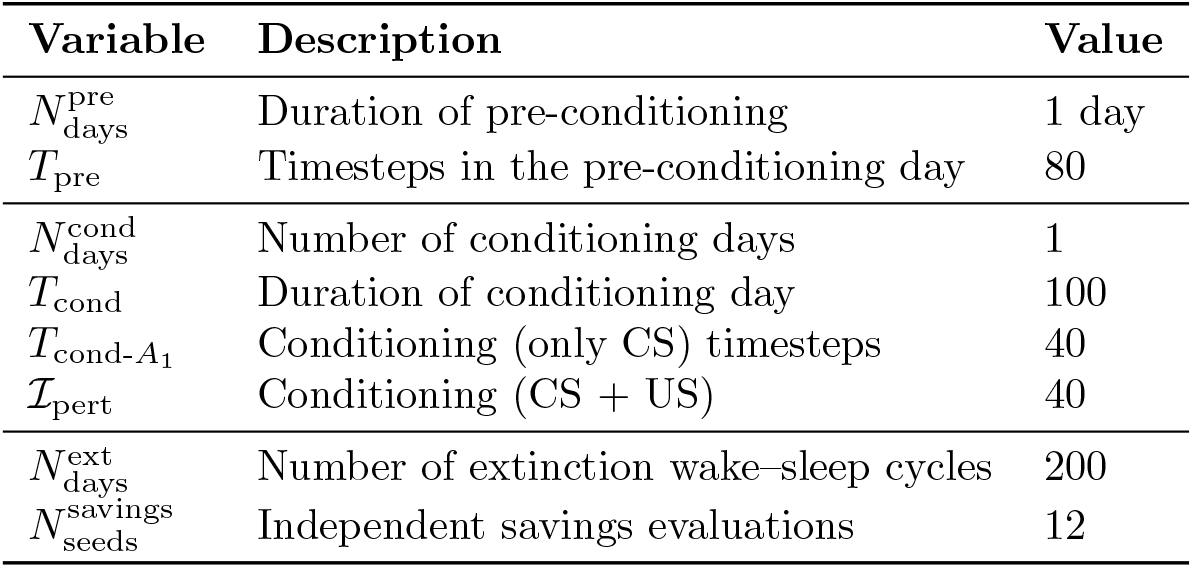
Figure 5 – Neocortical synaptic-engram parameters (different from or not specified in the default model).

**Table 13:** Figure 7- Semantic Facilitation of Recall parameters (different than or not specified in default)

| Variable | Description | Value |
| --- | --- | --- |
| $N_{\text{seeds}}$ | Number of independent experiments | 12 |
| $N_{\text{days}}$ | Number of encoding days per trial | 10 |
| $N_{\text{episodes}}$ | Episodes encoded per trial | 10 |
| $T_{\text{day}}$ | Timesteps per encoding day | 50 |
| $T_{\text{episode}}$ | Timesteps per episode | 5 |
| $N_{\text{swaps}}^{\text{stored}}$ | Sensory corruption of stored episodes | 1, 2, 3, ..., 15 |
| $N_{\text{swaps}}^{\text{cue}}$ | Sensory corruption of recall cues | 4 |
| $N_{\text{trials}}$ | Independent encoding-recall trials | 10 |

**Table 14:** Figure 7 - Blocked Versus Interleaved parameters (different than or not specified in default)

| Variable | Description | Value |
| --- | --- | --- |
| $N_{\text{seeds}}$ | Number of independent experiments | 12 |
| $N_{\text{swaps}}$ | Sensory-corruption levels | $\{1, 2, \dots, 15\}$ |
| Blocked training |  |  |
| $N_{\text{blocks}}$ | Number of successive concept-specific blocks | 10 |
| $N_{\text{days}}^{\text{block}}$ | Training duration per block | 10 days |
| $T_{\text{episode}}^{\text{train}}$ | Timesteps per training episode | 5 |
| Interleaved training |  |  |
| $N_{\text{days}}^{\text{train}}$ | Total training duration | 100 days |
| $T_{\text{episode}}^{\text{train}}$ | Timesteps per training episode | 1 |
| $p(A_i, B_j)$ | Probability of each latent pair | 1/25 |
| Post-training cortical test |  |  |
| $N_{\text{days}}^{\text{probe}}$ | Duration of the frozen evaluation dataset | 200 days |
| $N_{\text{days}}^{\text{recorded}}$ | Final probe days retained for analysis | 100 days |

**Table 15:** Figure 7 - Anterograde Semantic Amnesia parameters (different than or not specified in default)

| Variable | Description | Value |
| --- | --- | --- |
| Pre-training |  |  |
| $N_{\text{days}}^{\text{warmup}}$ | Duration of sparse-replay warmup | 100 days |
| $N_{\text{days}}^{\text{higher-order}}$ | Duration of higher-order training | 1,000 days |
| $N_{\text{swaps}}^{\text{pre}}$ | Sensory corruption during pre-training | 4 |
| $p(A_i, B_i)$ | Probability of diagonal pairs during pre-training | 0 |
| $p(A_i, B_j), i \neq j$ | Probability of each off-diagonal pair during pre-training | 1/20 |
| Novel-association encoding |  |  |
| $N_{\text{days}}^{\text{encoding}}$ | Duration of novel-pair encoding | 1 day |
| $T_{\text{day}}^{\text{encoding}}$ | Timesteps in the encoding day | 15 |
| $N_{\text{swaps}}^{\text{encoding}}$ | Sensory corruption during encoding | 4 |
| Recall evaluation |  |  |
| $N_{\text{seeds}}$ | Number of independent experiments | 96 |
| $N_{\text{swaps}}^{\text{cue}}$ | Sensory corruption levels of the recall cue | $\{1, 2, \dots, 15\}$ |

**Table 16:** Figure 7 - Retrograde Semantic Amnesia parameters (different than or not specified in default)

| Variable | Description | Value |
| --- | --- | --- |
| Recall evaluation |  |  |
| $N_{\text{seeds}}$ | Number of independent experiments | 96 |
| $N_{\text{swaps}}^{\text{CTX}}$ | Cortical cue-corruption levels | $\{1, 2, \dots, 9\}$ |

##### Overlap metric and summary statistics

For each region (MTL, CTX), we compute a pairwise Pearson correlation matrix across all object-evoked probe patterns. For object *i*, we define overlap as:

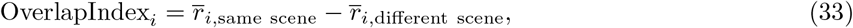

where 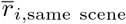 is the mean correlation between object *i* and other objects assigned to the same scene, and 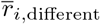 is the corresponding mean across objects assigned to different scenes. Positive values indicate stronger scene-dependent representational overlap.

#### Neocortical synaptic engrams for remote contextual memories (Lee et al., 2023)

##### Experimental goal and circuit mapping

To reproduce synaptic-engram signatures of contextual fear conditioning (Lee et al., 2023), we map prefrontal cortex (PFC) and basolateral amygdala (BLA) populations to CTX and MTL-semantic, respectively. We use two latent factors, context *A* and valence *B*, each comprising five concepts. We interpret *A*_1_ as the conditioned context (CS) and *B*_1_ as the fear-related unconditioned stimulus (US).

##### Pre-conditioning and conditioning

The experiment begins with one pre-conditioning day of *T*_pre_ timesteps. During this phase, *B*_1_ is present in every episode, whereas the activity identities in the *A* sensory channel are independently permuted at each timestep. The network subsequently undergoes its usual sleep consolidation cycle. The intent of this is to mimic the pre-existing fear-selective circuits in the animal. Similarly, this allows us to distinguish cortical *fear* cells, as those that become tuned to the Unconditioned Stimulus, for later behavioral probing.

Conditioning consists of one further day of *T*_cond_ timesteps. Out of these, 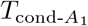 contain only the conditioned stimulus (as happens experimentally), with MTL-sensory cells corresponding to interoceptive signals being random. Then, 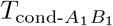 further steps contain the joint presentation, which would correspond to the electric shock. Conditioning is followed by the standard sleep cycle, with semantic replay available from the beginning of the experiment and a maximum replay semantic load of 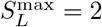.

##### Engram and recall-only populations

We identify engram cells from their mean awake activity during the complete conditioning window. For each region, we compute the temporal mean and apply the region’s standard activation function. The resulting active CTX cells define the PFC engram E_CTX_, while active MTL-semantic cells define the BLA engram E_MTL-sem_.

Recall consists of a further presentation of the same conditioning input, without a subsequent sleep cycle. Normal awake dynamics and plasticity remain active during this pass. Applying the same mean-activity procedure to the recall window yields 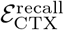, from which we define recall-only cells 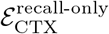 as the recall neurons that are not engram neurons.

##### Extinction and fear-expression proxy

Extinction consists of 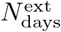 wake–sleep cycles. During extinction, the fear-related US feature *B*_1_ is absent: each of the remaining 20 latent pairs *A*_*i*_*B*_*j*_, with *j*≠ 1, is sampled with probability 1*/*20. Each extinction day contains *T*_ext_ timesteps, is composed of fixed-duration episodes of 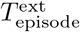 timesteps, and is followed by the standard sleep consolidation cycle.

Before extinction, we identify cortical assemblies selective for the conditioned context *A*_1_ and for the fearrelated feature *B*_1_ using a frozen probe that samples all 25 latent pairs uniformly. After each extinction day and its subsequent sleep cycle, we quantify fear-related cortical coupling as the mean recurrent CTX weight from the *A*_1_-selective assembly to the *B*_1_-selective assembly:

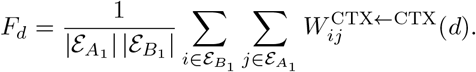

Here, rows *i* correspond to postsynaptic CS-selective neurons and columns *j* to presynaptic US-selective neurons. This measure tracks how extinction reshapes the predictability of fear given the conditioned stimulus. Note how these changes in semantic memory connections effectively translate to *the meaning of the conditioned stimulus being less and less related to that of fear*.

#### Reconditioning and cortical attractor stability

To assess reconditioning after extinction, we compare the post-extinction network with the pre-conditioning checkpoint. For each random seed, independent copies of both networks receive the same *A*_1_*B*_1_ input for *T*_recond_ timesteps. Awake learning remains enabled during this re-exposure, but no subsequent sleep cycle is performed. Thus, the analysis tests the stability of the cortical representation formed after a matched, sleep-free reconditioning period without modifying the stored network checkpoints. For each network state *s*, we define the reconditioned cortical engram as before (CTX activation of mean awake CTX activity). We then measure the memory stability by comparing the cortical pattern completion outcome of each network’s engram with the original state (Fig. 5k). Higher values indicate that the cortical pattern recruited during reconditioning is more stable under that network’s own recurrent dynamics.

#### Semantic facilitation of recall

To test whether semantic interpretability of sensory experience improves recall (Chase & Simon, 1973; Murray et al., 1976; Lin et al., 2021) (Figs. 7 a–b), we compare recall in pre-trained networks with intact sensory input against networks in which semantics are disrupted (or *scrambled* ). This is done by, for each *day*, choosing a fixed input permutation, which transforms neuronal identity of all stored MTL-sensory patterns in the condition *semantics scrambled*. The logic of this manipulation is to isolate the contribution of semantically interpretable input in episodic recall, while maintaining the overall statistics and information load.

We compare a *semantics intact* pretrained network against a *scrambled* version. Test input consists of a single day of length *T*_day_ = 50 timesteps, corresponding to 10 episodes of fixed duration 5. All simulations in this panel are run with a single random seed.

In each trial, the network is first run on this day-long input without sleep, and the resulting MTL activity patterns are stored as target episode representations. Recall is then probed separately for each stored episode by corrupting its MTL-sensory component with cue corruption 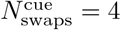, initializing the remaining

MTL units to zero, and iterating recurrent MTL dynamics for 10 steps. Recall performance is quantified as the cosine similarity between the recovered full MTL state and the original stored episode. This procedure is repeated across 10 trials, where each trial consists of a newly generated day-long input sequence followed by recall of all stored episodes.

#### Blocked versus interleaved training

To compare blocked and interleaved training schedules (Fig. 7d), we trained separate networks from scratch while varying only the temporal organization of simple-concept experience. In the *blocked* condition, training was organized into ten successive concept-specific blocks. For each simple concept *A*_*i*_ (*i* = 1, …, 5), the network was trained for 10 days with default temporal parameters, while *A*_*i*_ was held fixed and the paired *B*_*j*_ concepts were sampled uniformly. The same procedure was then repeated for each simple concept *B*_*j*_, holding *B*_*j*_ fixed while sampling *A*_*i*_ uniformly. In the *interleaved* condition, the network was instead trained for 100 days with the same day length, but maximizing variability, with each episode being 1 timestep long, and all 25 combinations equally likely. Thus, blocked and interleaved training matched total wake exposure, differing only in temporal ordering.

After training, we quantified how well simple concepts could be decoded from cortical activity. To do so, each trained network was evaluated on an additional probe dataset of 200 days generated from the default simple-concept distribution used elsewhere in the manuscript. Awake CTX activity from the last 100 probe days was collected, and cortical neurons were ordered by the simple-concept selectivity procedure defined above using the first half of the recorded samples. Accuracy was then measured on the held-out second half using the first half to group neurons according to their selectivity to individual concepts.

### Anterograde lesion of MTL for semantic recall

To model anterograde amnesia (Fig. 7e), we start from a pretrained network that has never seen pairs of the form (*A*_*i*_, *B*_*i*_).

Before learning the new pair, the cortical episode subregion corresponding to previously absent diagonal pairs is reset to an immature state, so that the new association requires fresh cortical recruitment. Two copies of the network are then exposed to the same single-day (*A*_*i*_, *B*_*i*_) association. In the sham condition, the network is allowed to undergo normal sleep-dependent consolidation. In the anterograde condition, MTL recurrent consolidation machinery is lesioned immediately after the awake association day, by setting recurrent MTL connectivity to zero (*W* ^MTL*←*MTL^, *W* ^MTL-sensory*←*MTL-sensory^, and *W* ^MTL-semantic*←*MTL-semantic^).

After this manipulation, semantic recall is tested by showing a corrupted *A*_*i*_*B*_*i*_, and measuring cortical recall (comparing pattern completed cortical pattern with the cortical pattern associated with *A*_*i*_*B*_*i*_).

### Retrograde lesion of MTL for semantic recall

To model retrograde semantic amnesia (Fig. 7f), we start from a pretrained network with default input parameters (output of training in Fig. 4).

Then, we test semantic recall (*B*_*i*_ from matching pair *A*_*i*_) turning on cortical *A*_*i*_ assembly, which is corrupted before pattern completion (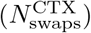). The sham network first does one full feed-forward cycle from CTX to MTL-semantic and back to CTX, then both networks undergo cortical pattern completion. Recall performance is quantified as the mean activity of the target *B*_*i*_ cortical assembly after the *A*_*i*_ cue.

We repeat the analysis over a noise sweep with 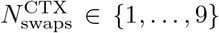 (the number of swaps is smaller because it is over a sparser pattern, which contains less active neurons (only *A*_*i*_)). This analysis isolates the contribution of MTL-semantic feedback to the recall of previously learned associations under degraded cortical cues.

## Acknowledgments

We would like to thank Eleanor Spens, Rachel Swanson, and Colleen J. Gillon, as well as the rest of the members of the Clopath Lab, for discussions and comments on earlier versions of the manuscript. We gratefully acknowledge the financial support of the Wellcome Trust, the Simons Foundation, the Engineering and Physical Sciences Research Council (EPSRC), and the European Research Council (ERC).

## Code Availability

All code supporting the findings of this study is available at https://github.com/albesagonzalez/sensory-semantic-episodes

## Declaration of generative AI and AI-assisted technologies in the manuscript preparation process

During the preparation of this work the authors used ChatGPT in order to assist with writing at the sentence level, and coding at the snippet level. After using this tool/service, the authors reviewed and edited the content as needed and take full responsibility for the content of the published article.

## Supplementary Material

**Figure S1:**
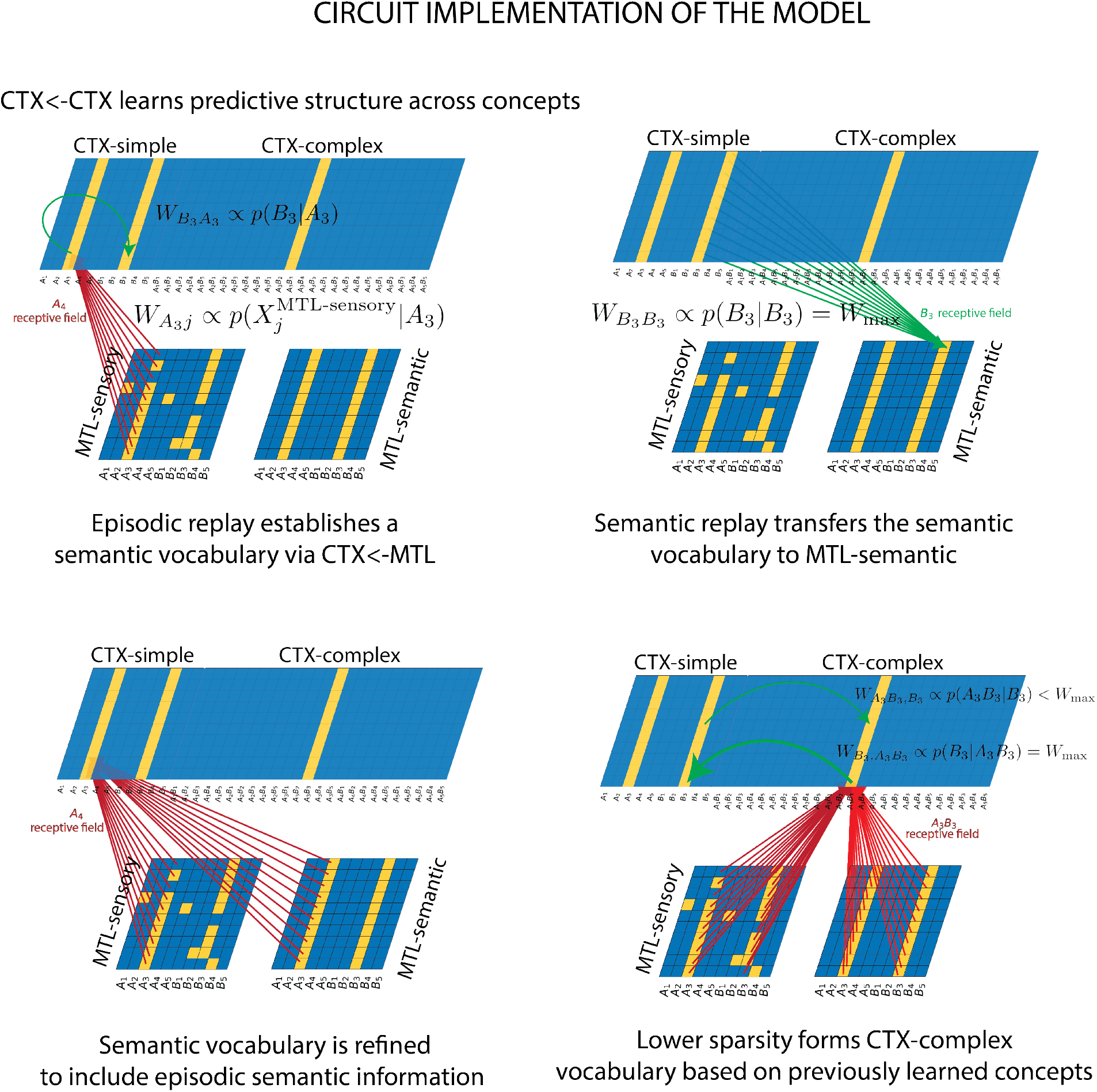
Circuit implementation of the model.

**Figure S2:**
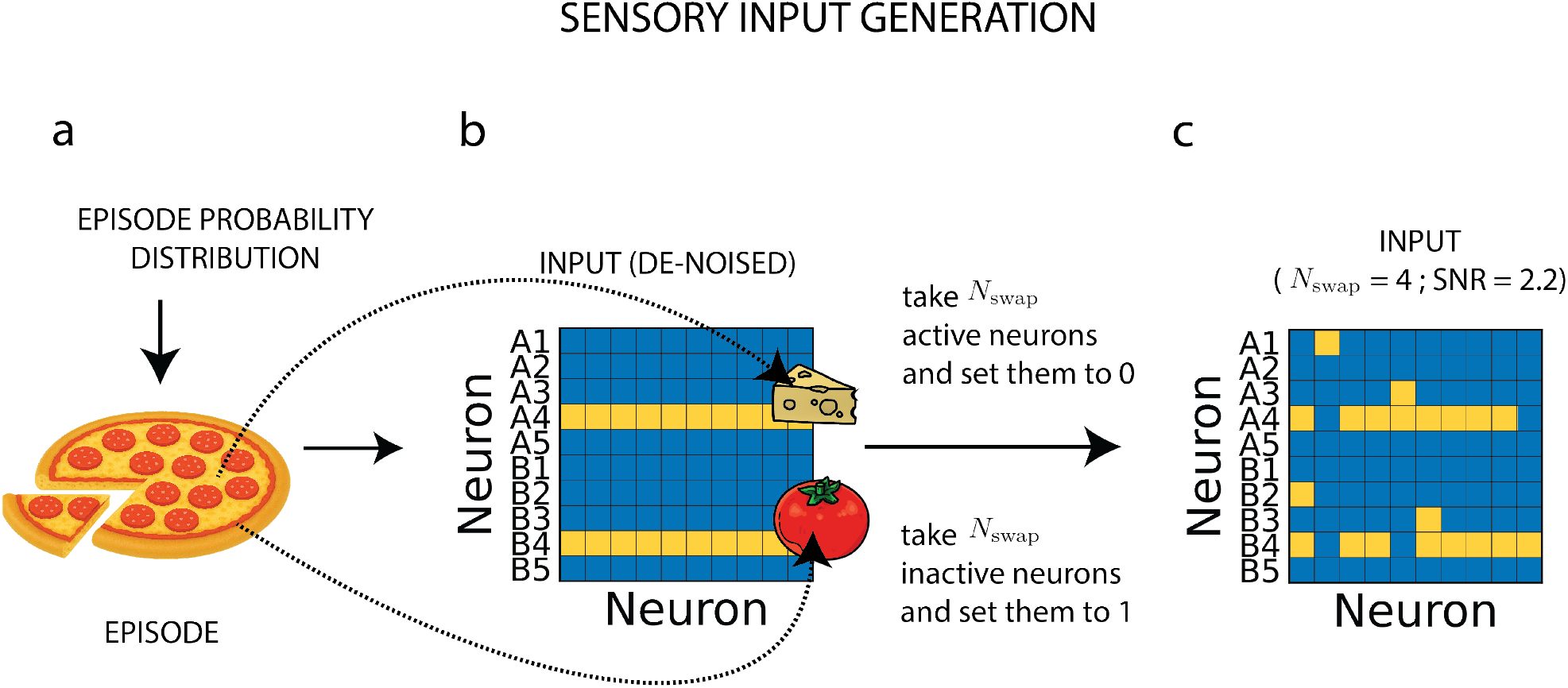
Episode generation protocol. **a**: Episodes are generated by first sampling latent factor values according to the predefined episode probability distribution *p*(*A*_*i*_, *B*_*j*_). Each sampled episode corresponds to a combination of semantic components that define the underlying event. **b**: The sampled factor values (concepts) activate their corresponding sensory features, producing a de-noised input pattern. In the example shown, the episode contains two concepts (illustrated as pepperoni and cheese), each represented by a dedicated subset of active neurons. **c**: Sensory variability is introduced through swap noise. A total of *N*_swaps_ active neurons are randomly deactivated and *N*_swaps_ inactive neurons are randomly activated, generating a noisy sensory realization of the same latent episode. This procedure preserves the underlying semantic content while controlling the input signal-to-noise ratio (SNR).

**Figure S3:**
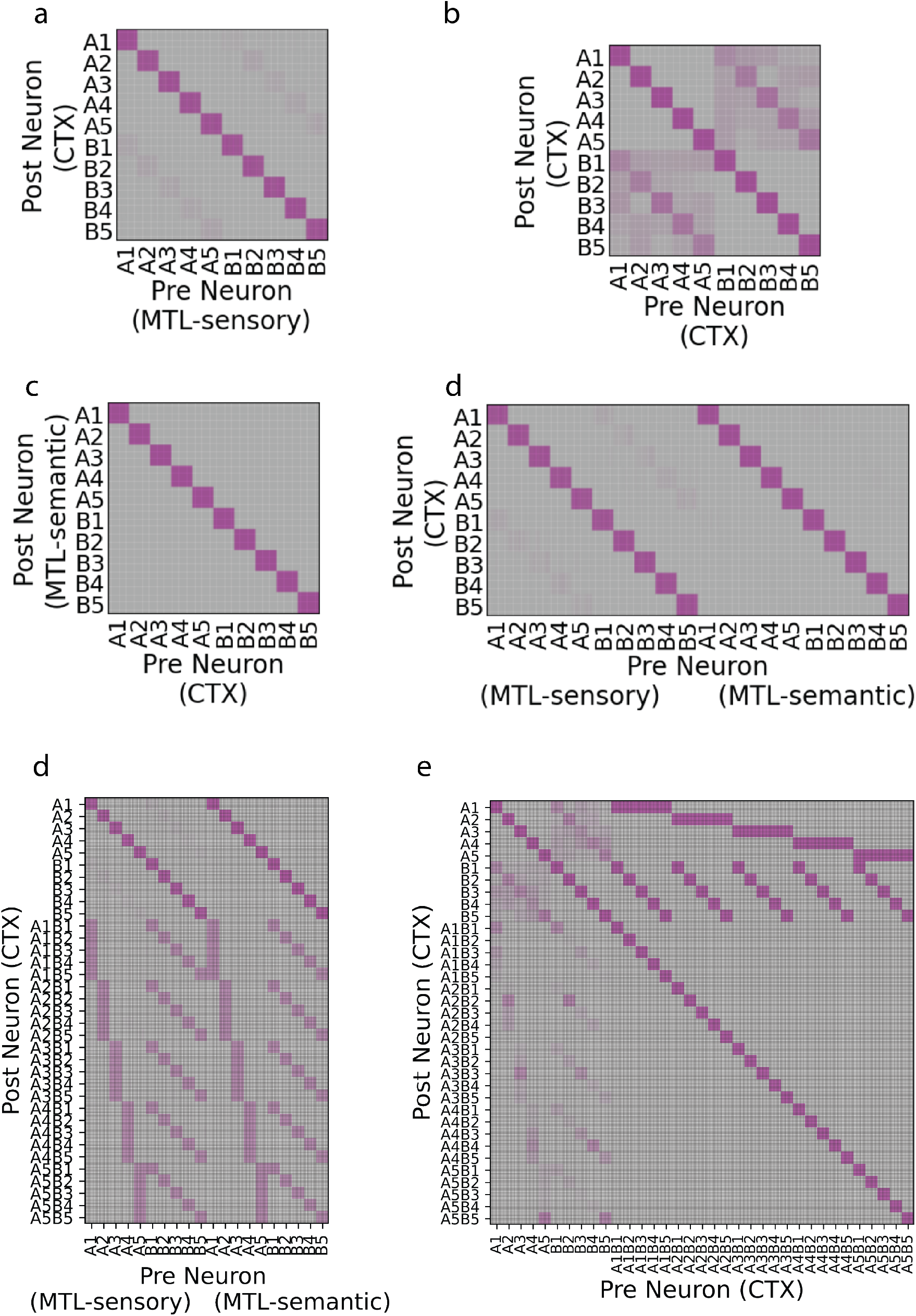
Connectivity matrices across the circuit (after learning)

**Figure S4:**
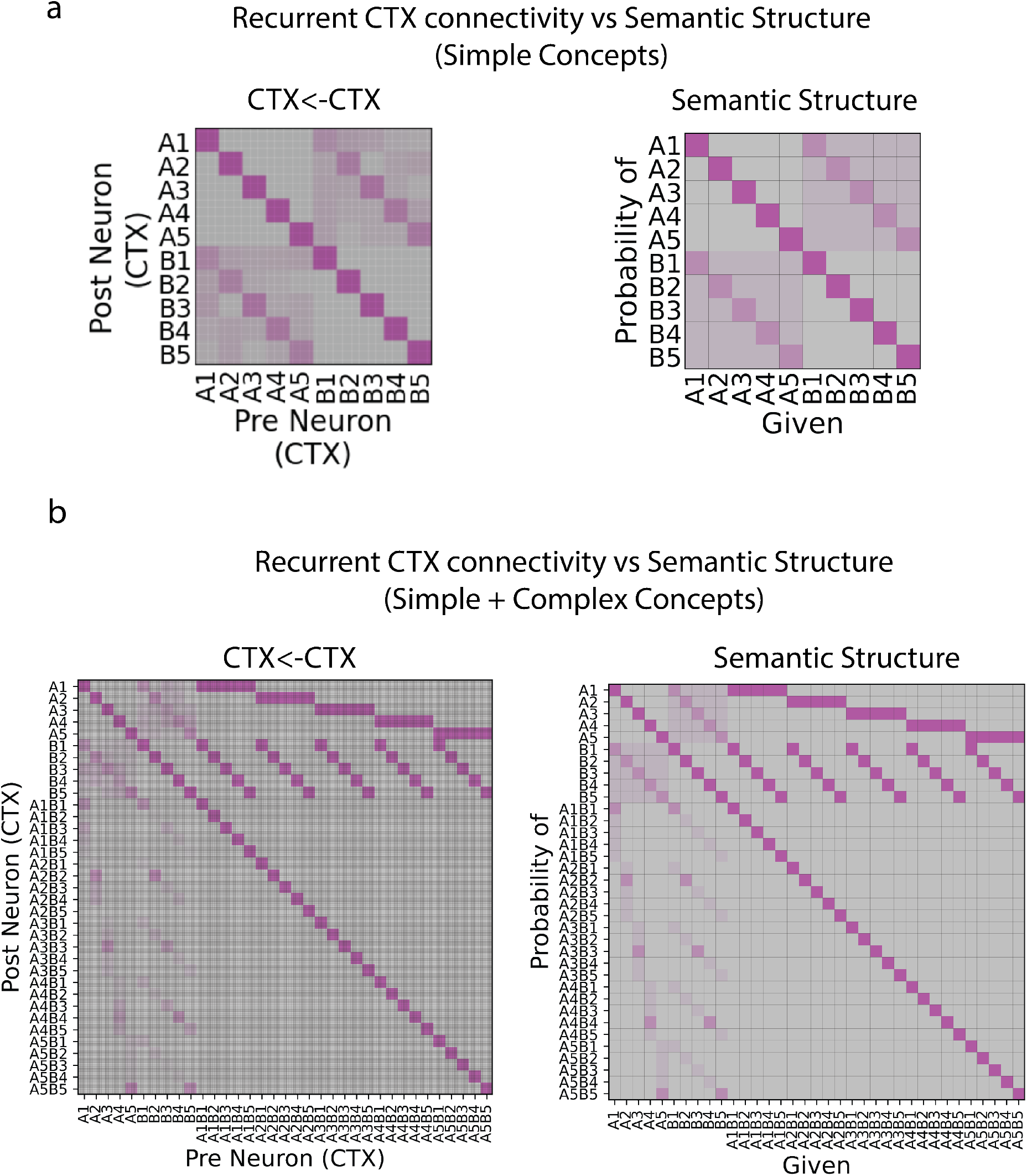
Comparison of semantic structure (model) vs recurrent cortical connectivity (circuit implementation). **a**: Comparison after simple concepts vocabulary has been formed. **b**: Comparison after initial vocabulary has been extended to include complex concepts.

**Figure S5:**
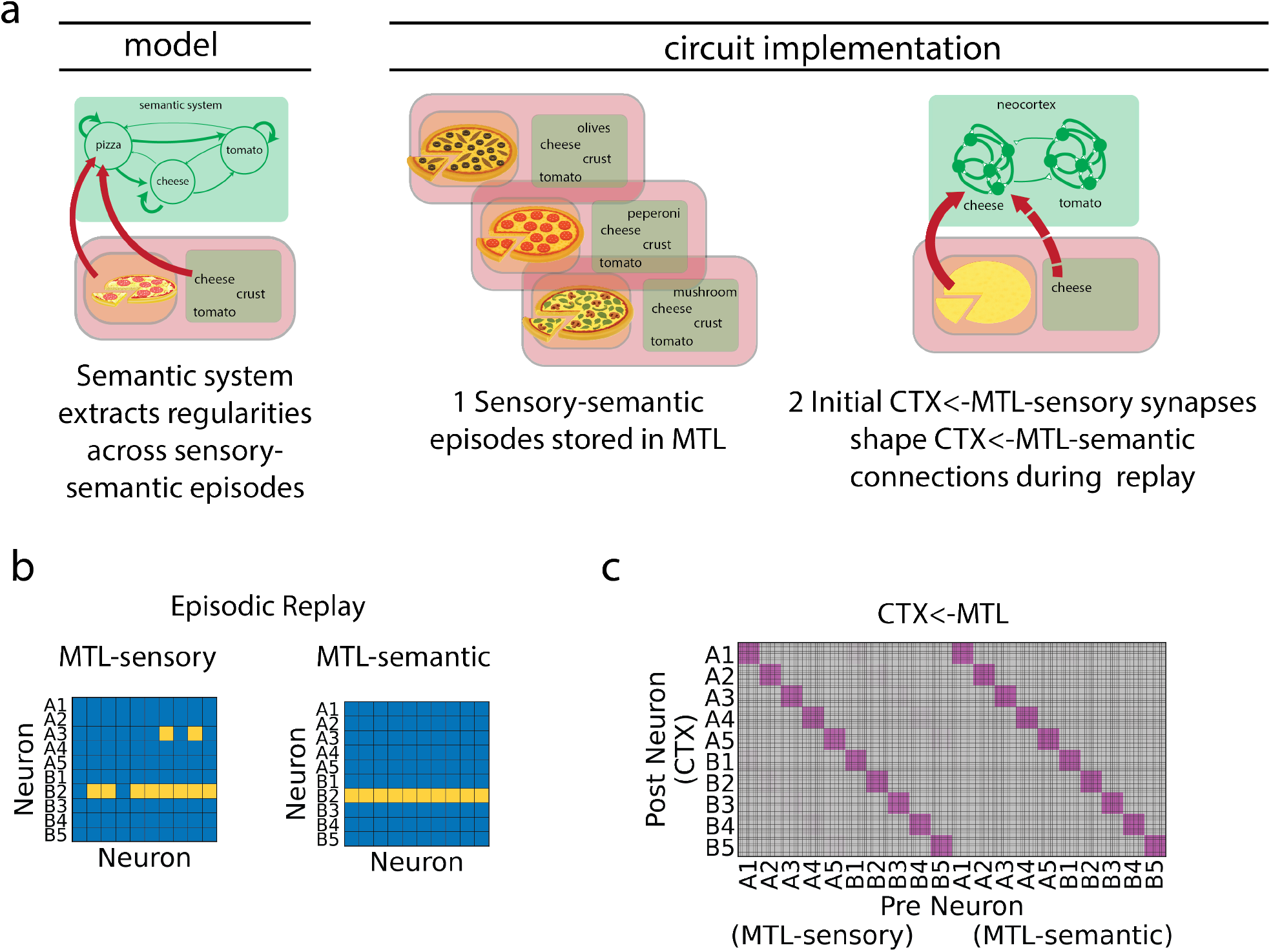
**a**: Schematics of cortical dictionary update after MTL-semantic is developed. Initially, CTX ←MTL-sensory drives CTX activity during episodic replay. Because there is structure in MTL-semantic, this slowly updates CTX ←MTL-semantic connectivity. At learning convergence, this allows semantic indexation from MTL to CTX during episodic replay. **b**: Example MTL activity during episodic replay when replay is driven by full recurrent connectivity (MTL-sensory and MTL-semantic are coupled via reciprocal connections too). **c**: CTX← MTL connectivity after learning with full MTL episodic replay.

**Figure S6:**
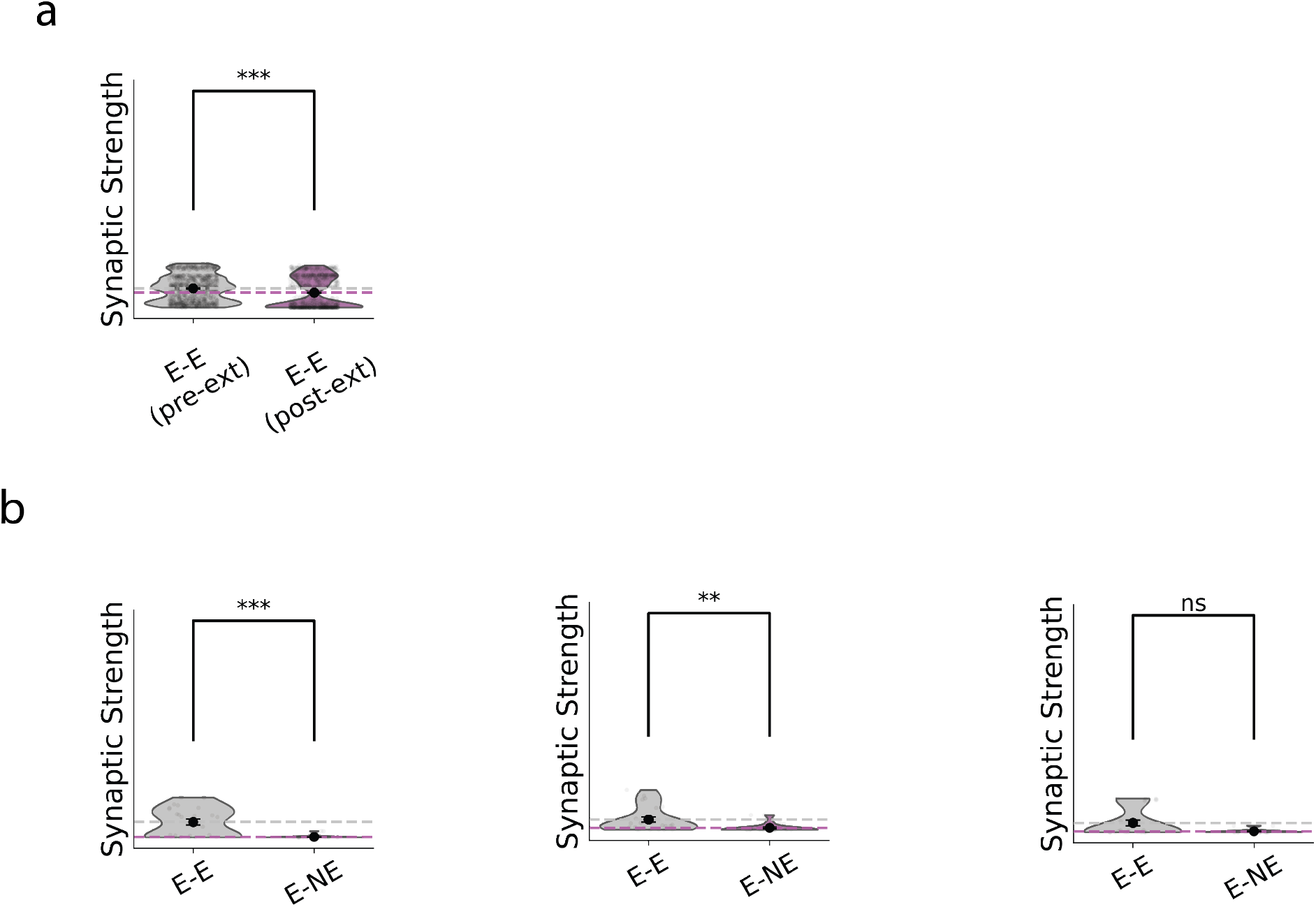
**a**: After extinction, there is a decrease in the mean synaptic coupling from engram to engram neurons. **b**: Statistical significance across E-E and E-NE, after extinction, when matching the number of synapses to the original experiment (5prex5post-synaptic neurons). The test significance is highly dependent on the subsampling random seed.

**Figure S7:**
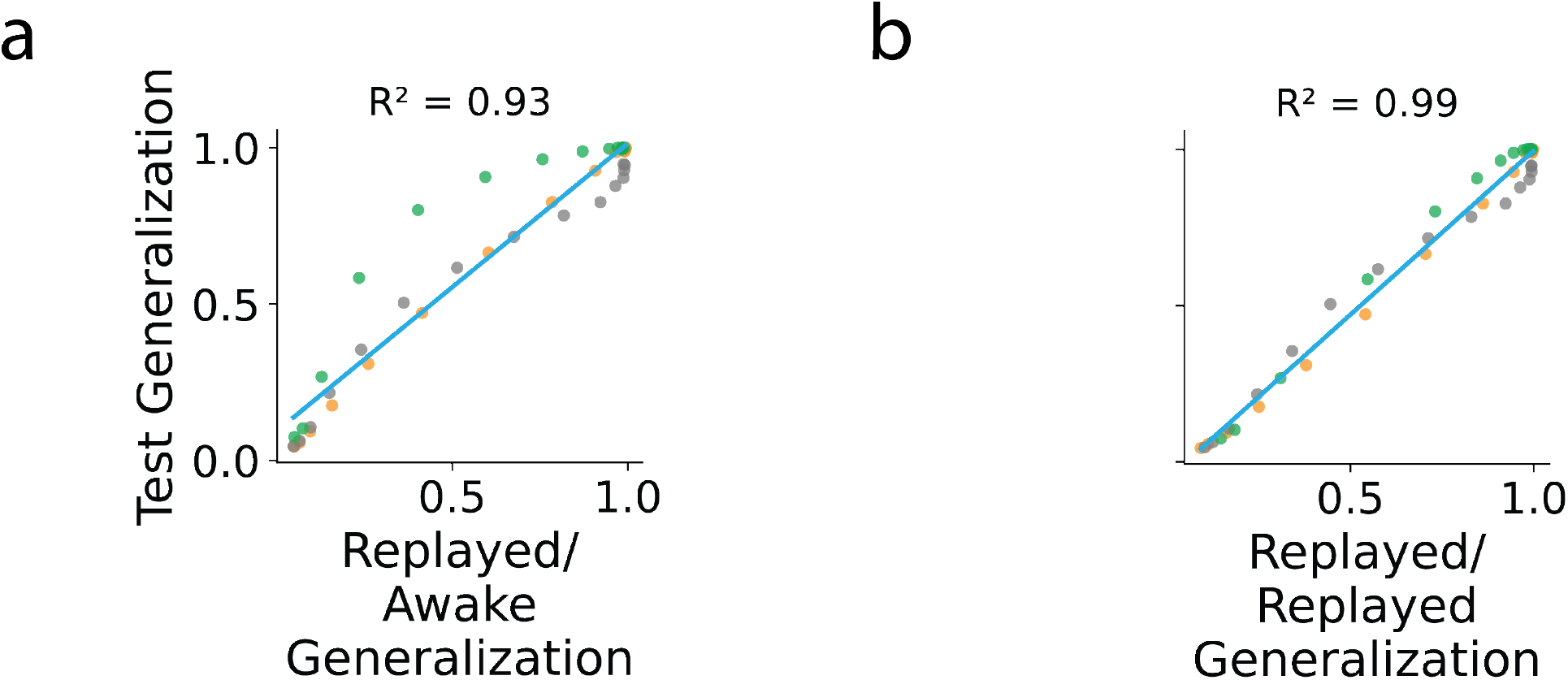
Generalization of replayed patterns and compositional generalization. **a**: Relationship between cortical test-time generalization and replay-to-awake generalization. Each point corresponds to an independent network realization, including different types of networks. Networks in which replayed representations more closely match their corresponding awake representations exhibit higher test-time generalization performance (*R*^2^ = 0.92). **b**: Relationship between cortical test-time generalization and replay-to-replay generalization. Networks in which replay events that correspond to the episode identity (e.g. *A*_1_*B*5) are mapped more similarly onto cortex show improved generalization performance (*R*^2^ = 0.98).

**Figure S8:**
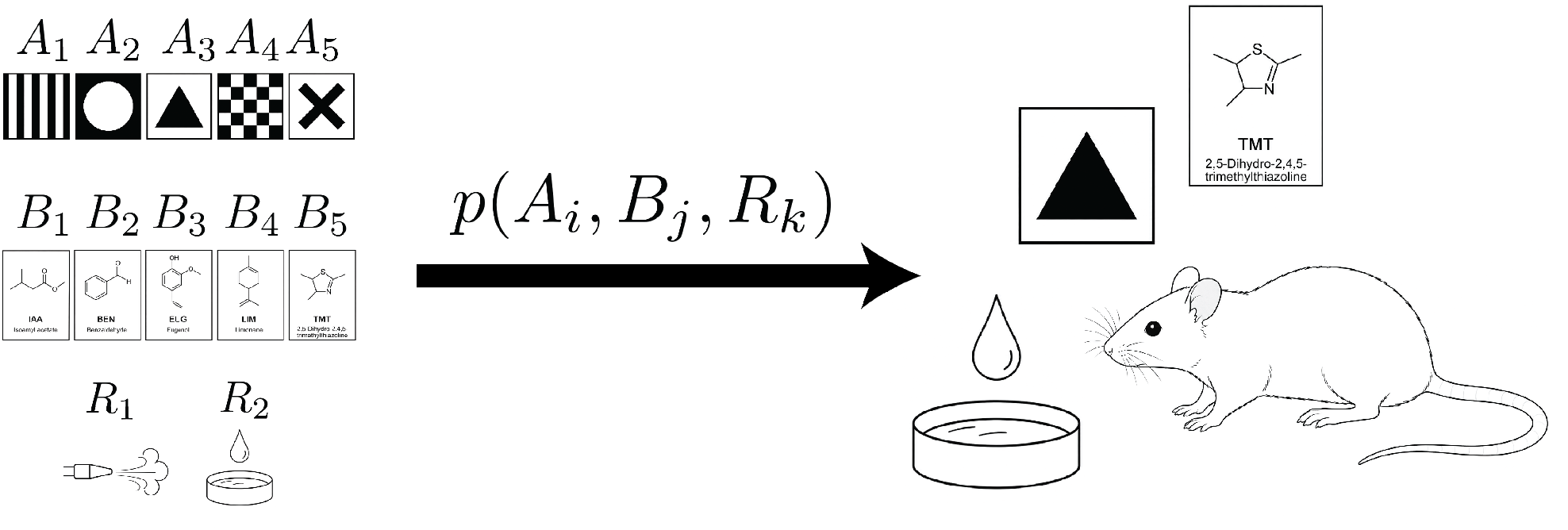
Schematics of an experimental paradigm designed to test the predictions of the model. Compositional visual-odor cues appear following a specific probability on a treadmill, and are associated with either a reward or a negative reinforcement.

**Figure S9:**
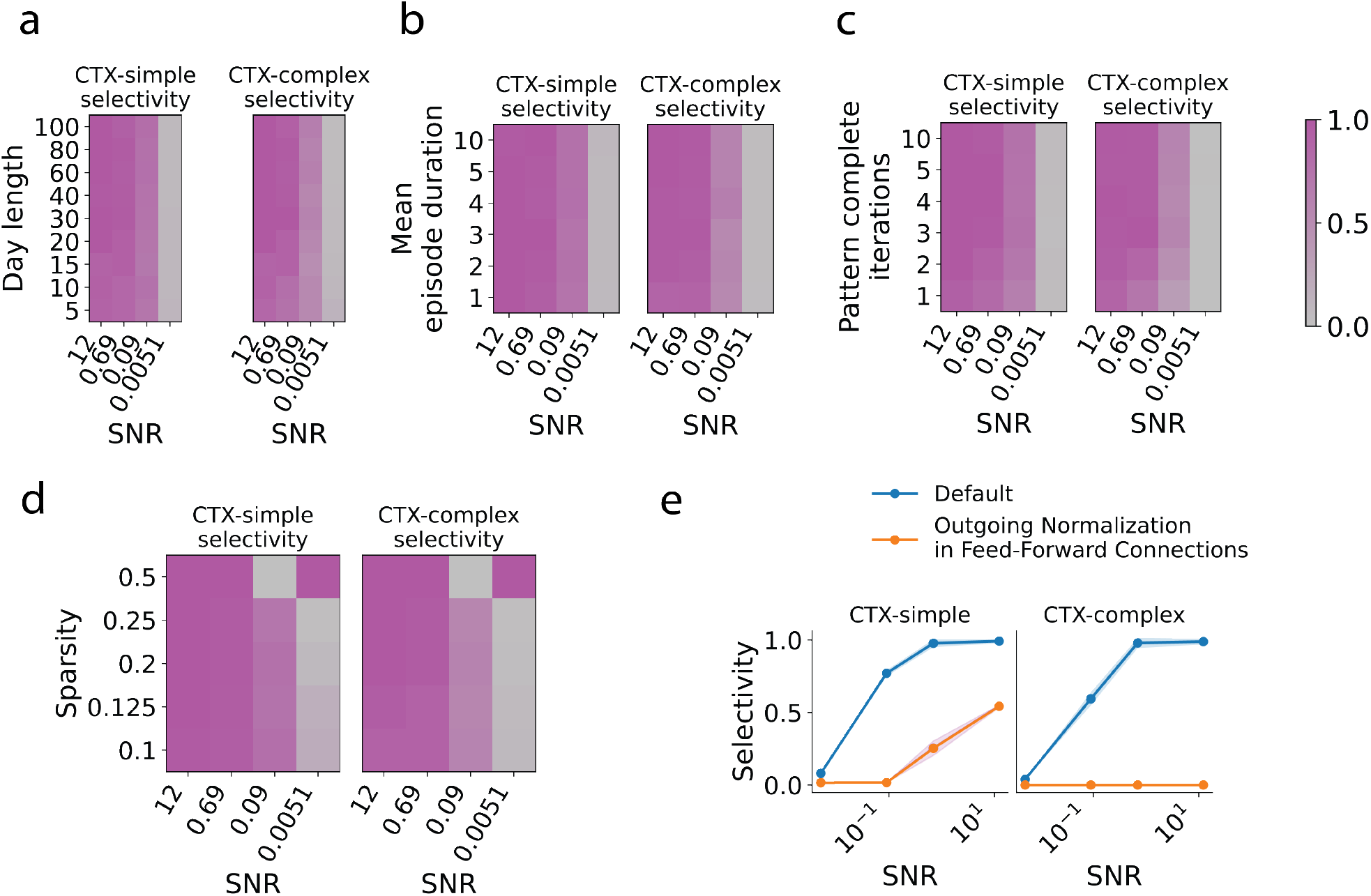
Sweeping across multiple model and input parameters, such as **a**: Day Length *T*_day_, **b**: Episode Duration *T*_episode_, **c**: number of pattern completion iterations in attractor dynamics, and **d**: network sparsity (co-varied with number of distinct concepts per latent factor), shows a high robustness of the model to changes in default parameters. **e**: A dominance of incoming homeostasis in feed-forward synapses is needed in the model to learn selective cortical words.

